# Dissecting tumor transcriptional heterogeneity from single-cell RNA-seq data by generalized binary covariance decomposition

**DOI:** 10.1101/2023.08.15.553436

**Authors:** Yusha Liu, Peter Carbonetto, Jason Willwerscheid, Scott A. Oakes, Kay F. Macleod, Matthew Stephens

## Abstract

Profiling tumors with single-cell RNA sequencing (scRNA-seq) has the potential to identify recurrent patterns of transcription variation related to cancer progression, and produce new therapeutically relevant insights. However, the presence of strong inter-tumor heterogeneity often obscures more subtle patterns that are shared across tumors, some of which may characterize clinically relevant disease subtypes. Here we introduce a new statistical method, generalized binary covariance decomposition (GBCD), to address this problem. We show that GBCD can help decompose transcriptional heterogeneity into interpretable components — including patient-specific, dataset-specific and shared components relevant to disease subtypes — and that, in the presence of strong inter-tumor heterogeneity, it can produce more interpretable results than existing methods. Applied to data from three studies on pancreatic cancer adenocarcinoma (PDAC), GBCD produces a refined characterization of existing tumor subtypes (e.g., classical vs. basal), and identifies a new gene expression program (GEP) that is prognostic of poor survival independent of established prognostic factors such as tumor stage and subtype. The new GEP is enriched for genes involved in a variety of stress responses, and suggests a potentially important role for the integrated stress response in PDAC development and prognosis.

## Introduction

Profiling tumors with single-cell RNA sequencing technologies (scRNA-seq) has the potential to generate new insights into progression and metastasis of cancer as well as therapeutic response [1]. In particular, by analyzing patterns of transcriptional variation, studies can identify “gene expression programs” (GEPs) — i.e., sets of genes whose transcription tends to vary in a coordinated fashion [2–5]. Some GEPs may be patient-specific, whereas others may be shared across (subsets of) patients. Crucially, shared GEPs may characterize different molecular subtypes or cellular states, and so provide insights into disease etiology. Further, associating such GEPs with disease progression or therapeutic response may provide medically relevant insights.

A fundamental challenge to identifying shared GEPs is that tumors often exhibit substantial *inter-tumor* heterogeneity (differences in expression from one tumor or patient to another) [1, 6]. Thus, in scRNA-seq datasets involving multiple tumors, malignant cells typically cluster by patient, with little overlap between patients. This strong inter-tumor heterogeneity tends to obscure the more subtle, but nonetheless important, patterns of heterogeneity shared among tumors. Indeed, as we illustrate in a simulation later, integrative analyses that first combine expression data from malignant cells across all tumors, and then apply off-the-shelf statistical methods (e.g., nonnegative matrix factorization, NMF [7–9]) to identify GEPs, tend to identify patient-specific GEPs, but often miss shared GEPs. Further, the combined analysis of multiple studies — which is desirable due to the typically small number of tumors in individual scRNA-seq studies — introduces potential dataset or batch effects on top of patient effects, further masking shared GEPs.

To address these issues, many recent studies have taken a *tumor-by-tumor* approach in which GEPs that capture *intra-tumor* heterogeneity (differences in expression among cells in a given tumor or patient) are identified separately in each tumor (e.g., by NMF), then compared across tumors to compile a candidate set of shared GEPs [10–16]. However, this strategy often fails to identify some of the more subtle shared GEPs. For example, when tumors are comprised of cells that are homogeneous in terms of molecular subtype identity, the tumor-by-tumor strategy tends to miss GEPs that distinguish among cancer subtypes (see for example the head and neck squamous cell carcinoma case study below). Moreover, this strategy does not benefit from the improved statistical power of an integrated analysis that combines information across tumors.

Another strategy, which tries to combine the benefits of integrated and tumor-by-tumor analyses, is to treat patient effects as “unwanted variation” to be removed [17–19] — a process sometimes known as “harmonization” — and then apply clustering, NMF, or another method for GEP identification to the harmonized data [13, 14, 20–23]. Many harmonization methods have recently been developed for scRNA-seq data [24–29]. While such methods can successfully harmonize datasets from non-diseased tissues [30], they are known to work best when cell types and states are present in comparable proportions across individuals, which is often not the case for tumor data [6]. A recent benchmark study [31] showed that Harmony [24], Conos [28] and Reciprocal PCA (implemented in Seurat 3 [26]) tend to “over-harmonize” cancer datasets, removing genuine variation between patients and thus failing to separate patients with different glioma types.

Therefore, there is an urgent need for improved statistical methods that can better dissect transcriptional heterogeneity in scRNA-seq data into both patient/dataset-specific and shared GEPs. Here we introduce such a method, based on a new approach to matrix factorization that we call “generalized binary covariance decomposition” (GBCD). We illustrate with simulations how GBCD addresses limitations of existing approaches and yields more interpretable results in the presence of strong inter-tumor heterogeneity. To demonstrate its potential to yield new biological insights we apply GBCD to scRNA-seq data from a study on head and neck squamous cell carcinoma (HNSCC), and to three studies on pancreatic cancer adenocarcinoma (PDAC). In HNSCC our approach captures subtype-related GEPs contributing to shared patterns of inter-tumor heterogeneity which were missed by a tumor-by-tumor NMF analysis [10]. In PDAC our analyses provide a refined characterization of previously-identified clinically relevant subtypes, and identify a novel GEP related to stress response pathways that is prognostic of poor survival even when accounting for established prognostic factors such as tumor stage and subtype.

## Results

### Methods overview

We assume that the combined scRNA-seq data from all tumors are contained in an *N ×J* matrix ***Y*** of expression values with entries *y*_*ij*_, where *i* = 1, …, *N* indexes cells and *j* = 1, …, *J* indexes genes. In typical applications, ***Y*** contains log-transformed pseudo-count-modified UMI counts (“log-pc counts” for brevity; see Section S.1.1.1 for details). Our method is “unsupervised” in that, unlike tumor-by-tumor and many harmonization approaches, it does not use information about which cell comes from which tumor.

Our method ultimately yields a decomposition of the data matrix ***Y*** into matrices ***L*** and ***F*** such that ***Y*** *≈* ***LF*** ^*T*^, or equivalently

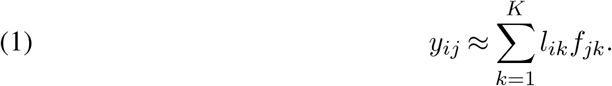

Our goal is that the *K* components in (1) should be interpretable as GEPs, with *l*_*ik*_ representing the membership of cell *i* in GEP *k*, and *f*_*jk*_ representing the effect of GEP *k* on the expression of gene *j*. When *y*_*ij*_ are log-pc counts, each *f*_*jk*_ approximately represents the *log-fold change* (LFC) of gene *j* in a cell, *i*, with membership *l*_*ik*_ = 1 relative to a cell *i*^*′*^ with membership *l*_*i′k*_ = 0. We refer to the *f*_*jk*_ values as LFCs, and we call ***f***_*k*_ := (*f*_1*k*_, …, *f*_*Jk*_)^*T*^ the “signature” of GEP *k*.

Throughout, we adopt “GEP” as a term of art for the components in (1), but we emphasize that some components may not have a coherent biological function; for example, some components may capture technical effects, or effects on gene expression induced by copy number variation. Further, since our method does not know which cells come from which tumors, any patient-specific or batch effects would also appear as “GEPs”. That is, our matrix decomposition takes an agnostic, unsupervised, approach to discovering components of transcriptional variation, which may be both technical and biological; determining which components are most biologically relevant is done by subsequent analyses of the gene signatures (see below).

We make two key assumptions about the matrix decomposition (1):

#### Assumption 1.

Memberships *l*_*ik*_ are nonnegative, and often nearly binary (close to 0 or 1).

#### Assumption 2.

The GEP signatures ***f***_*k*_ are mutually orthogonal.

Assumption 1 is motivated by our particular interest in capturing discrete substructures such as patient effects and tumor subtypes, where membership among cells can be modeled as binary. However, since non-discrete structures may also be of biological interest, and can naturally arise from spatial or continuous cell processes, we implement this assumption as a “soft” assumption; that is, our modeling approach encourages, but does not require, binary membership values. We refer to this assumption as the *generalized binary assumption*.

Assumption 2 is motivated by our empirical observation that, without this assumption, a shared GEP can get absorbed into multiple patient-specific effects (namely, the effects corresponding to the patients who share that GEP). The orthogonality assumption helps to preserve the shared GEP because absorbing it into multiple patient-specific GEPs would make the signatures of the patient-specific effects dependent, and so non-orthogonal. The MNN method of [27] also makes an orthogonality assumption, and indeed we see in the simulation study that this helps MNN separate shared GEPs from patient effects. Similar assumptions have also been suggested in computer vision as a way to force matrix decomposition methods to learn factors that capture shared image features [32–34].

Note that (1) allows for modeling *overlapping structures* in the cell population, in that multiple nonzero *l*_*ik*_’s are permitted for each cell *i*. For example, a given cell might have membership in four GEPs — a dataset-specific GEP, a patient-specific GEP, a tumor subtype GEP, and a cellular state GEP — and this configuration would be represented by four nonzero *l*_*ik*_’s. This ability to model overlapping structures is a key feature that distinguishes our approach from clustering methods, which identify non-overlapping structures.

Combining (1) with Assumption 2 implies the following:

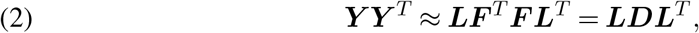

where ***D*** := ***F*** ^*T*^ ***F*** is a diagonal matrix. In practice, our approach proceeds by first finding ***L*** and ***D*** in (2), and then estimating ***F*** based on (1) (at which point we do not force the orthogonality assumption, so in this sense Assumption 2 is also a “soft” assumption.) The matrix ***Y Y*** ^*T*^ is closely related to the cell-by-cell covariance matrix, and so we refer to the decomposition (2) as *generalized binary covariance decomposition* (GBCD), and for brevity here we also use GBCD to refer to our overall approach. To implement GBCD we apply the empirical Bayes matrix factorization (EBMF) method [35] implemented in the R package `flashier` [36–38]. Conveniently, EBMF provides an automatic way to select the number of components *K* (Section S.1.3).

We assess the biological relevance of inferred GEPs by examining the top “driving genes” — that is, genes *j* in GEP *k* with largest magnitude of *f*_*jk*_ — and we perform gene set enrichment analyses [39] to identify biological processes enriched for the driving genes. While driving genes can be either up-regulated (positive *f*_*jk*_) or down-regulated (negative *f*_*jk*_), in our analyses we focus on up-regulated genes. This facilitates comparisons with previously derived expression programs which have usually focused on up-regulated genes.

### Illustrative example

To illustrate the difference between GBCD and existing integrative strategies — including NMF methods, which are commonly used to identify structure in scRNA-seq data — we use a simple simulation designed to mimic patient-specific effects and different types of shared gene expression programs. Specifically we simulate scRNA-seq counts for *J* = 10, 000 genes in *N* = 3, 200 cells from 8 patients (400 cells per patient), with three different types of GEPs (Fig. 1A):

**Fig. 1.**
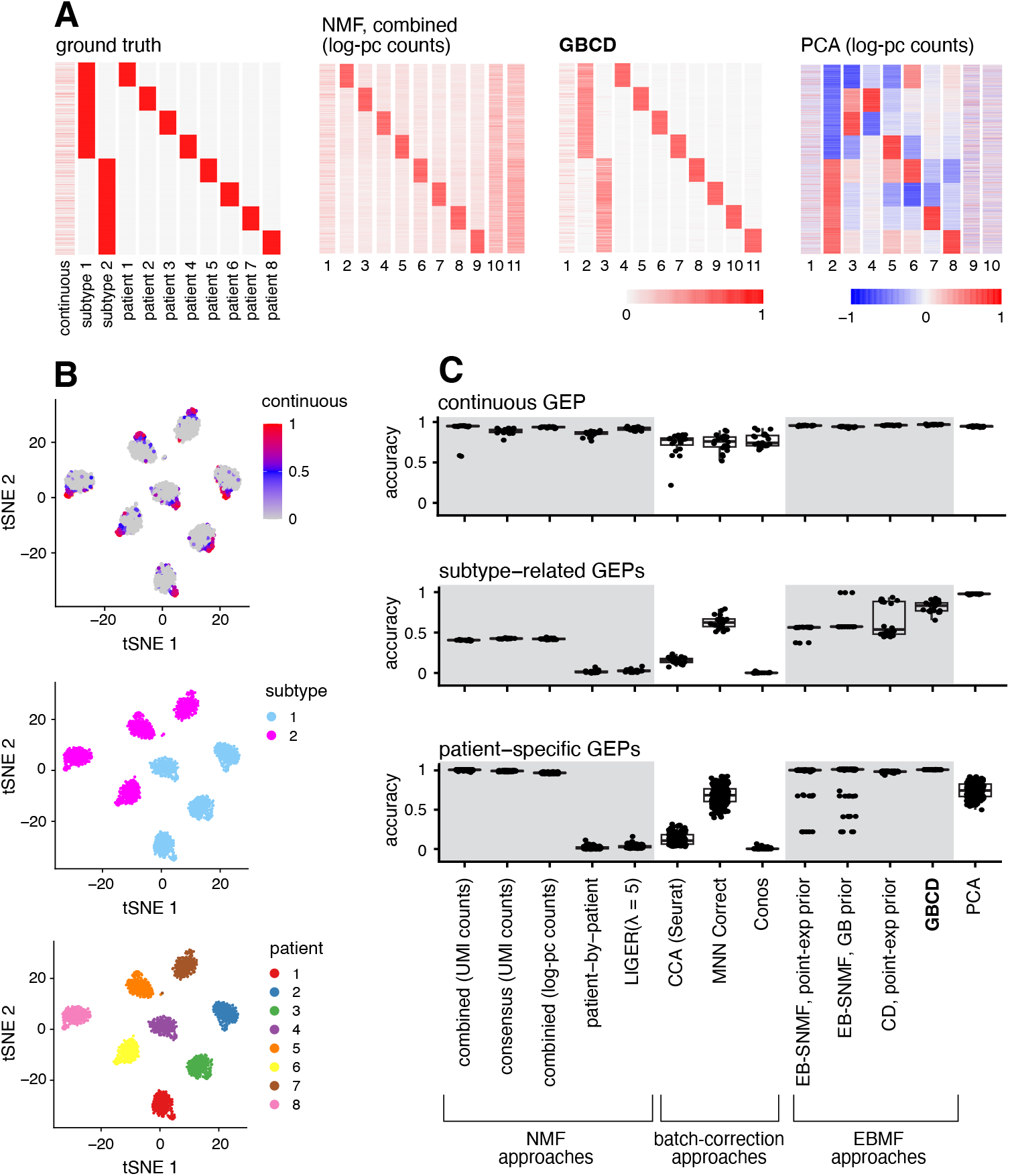
Results of applying different methods to identify GEPs from simulated multi-patient scRNA-seq data. Panel A: the 3,200 *×* 11 membership matrix ***L*** used to simulate the data, and estimates of ***L*** obtained by applying NMF, PCA and GBCD to one of the 20 simulated datasets. Rows are cells, columns are GEPs. In these heatmaps, the columns were scaled separately so that the largest value in each column was 1. All methods identified one or more components that were strongly correlated with cellular detection rate [40]; these components were not included in the heatmaps. See Supplementary Fig. S1 for membership estimates produced by the other methods compared. Panel B: *t*-SNE visualizations of the same scRNA-seq data, in which cells are colored by membership to the continuous GEP, subtype GEPs, and patient-of-origin GEPs. Panel C: Performance of all methods in recovering the continuous, subtype-related and patient-specific GEPs in 20 simulated datasets. See Online Methods for a detailed description of the methods compared. For each true GEP, accuracy of a method’s estimate was measured using the highest Pearson correlation between the true GEP membership and the estimated membership among all GEPs identified by the method.

- Eight patient-specific GEPs: GEPs P1 to P8 are each uniquely expressed by a single patient, with binary memberships. These GEPs mimic the strong patient effects that are present in single-cell tumor data.
- Two subtype-related GEPs: GEPs S1 and S2 are respectively expressed in patients 1–4 and 5–8 with binary memberships. These GEPs mimic cancer subtype programs shared by multiple patients.
- One continuous GEP: this GEP is expressed in 600 cells randomly chosen from the 3,200 cells independently of patient identity, with non-zero membership values drawn from Unif(0.4, 2). This GEP mimics some cellular process or activity whose expression intensity shows continuous variation across cells and may be unrelated to cancer subtype.

In short, the continuous GEP contributes to shared patterns of *intra-tumor* heterogeneity across patients whereas the subtype-related GEPs do not. Further, the subtype-related GEPs induce *hierarchical* relationships between patients, in the sense that patients 1–4 are transcriptionally more closely related to each other than patients 5–8; however, we note that the simulated data do not conform to a strict hierarchy due to the continuous GEP whose expression levels in cells are independent of patient or subtype identity.

Each of the 11 GEPs has a set of differentially expressed genes unique to that program, resulting in GEP signatures that are orthogonal to each other (satisfying Assumption 2). In order not to unfairly disadvantage the NMF-based approaches, the differentially expressed genes for each GEP are up-regulated relative to some baseline expression level by design, so that the true GEP signatures are non-negative. More details on data generation are provided in Section S.1.2.

We independently simulated 20 scRNA-seq datasets under this scenario. Results for a typical simulation are shown in Fig. 1A, Supplementary Fig. S1 and S2. In this dataset, the strong simulated patient effects — that is, the large expression differences between patients — are reflected in the *t*-SNE embedding [41], in which the primary structure is the clustering by patient (Fig. 1B).

We first compared GBCD with NMF applied to combined expression data across all patients (“combined NMF”). GBCD and both versions of combined NMF (applied to UMI or log-pc counts) successfully identified patient-specific components (Fig. 1A, Supplementary Fig. S1), with GBCD providing corresponding membership estimates that were visually the least noisy. All three methods also effectively found the continuous GEP (Supplementary Fig. S2). Notably, GBCD was able to denoise the membership estimates of patient-specific components without compromising the estimates of the non-zero membership values of the continuous component, highlighting the flexibility of the generalized binary assumption.

The primary difference among the three methods was that GBCD accurately identified the two subtype-related GEPs, whereas the combined NMF methods did not (Fig. 1A, Supplementary Fig. S1). This difference is traceable to the orthogonality assumption of GBCD; without an orthogonality constraint, the combined NMF methods can produce a solution where the subtype-related GEP signatures are absorbed into the patient-specific GEP signatures. That is, instead of estimating the 8 patient-specific components to have the “true” orthogonal gene signatures ***f***_P1_, …, ***f***_P8_, the combined NMF methods estimated the patient-specific gene signatures as ***f***_P1_ + ***f***_S1_, ***f***_P2_ + ***f***_S1_, ***f***_P3_ + ***f***_S1_, ***f***_P4_ + ***f***_S1_, ***f***_P5_ + ***f***_S2_, ***f***_P6_ + ***f***_S2_, ***f***_P7_ + ***f***_S2_, ***f***_P8_ + ***f***_S2_. This decomposition of the expression data ***Y*** is less desirable, because it does not separate the shared components of expression from the patient-specific components, thus failing to recover the largely hierarchical relationships between patients. Since this solution does not not satisfy the orthogonality assumption (e.g., ***f***_P1_ + ***f***_S1_ and ***f***_P2_ + ***f***_S1_ are not orthogonal to each other because they share ***f***_S1_), it is ruled out by GBCD, which is thus encouraged to separate out the subtype-related GEPs.

To confirm that the differences shown above were systematic, rather than the chance outcome of a single simulation, we summarized the performance of these approaches to recover different types of GEP across the 20 replicate simulations. The results (Fig. 1C) confirmed that both GBCD and combined NMF consistently detected the patient-specific and shared continuous GEPs, whereas GBCD more reliably recovered the shared subtype-related GEPs.

We next compared GBCD with principal component analysis (PCA) applied to combined log-pc counts, another widely used approach for dimension reduction of scRNA-seq data. PCA successfully identified both the subtype-related and the continuous GEP, but it failed to identify the patient-specific GEPs (Fig. 1A). In place of patient-specific GEPs, PCA identified several principal components (PCs) that were loaded on cells belonging to a seemingly random subset of patients. This is problematic for interpretation because these PCs appear, falsely, to capture features in the data that are shared across patients. Part of the problem with PCA is that it constrains the PCs ***l***_*k*_ to be orthogonal to each other, which is not satisfied when hierarchical relationships exist between patients (e.g., ***l***_S1_ and ***l***_P1_ are not orthogonal to each other). Analysis of results across all the 20 replicate simulations confirmed that PCA had substantially worse specificity for GEP discovery than GBCD, in the sense that a much larger proportion of the GEPs estimated by PCA (i.e., PCs) did not correlate with any of the 11 true GEPs (Supplementary Fig. S3).

Other strategies similarly failed to identify the shared subtype-related GEPs. (Details of these methods and their implementations are given in Sections S.1.3 and S.1.4.) Consensus NMF [5], which averages the solutions of combined NMF over multiple runs, failed to identify subtype effects because NMF could not detect such effects in the first place. Patient-by-patient NMF, which fits NMF to log-pc counts separately for each patient and then clusters the GEPs detected, failed to identify subtype effects because such effects do not vary across cells within a patient thus do not contribute to intra-tumor heterogeneity. A variation on our GBCD method, which replaces Assumption 1 with a generic sparsity assumption (point exponential priors), identified some subtype effects, but not as reliably as GBCD, and did not provide as clear a separation between cells that were active and inactive in each GEP having binary memberships. This finding demonstrates the importance of Assumption 1 in identifying discrete structures. Other variants of empirical Bayes matrix factorization that drop Assumption 2 tend, like NMF, to miss subtype-related GEPs. “Batch effect correction”-type methods that integrate scRNA-seq data from multiple samples, including LIGER [25], CCA implemented in Seurat [26], MNN Correct [27] and Conos [28], likewise failed to identify subtype effects, and also occasionally produced much worse membership estimates of the continuous GEP, consistent with [31] which observed a tendency of these methods to over-correct cancer data. Among the batch-correction methods, MNN Correct was most successful in identifying components related to the subtype-related GEPs, and MNN Correct also makes a orthogonality assumption.

This simple example demonstrates that GBCD can produce qualitatively different results from NMF and other strategies, and highlights the potential of GBCD to identify GEPs that are shared across patients but do not necessarily lead to intra-tumor heterogeneity, which other methods may miss. However, we note that the representations obtained by existing methods here are not “incorrect”. For example, the combined NMF solution is essentially equivalent to clustering cells by patient, which in some contexts could be the desired goal. In practical applications it may be helpful to run multiple methods to obtain different views of the structure present in the data, and GBCD may be seen as complementing existing analysis methods, rather than replacing them.

### Head and neck squamous cell carcinoma data

To compare GBCD and NMF-based methods on real cancer data, we analyzed scRNA-seq data collected by Puram et al. [10] from primary tumors from 10 HNSCC patients and matching lymph node (LN) metastases from 5 of these patients. Puram et al. found that each of these 10 patients clearly mapped to a molecular subtype of HNSCC, whose signatures were previously defined by analysis of bulk expression data from 279 HNSCC tumors [42]. Here we demonstrate that GBCD can extract this molecular subtype information *de novo* from the single cell data alone, which other methods struggle to do.

We analyzed the *N* = 2,176 malignant cells, whose integration among tumors presents a greater challenge than non-malignant cells. As noted by Puram et al. [10], these cells demonstrate strong patient effects that are typical of cancer data; the major structure in the *t*-SNE visualization is the clustering of the cells by patient (Fig. 2A). Because of these strong patient effects, Puram et al. took a tumor-by-tumor analysis approach: they applied NMF separately to each tumor to identify GEPs, and then used clustering to identify similar GEPs from different tumors and create consensus GEPs that they called “meta-programs”. Their analysis identified 6 meta-programs associated with cell cycle, stress, hypoxia, epithelial differentiation, and epithelial-mesenchymal transition or EMT (referred to by [10] as “partial EMT” because it demonstrated some features of classical EMT but lacked others). However, this tumor-by-tumor analysis of the single cell data failed to identify GEPs that classify the patients into their molecular subtypes. This illustrates a limitation of tumor-by-tumor analysis, which can only identify GEPs whose memberships vary sufficiently across cells within each tumor.

**Fig. 2.**
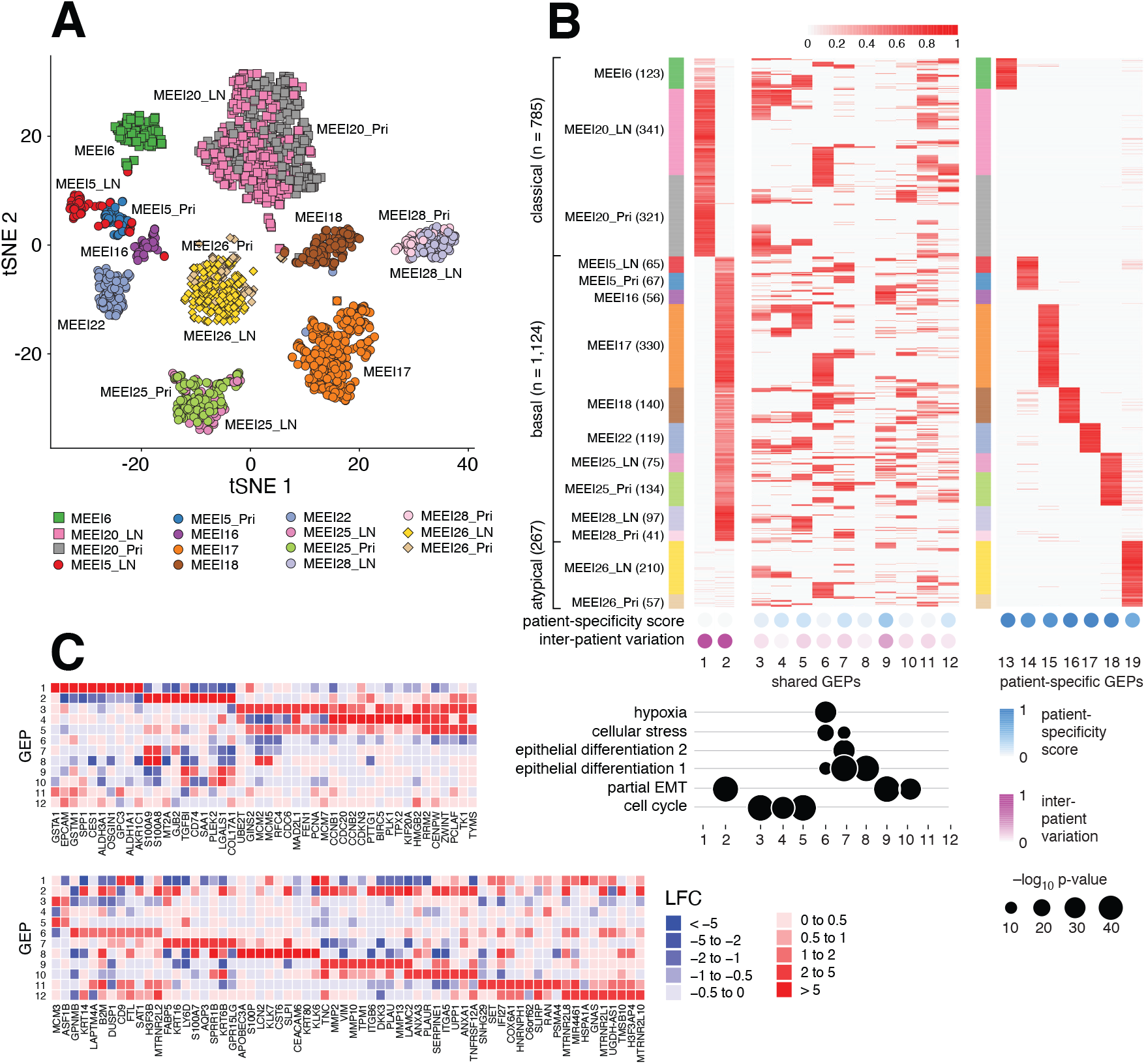
GBCD analysis of HNSCC data. Panel A: *t*-SNE embedding from *N =* 2,176 scRNA-seq profiles in 10 primary tumors and 5 matched lymph node (LN) metastases. Cells are colored by patient of origin and tumor stage (primary tumor, LN metastasis), and shaped by tumor molecular subtype (square: classical; circle: basal; diamond: atypical). Panel B: Heatmap showing membership values of the *N* = 2,176 cells (rows) for the 19 GEPs (columns) identified by GBCD, in which cells are arranged top-to-bottom by tumor molecular subtype and patient, and GEPs are grouped left-to-right based on whether they are more shared across patients (GEP 1–12) or more patient-specific (GEP 13–19). For the heatmap, membership values were rescaled separately for each GEP so that the maximum membership for each GEP was always 1. Numbers in parentheses give the number of cells per tumor. Measures of patient-specific expression (relative to shared expression across patients) and inter-patient variation in expression (relative to intra-patient variation) are shown. See Section S.1.8 for definitions. Concordance between the gene signatures of shared GEPs and 6 previously identified meta-programs from a tumor-by-tumor analysis [10] is quantified by *−* log_10_(*p*-value), where *p*-values are produced from one-sided Wilcoxon rank-sum tests (Online Methods). Panel C: LFC estimates for selected genes in the shared GEPs.

We applied GBCD to these data, estimating GEP memberships and signatures, and annotating the GEPs by their driving genes and gene set enrichment analysis, or GSEA (Fig. 2B, Table 1, Supplementary Fig. S18, Supplementary Tables S1–S2, Section S.1.6). In total, GBCD identified 19 GEPs (not including an “intercept” GEP on which all cells are loaded). Among the 19 GEPs, 12 were active in multiple patients and the remaining 7 were specific to an individual patient (Section S.1.8). Among the 12 shared GEPs, at least one GEP showed significant overlap with each of the 6 meta-programs from [10]. However, some of the shared GEPs were not strongly correlated with any of the 6 meta-programs; for example, GEP11 (enriched for genes involved in respiratory electron transport) and GEP12 (mRNA splicing). Most importantly, GBCD identified GEPs corresponding closely to the previously defined molecular subtypes: GEP1 was largely active only in cells from the 2 classical patients, and GEP2 was mainly active in cells from the 7 basal patients. Therefore, GEPs 1 and 2 were sufficient to accurately classify the 10 patients into the 3 subtypes. We emphasize that this analysis was entirely unsupervised; GBCD does not use the patient or subtype labels. Analysis of variance (ANOVA) of GEP memberships (Section S.1.8) for each of the 12 shared GEPs revealed that GEPs 1 and 2 showed much greater inter-patient than intra-patient variation whereas the opposite was true for the other shared GEPs (Fig. 2B). This is consistent with the observation that tumor cells from an individual HNSCC patient are largely homogeneous in terms of molecular subtype identity [10].

**Table 1.**
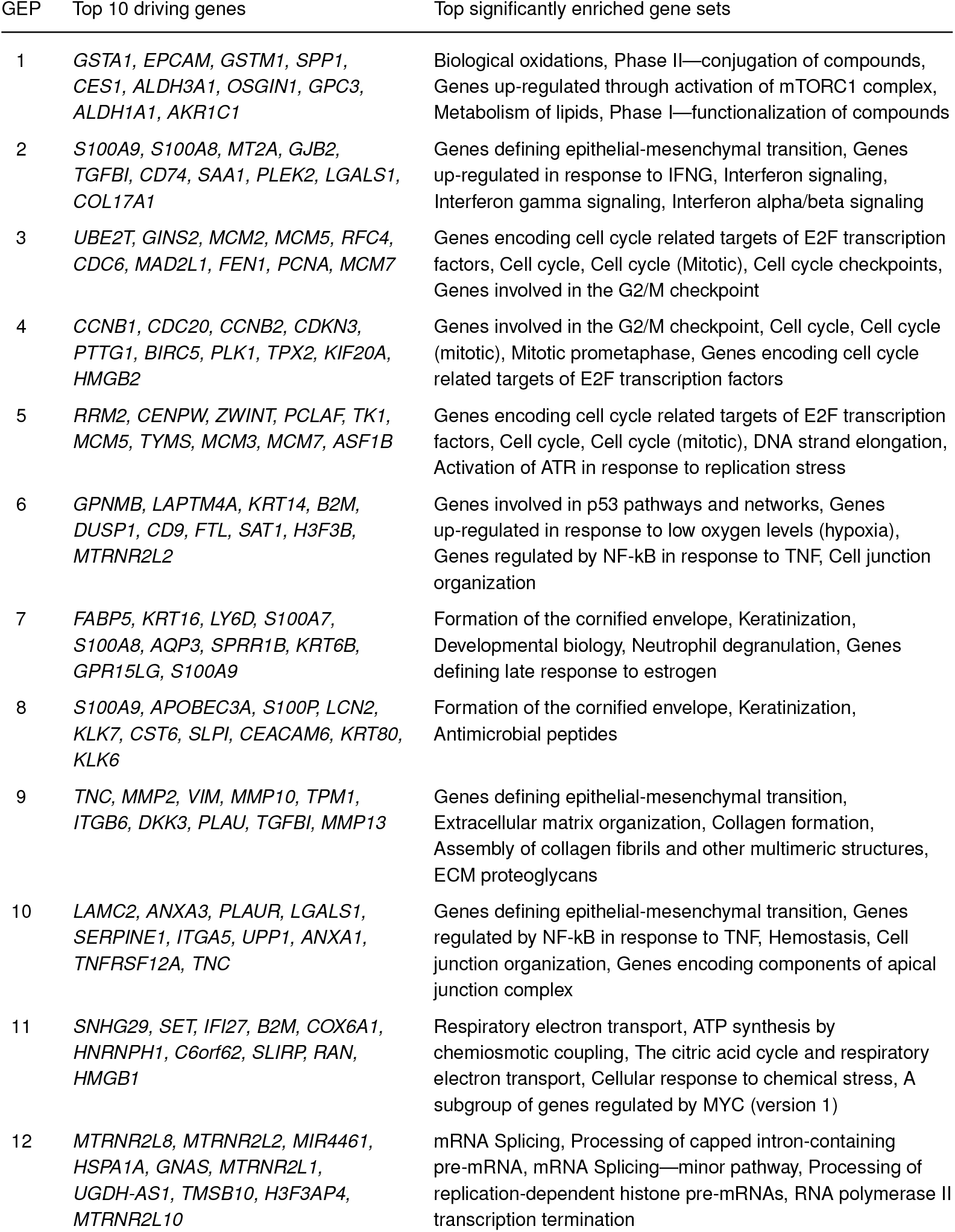
Top driving genes and enriched gene sets of shared GEPs identified in the HNSCC data. The left-hand column lists the genes with the largest increases in expression (LFC) in the GEP. The right-hand column lists the enriched MSigDB Hallmark and CP:Reactome gene sets [39, 43, 44] with Bonferroni-adjusted *p*-value *<* 0.05.

Results of alternative analysis methods, including PCA and NMF-based methods, are summarized in Supplementary Fig. S5–S11. These results highlight behaviors that are qualitatively similar to those seen in the illustrative simulation above; for example, NMF captured the patient-specific programs, but failed to identify the molecular subtypes. PCA successfully identified the molecular subtypes, but it did not separate the patient-specific programs; instead, it reported multiple factors that appear to be random combinations of patient-specific effects. PCA was the only method other than GBCD to identify the molecular subtypes but otherwise produced results quite different to GBCD; indeed PCA failed to recapitulate any of the remaining 10 shared GEPs identified by GBCD (Supplementary Fig. S12). Taken together, these results illustrate the ability of GBCD to identify biologically meaningful expression programs explaining both inter-tumor and intra-tumor transcriptional heterogeneity.

### Pancreatic ductal adenocarcinoma data

PDAC is highly lethal and demonstrates extensive heterogeneity in disease progression and treatment response among patients [45]. Many studies have attempted to define clinically relevant subtypes with the goal of personalizing treatment and improving outcomes [48]. An early study [49] applied NMF to microarray expression data and identified “classical” and “basal” subtypes. Later studies used newer technologies to refine these subtypes: Chan-Seng-Yue et al. [45] identified 4 subtypes — “classical-A”, “classical-B”, “basal-A” and “basal-B” — from an NMF analysis of bulk RNA-seq data in tumor specimens that underwent laser capture microdissection to improve tumor purity; Raghavan et al. [47] refined the basal and classical gene signatures derived from [49] using scRNA-seq data from metastatic patients; Hwang et al. [50] identified a classical and three basal-related expression programs (“basaloid”, “squamoid” and “mesenchymal”) from an NMF analysis of single-nucleus RNA sequencing data in primary tumors.

Understanding transcriptional heterogeneity in PDAC tumors is, therefore, a complex and evolving area. To examine whether GBCD can contribute new insights, we performed a *de novo* analysis of combined scRNA-seq data on PDAC tumors from three recent studies, which together include data on 35,670 malignant cells from 59 PDAC tumors (Table 2). Focusing on malignant cells prevents our analysis of tumor transcriptional heterogeneity from being influenced by a high degree of immune and stromal cell infiltration, which is a particular issue for bulk RNA-seq data analysis of PDAC tumors given their typically low tumor purity [45]. Further, since GBCD can effectively disentangle shared GEPs from patient- and study-specific effects, we expected our analysis to provide a deeper characterization of recurrent gene expression patterns by pooling tumors from multiple studies to achieve a larger sample size.

**Table 2.** PDAC scRNA-seq datasets analyzed. The datasets are labeled as “A”, “B” and “C” in the results. The right-most column gives the number of high-quality malignant cells retained for analysis.

| label | study | assay | specimens | *N* |
| --- | --- | --- | --- | --- |
| A | Chan-Seng-Yue et al. [45] | 10x 3' v2 | 13 primary, 1 metastatic | 18,920 |
| B | Peng et al. [46] | 10x 3' v2 | 24 primary | 10,936 |
| C | Raghavan et al. [47] | Seq-Well | 21 metastatic | 5,814 |

A 2-D embedding of the 35,670 scRNA-seq profiles by *t*-SNE shows that the cells cluster by patient, reflecting substantial inter-patient heterogeneity, and also by cohort, potentially reflecting (expected) technical and biological differences among studies (Fig. 3A). The substantial inter-patient and inter-study structure could obscure biologically interesting expression patterns recurring among the tumors. Applying GBCD to the combined expression data identified a total of 33 GEPs (not including an intercept GEP), whose memberships are shown in Fig. 3B. Consistent with the *t*-SNE analysis, a good number of GEPs identified by GBCD (GEPs 21–33) are each active predominantly in one patient, and thus mainly capture patient-specific effects, in part due to the influences of aneuploid copy number profiles that are common in most human tumors and can vary from patient to patient [51]. The patient-specific GEPs exhibit strong chromosomal structure (Fig. 3B, Supplementary Fig. S13; Online Methods), which suggests a link to copy number variation in tumor cells caused by aneuploidy.

**Fig. 3.**
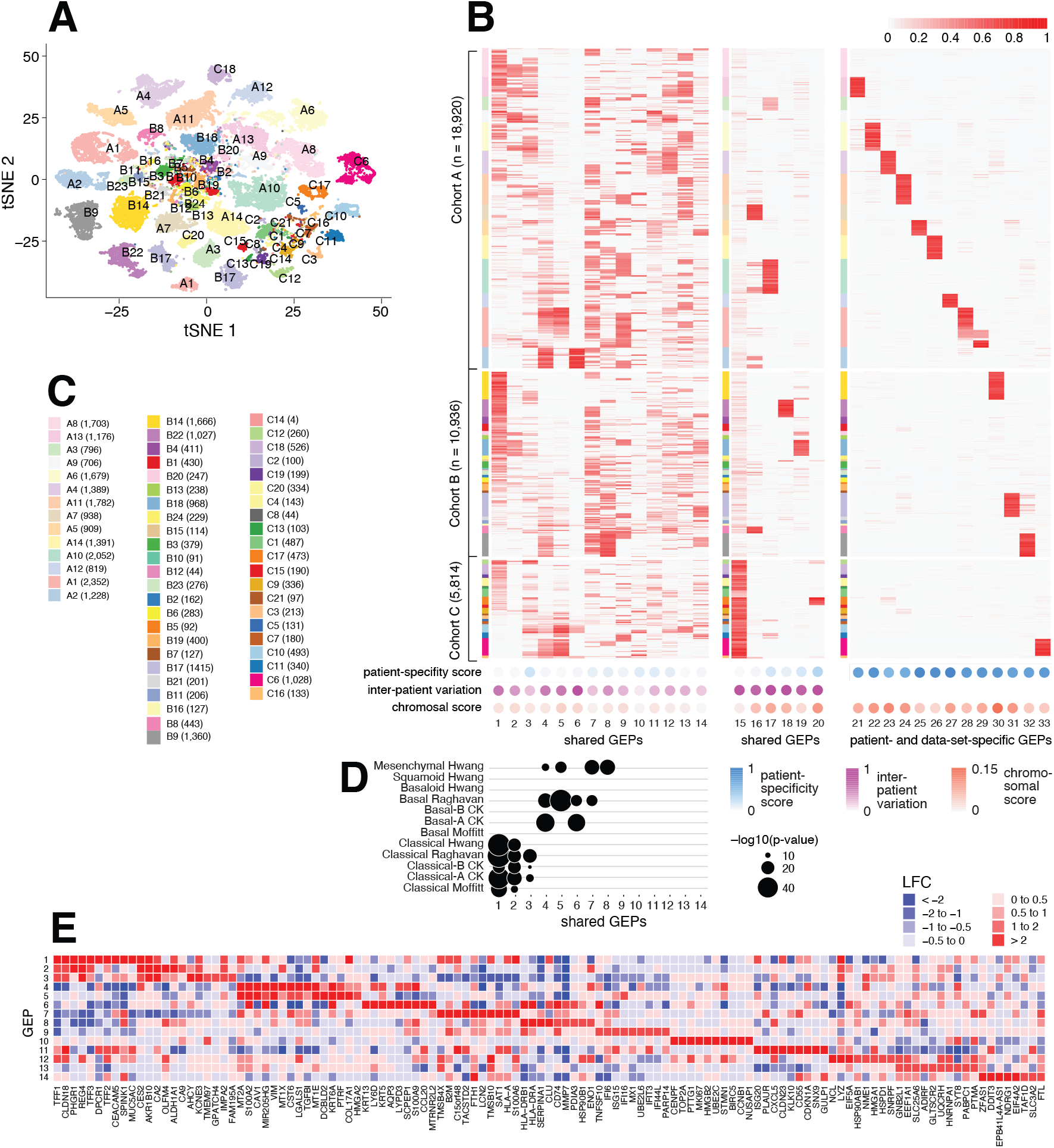
GBCD analysis of PDAC data. Panel A shows the *t*-SNE embedding from scRNA-seq profiles of *N* = 35,670 malignant cells from 59 PDAC tumor samples (Table 2; Panel C). Cells are colored by patient of origin. Two patients (A1, B17) are labeled twice since the cells from those patients were split into two distinct clusters. Panel B is a heatmap showing membership values of the *N* = 35,670 cells (rows) in the 34 GEPs (columns) identified by GBCD. Cells are arranged top-to-bottom by study and patient. Within each study, patients are ordered by the proportion of cells expressing GEP1, which strongly correlates with the classical subtype of PDAC (Panel D). Measures of patient-specific expression (relative to shared expression across patients), inter-patient variation in expression (relative to intra-patient variation) and chromosomal structure of gene signature are shown. See Sections S.1.8 and S.1.9 for definitions. Panel D summarizes concordance between the GBCD-identified shared GEPs (GEPs 1– 14) and literature-derived expression signatures for classical and basal PDAC subtypes, as quantified by *−* log_10_(*p*-value), where *p*-values are produced by one-sided Wilcoxon rank sum test (Online Methods). Literature-derived expression signatures include: Chan-Seng-Yue, Kim et al. [45] (CK), Hwang et al. [50], Moffitt et al. [49], Raghavan et al. [47]. Panel E: LFC estimates for selected genes in the shared GEPs.

Despite the strong inter-tumor expression heterogeneity, GBCD identified many factors (GEPs 1–20 in Fig. 3B) recurring in multiple tumors that represent potentially interesting components of transcriptional variation in PDAC tumors. To interpret the shared GEPs, we identified driving genes (most strongly up-regulated genes) and biological processes enriched for the driving genes of each GEP (Table 3, Supplementary Fig. S19, Supplementary Tables S3–S4, Section S.1.6), then we compared the driving genes with expression signatures from relevant PDAC literature [45, 47, 49, 50]. These analyses did not produce plausible interpretations for GEPs 15–20. GEP15 mainly captures dataset effects and distinguishes cohorts A and B from cohort C based on tumor stage (metastatic vs. primary) and sequencing platform (Seq-Well vs. 10x). GEPs 16–20 are each active in at most a few individuals and demonstrate much less within-patient than between-patient variation, as confirmed by ANOVA (Fig. 3B, Section S.1.8). We thus speculate that GEPs 16–20 capture patterns of copy number variation common to a few patients, which is supported by the fact that GEPs 16–20 display markedly stronger chromosomal structure than GEPs 1–14 (Fig. 3B, Supplementary Fig. S13).

**Table 3.**
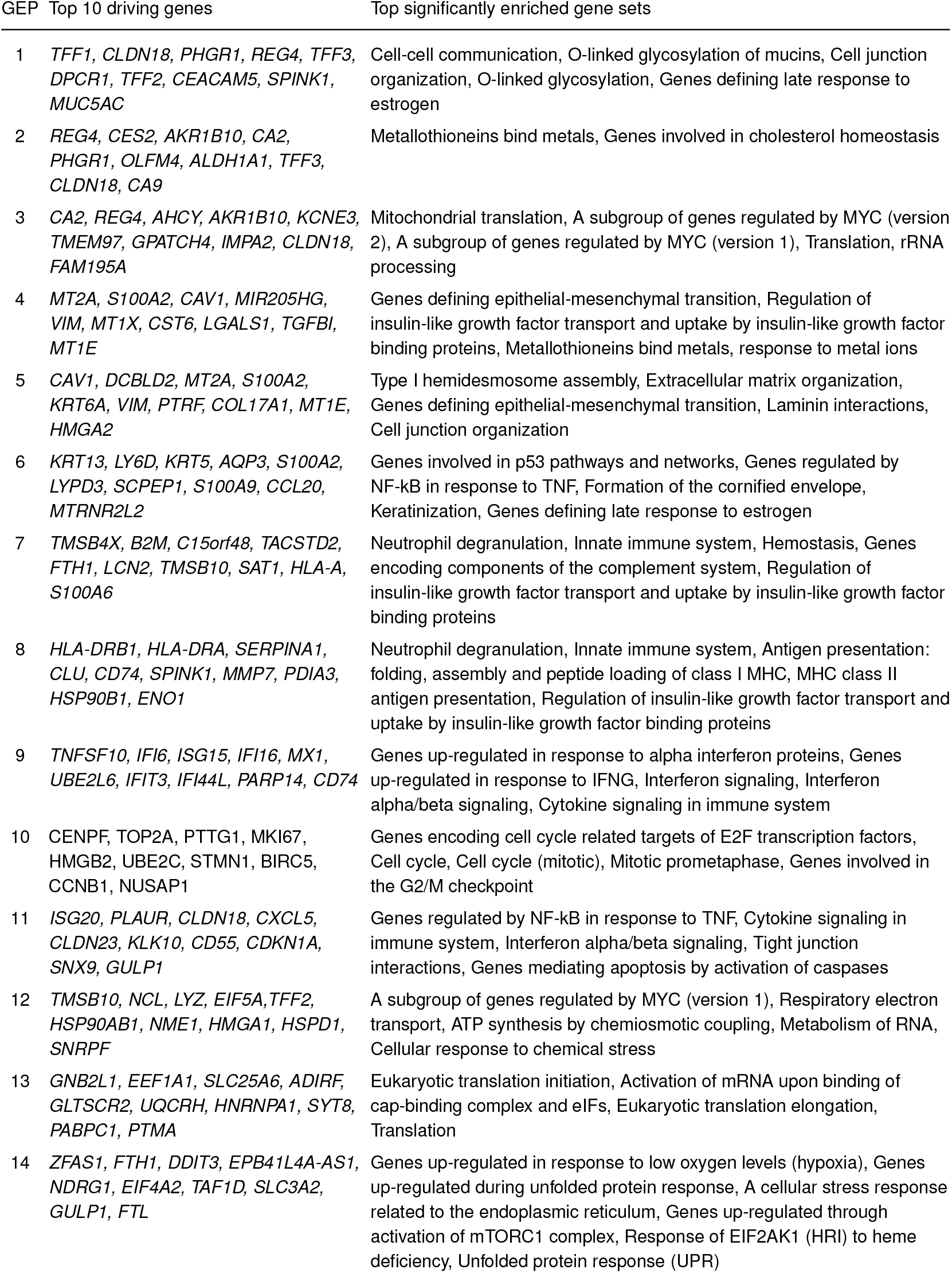
Top driving genes and enriched gene sets of shared GEPs identified in PDAC data. The left-hand column lists the genes with the largest increases in expression (LFC) in the GEP. The right-hand column lists the enriched MSigDB Hallmark and CP:Reactome gene sets [39, 43, 44] with a Bonferroni-adjusted *p*-value *<* 0.05.

A subset of shared GEPs (GEPs 1–8) align well with literature-derived classical or basal-like PDAC subtype signatures, confirming that these subtypes can be identified *de novo* from scRNA-seq data (Fig. 3D). The classical-subtype-associated programs (GEPs 1–3) are enriched for biological processes including O-linked glycosylation of mucins (GEP1), cholesterol homeostasis (GEP2), and MYC regulation (GEP3). GEPs 4–8 are associated with basal-related signatures, and are enriched for many different biological processes, some of them related. GEPs 4 and 5 are both enriched for genes involved in defining the EMT; other enriched processes include regulation of insulin-like growth factor transport (GEPs 4, 7, 8), extra-cellular matrix organization (GEP5), p53 pathway, cornification and keratinization (GEP6), neutrophil degranulation (GEPs 7, 8) and MHC-related processes (GEP8). Many of these enriched biological processes have also been found to be enriched in literature-derived subtype signatures [45, 47, 50, 52]. Notably, we observe at least 2-fold up-regulation of several previously identified marker genes of the classical subtype in at least one of the classical-related GEPs 1–3, including *AGR2, ANXA10, CLDN18, DDC, GATA6, LGALS4, REG4, TFF2*, and of several basal subtype marker genes in at least one of the basal-related GEPs 4–6, including *C16orf74, GPR87, KRT5, KRT6A, ITGA3, PTGES, S100A2* [49, 53, 54]. The classical-associated GEPs 1–2 and the basal-associated GEPs 4–6 are active in two largely exclusive subsets of cells, which sometimes co-occur in the same tumor (e.g., B17), consistent with previous findings that basal and classical programs can co-exist intratumorally [45, 55].

Despite the strong connection between GEPs 1–8 and literature-derived subtype signatures, these results do not simply recapitulate previous findings. Indeed, some previously identified signatures (e.g., the squamoid signatures from [50]) are not strongly correlated with any of our GEPs, and other previously identified signatures reflect some combination of our eight GEPs (Fig. 3D). Thus our GEPs 1–8 represent a new decomposition of classical- and basal-related transcriptional heterogeneity into eight components, with the potential to provide new insights into the disease (see the survival analysis below).

In contrast, GEPs 9–14 do not correlate with previously defined subtype signatures. These GEPs are enriched for interferon signaling (GEP9), cell cycle (GEP10), TNF-NFκB signaling (GEP11), respiratory electron transport (GEP12), translation (GEP13) and stress-induced pathways (GEP14).

#### A stress signaling program is strongly prognostic of poor survival

To assess the prognostic value of GEPs 1–14 in the context of previously identified subtype signatures, we scored each gene expression signature in bulk RNA-seq data of 391 resected primary PDAC tumors from 4 studies [52, 56–58] involving 260 deaths, and performed survival analysis (Section S.1.10). Specifically, we performed stepwise variable selection among all gene signatures, including the previously identified subtype signatures, in a Cox proportional hazards regression model with overall survival as the endpoint, adjusting for age, sex, and tumor stage [59]. The final model selects two of our GEPs, GEP4 and GEP14, as prognostic (at *p*-value *<* 0.01) but selects none of the previously identified subtype signatures (Fig. 4A). GEP4 is prognostic of poor survival (HR = 1.58, *p*-value = 2 *×* 10^*−*10^) and is the only selected program related to subtype (basal), indicating that it is more strongly prognostic of survival than previously defined subtype signatures. GEP14 is also prognostic of poor survival (HR = 1.32, *p*-value = 5 *×* 10^*−*5^), and represents a novel expression signature that is predictive of prognosis even when accounting for tumor subtype and tumor stage. Further analyses, stratified by tumor stage and classical/basal subtype, show that GEP14 is particularly prognostic of patient survival in earlier stage basal-like tumors (Fig. 4C).

**Fig. 4.**
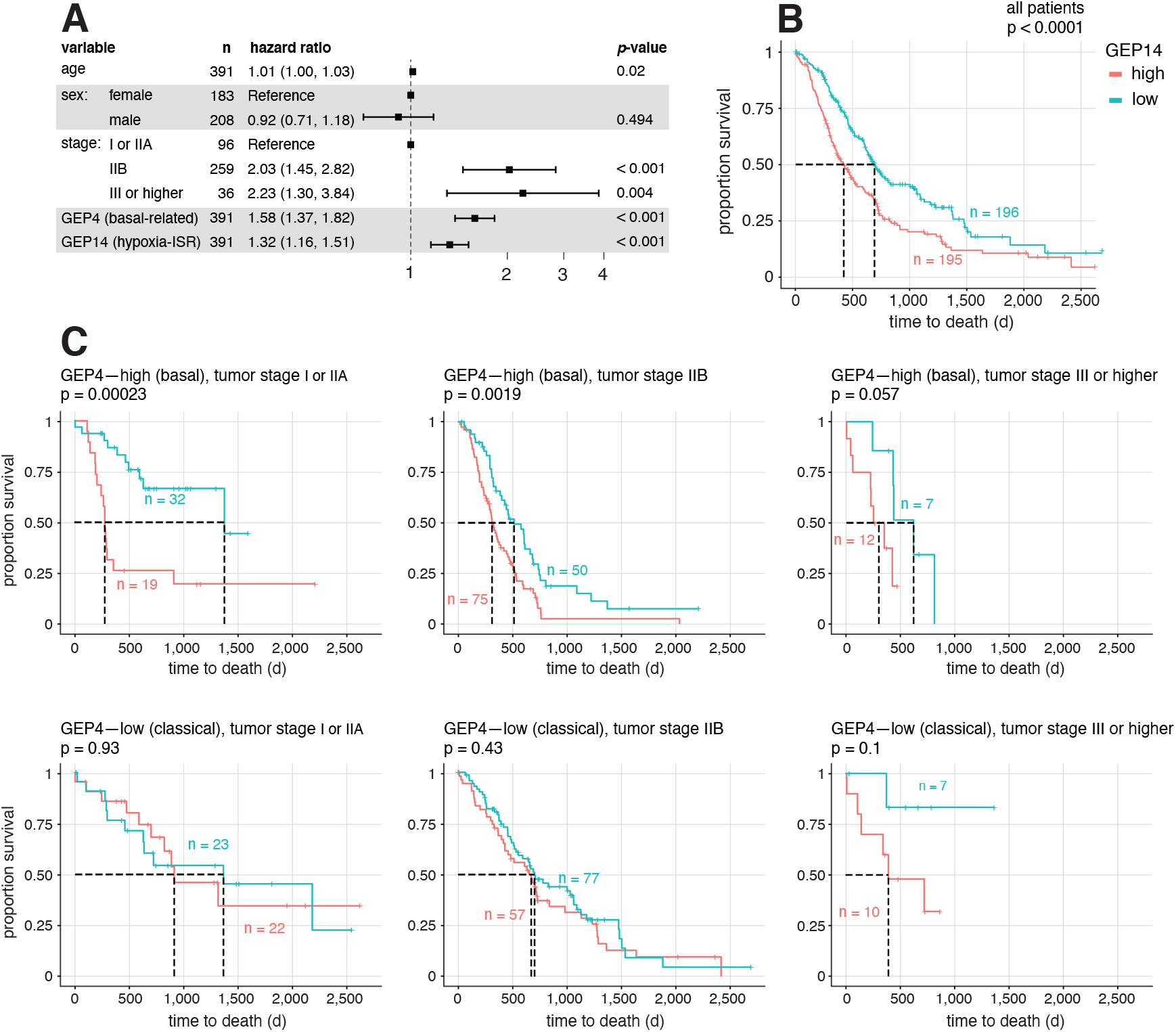
Assessment of prognostic relevance of expression program signatures in bulk RNA-seq data from 391 primary PDAC tumors. Panel A shows the final model selected by stepwise variable selection in Cox regression analysis, with overall survival as endpoint and all gene expression signature scores as covariates to be selected. Age, sex and tumor stage are always included as covariates in the model. For each covariate included in the model, hazard ratio and its 95% confidence interval are presented. Panel B and C show the Kaplan-Meier plots comparing the overall survival between patients with relatively high or low levels of GEP14, together with two-sided log-rank test *p*-value. This comparison is done for all 391 patients in Panel B, and separately for subgroups of patients stratified by tumor subtype (dichotomized GEP4) and tumor stage in Panel C.

Inspection of the top driving genes in GEP14 reveals interesting connections with previous literature on PDAC. The top driving gene is *ZFAS1*, a long non-coding RNA gene that has been shown to promote PDAC metastasis [60]. Other top driving genes include two genes related to iron metabolism, *FTH1* and *FTL*, both of which have been recently linked with PDAC progression arising from increased ferritinophagy [61, 62].

Enrichment analysis of a longer list of driving genes (Supplementary Fig. S19) highlights several gene sets related to stress response. These include genes up-regulated in response to hypoxia (e.g., *BNIP3L, DDIT3, VEGFA*), in response to heme deficiency, and in response to misfolded protein accumulation in the endoplasmic reticulum. Notable driving genes related to stress response include activating transcription factor 4 *ATF4* — the main effector of the Integrated Stress Response [63] — and several of its validated targets, including *DDIT3/CHOP, HSPA5, MAP1LC3B, MTHFD2, SQSTM1* [64–66]. Among these, *SQSTM1* and *MAP1LC3B* are key modulators of autophagy [65, 67], which is known to promote PDAC progression and therapy resistance [68, 69], and *HSPA5* has been previously linked with PDAC progression through decreased ferroptosis [70, 71].

### Computing requirements

GBCD builds on the general EBMF framework, and the fast model fitting algorithms implemented for EBMF makes it possible to apply GBCD to moderately sized data sets (e.g., the HNSCC data). However, it can be costly to explicitly compute the matrix ***Y Y*** ^*T*^ and then apply EBMF to that *N × N* (non-sparse) matrix for large data sets (e.g., the PDAC data). Therefore, we implemented the ability to apply EBMF to a matrix ***Y Y*** ^*T*^ *by performing computations using only the (sparse) matrix* ***Y*** (i.e., without ever explicitly forming the matrix ***Y Y*** ^*T*^), which is mathematically equivalent to the former but reduces computing costs when *N* is large and ***Y*** is sparse. This approach reduces the per-iteration computational complexity from *O*(*N* ^2^*K*) to *O*(*SK*), and reduces the memory usage from *O*(*N* ^2^) to *O*(*NK* + *S*), where *S* is the number of non-zero entries in ***Y*** . For example, for the HNSCC data (2,176 cells, 17,113 genes), this approach reduced the running time and memory from 2 hours and 16 GB to 50 minutes and 2 GB; for the PDAC data (35,670 cells, 13,844 genes), this approach reduced computing time and memory from 50 hours and 200 GB to 21 hours and 48 GB. In this paper, we gave results for the approach that explicitly computes ***Y Y*** ^*T*^, but we note that results were highly consistent between the two approaches (Supplementary Fig. S17). Computing costs can also be reduced by optimizing the number of backfit iterations for estimating ***L*** (see Section S.1.11). We note that further improvements to the GBCD pipeline will likely be needed to analyze very large scRNA-seq datasets.

## Discussion

In this paper, we introduced a new matrix factorization method, GBCD, and demonstrated its ability to provide new insights into transcriptional heterogeneity in scRNA-seq tumor data from multiple patients and studies. A key feature of GBCD is that it assumes expression programs to be orthogonal to one another, which helps avoid absorbing shared components of expression variability into patient-specific expression programs. When cells show strong inter-patient heterogeneity in their transcriptional profiles, as is typical of malignant cells, we demonstrated in both simulated and real data that GBCD can capture shared patterns of variation that existing MF methods miss. In contrast to NMF, GBCD does not constrain expression programs to be nonnegative, which allows it to model transcriptional repression of genes in a given program, as well as up-regulation.

In applications to scRNA-seq tumor data from PDAC and HNSCC patients, GBCD identifies GEPs that are shared across tumors and datasets, as well as GEPs that characterize patient and dataset effects. Some of the shared GEPs correspond to previously defined tumor subtypes or generic cellular processes such as cell cycle and immune activation, but others represent new findings. Most notably, we identified a new expression program (GEP14) that is an independent predictor of poor prognosis in patients with primary PDAC. GEP14 contains many genes related to stress response, including hypoxia. Hypoxia gene expression signatures have been previously related to worse PDAC outcomes and hypoxia is a promising target of innovative therapeutic strategies [72–76]. However, GEP14 is also associated with many other stress-response processes, including many genes that are induced by *ATF4* as part of the Integrated Stress Response (ISR). The ISR is activated in response to stress, including hypoxia, by phosphorylation of the *α* subunit of eukaryotic initiation factor 2 (*eIF2α*) at serine 51, which suppresses global protein synthesis to reduce energy expenditure while enhancing translation of specific mRNAs, including *ATF4*. This then up-regulates the expression of many genes involved in stress adaptation [66]. Recently, *ATF4* has been shown to promote amino acid biosynthesis and tumor cell survival in PDAC [77] while *eIF2α* phosphorylation significantly correlates with tumor recurrence/metastasis and shorter overall survival [78]. Interestingly, the *ATF4* expression signature has been found to be a predictor of therapy resistance in other cancers. For example, it was revealed to be induced in melanoma cells that escaped RAF kinase inhibitor treatment in a recent scRNA-seq study [79]; tumor cells that persisted after treatment with BH3 mimetics similarly depended on *ATF4* and the ISR to survive [80]. The fact that GEP14 includes many genes induced during the ISR, and is an independent predictor of PDAC patient survival, suggests an important role for the ISR in PDAC progression, and raises the potential that pharmacological interventions targeting the ISR could provide useful therapeutic strategies. More work is needed to test these hypotheses.

Like other matrix factorization and clustering methods, GBCD is perhaps best viewed as a useful exploratory tool for identifying interesting structures in large datasets. And, like existing tools, GBCD has limitations that users should be aware of. For example, GBCD is solving a non-convex optimization problem, and so its solutions will typically depend on initialization, and it may be very difficult to find the global optimal solution. Indeed, although in our examples GBCD finds many GEPs corresponding to individual patient effects, there are some patients for which it finds no patient-specific program, and we believe this is likely a failure of the optimization routine to identify such programs. Applications of NMF sometimes try to ameliorate this issue by combining results across multiple runs with different initializations [5], and it is possible that similar strategies could be helpful for GBCD.

Also like other matrix factorization methods, GBCD requires user judgement in choosing *K*, the number of factors. The EBMF method that underlies GBCD requires users to set an upper bound, *K*_max_, on the number of factors, and then provides a method to automatically select *K* (up to *K*_max_). Thus, one possible strategy is to simply set *K*_max_ to be very large, and rely on this automatic method to select *K*. However, for several reasons — not least that computation increases with *K* — in practice users might want to more strictly limit *K*. In experimenting with various values of *K* in our analyses, we found that GEPs identified with a smaller *K* are generally also present when a larger *K* is used (Supplementary Fig. S16). Generally speaking then, a larger *K* allows for identifying finer structure in tumor transcriptional heterogeneity at the expense of higher computational cost, and the main downside to limiting *K* is that one may miss interesting structure in the data. In practice the value of *K* needed to capture most of the interesting structure will inevitably depend on the complexity of the data, and this will likely increase with the number of cohorts, patients and tumor cells analyzed.

Finally, we note that we do not expect GBCD to be uniformly superior to other methods, such as NMF, in all applications. The assumptions of GBCD, including orthogonality of GEPs, may be more appropriate in some applications than others. Since it may be difficult to know in advance whether one set of assumptions or another are more appropriate to a given dataset, it is common to apply multiple methods to see how the results differ. Our results here show that GBCD can produce results that are quite different from existing methods, and in so doing has the potential to provide new scientific insights.

### URLs

The code and data resources for simulations and analysis of HNSCC and PDAC data are stored in the Zenodo repository https://doi.org/10.5281/zenodo.8271036. Other R packages used in this work include: gbcd (https://github.com/stephenslab/gbcd), ebnm (https://github.com/stephenslab/ebnm), ashr (https://github.com/stephens999/ashr), flashier (https://github.com/willwerscheid/flashier), Seurat (https://satijalab.org/seurat/), fast-Topics (https://github.com/stephenslab/fastTopics), NNLM (https://github.com/linxihui/NNLM), rliger (https://github.com/welch-lab/liger), cNMF (https://github.com/dylkot/cNMF), batchelor (https://bioconductor.org/packages/devel/bioc/html/batchelor.html), conos (https://github.com/kharchenkolab/conos), splatter (https://github.com/Oshlack/splatter), scran (10.18129/B9.bioc.scran) seqgendiff (https://github.com/dcgerard/seqgendiff), and infercnv (https://github.com/broadinstitute/infercnv).

## Acknowledgements

This work was supported by NIH grants R01HG002585, DP2AI145100, U01AI160418, and a Gut Cell Atlas grant from The Leona M. and Harry B. Helmsley Charitable Trust. We thank the staff at the Research Computing Center at the University of Chicago for providing the high-performance computing resources used to implement the numerical experiments. This study was conducted using data provided by the Ontario Institute for Cancer Research which is supported by funding provided by the Government of Ontario. The views expressed in the publication are those of the author and do not necessarily reflect those of the Ontario Institute for Cancer Research or the Government of Ontario.

## Author contributions

Y.L. and M.S. conceived the project. Y.L. and J.W. developed the method and implemented it in R under supervision of M.S. Y.L. performed the simulation study and analyses of real data. Y.L. and P.C. prepared the online code. Y.L., S.A.O. and K.F.M. interpreted the PDAC data analysis results with contributions from P.C., J.W. and M.S. All authors wrote the manuscript.

## S.1. Online methods

### S.1.1. GBCD analyses

In this section, we detail the steps that were taken to fit a GBCD model to a dataset and then obtain statistical quantities used in interpretation of the GBCD results.

#### S.1.1.1. Preparation of scRNA-seq data

In all our GBCD analyses, the initial data were combined into an *N × J* counts matrix ***X*** across tumors, cohorts, etc. In most cases, ***X*** was a matrix of unique molecular identifier (UMI) counts in which element *x*_*ij*_ was the UMI count of gene *j* in cell *i*. Here we describe the steps that were taken to convert the initial data ***X*** to the matrix ***Y*** that was then analyzed by GBCD.

Let *s*_*i*_ denote the “size factor” for cell *i*. For UMI count data, we took this size factor to be the total count over all genes, 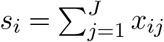 . (We note that other size factor definitions [81–83] are possible, and in some settings may be preferred.) We scaled the data by the size factors, added a “pseudo-count” *c >* 0, then took the log:

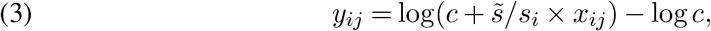

in which 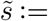 median*{s*_1_, …, *s*_*N*_ *}* denotes the median size factor. We subtracted log *c* from the log-transformed data, which ensured that all *y*_*ij*_’s were non-negative. This also preserved sparsity in the original counts *x*_*ij*_; that is, *y*_*ij*_ was zero if and only if *x*_*ij*_ was zero.

Regarding the choice of pseudo-count *c*, different pseudo-counts have been proposed (e.g., [84, 85]). Generally speaking, pseudo-counts that are too small overemphasize differences in normalized expression between zero and nonzero UMI counts, whereas pseudo-counts that are too large reduce differences in normalized expression [85]. We chose a reasonable “middle ground” value of *c* = 0.1; see [36, 86] for further discussion.

Unlike the other datasets, the HNSCC data were not UMI counts; rather, they were read counts produced by SMART-Seq2 [87]. Following [10], we defined the transformed counts as *y*_*ij*_ = log_2_(1 + TPM_*ij*_*/*10), where TPM_*ij*_ was the transcript-per-million (TPM) value [88, 89] for gene *j* in cell *i*. Note that this transformation is equivalent to (3) (up to a constant of proportionality) with 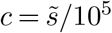 .

#### S.1.1.2. Estimating GEP memberships

In GBCD, we estimate the GEP membership matrix ***L*** in the matrix factorization (2) by decomposing ***Y Y*** ^*T*^ using the empirical Bayes matrix factorization (EBMF) framework [35]. In detail, we model ***Y Y*** ^*T*^ as

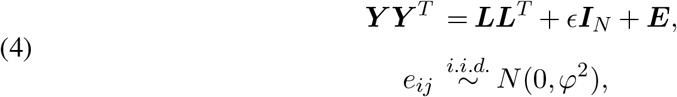

where ***I***_*N*_ is the *N × N* identity matrix, ***E*** is an *N × N* matrix with entries *e*_*ij*_, and *ϵ, φ*^2^ *>* 0 are additional unknowns to be estimated.

If the rows of ***Y*** are centered, ***Y Y*** ^*T*^ is proportional to the sample covariance matrix that characterizes the covariation between the transcriptional profiles of every pair of cells. Thus, we refer to (4) as a “covariance decomposition” since it approximates ***Y Y*** ^*T*^ by ***LL***^*T*^, where ***L*** is an *N × K* nonnegative matrix.

GBCD assigns a “generalized binary” prior independently to each entry of ***L***,

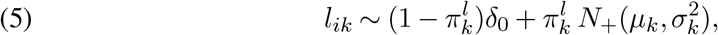

in which *N*_+_(*µ, σ*) is a left-truncated normal distribution with mean *µ* and variance *σ*^2^, with the truncation point at zero. The functional form of this prior is chosen to balance modelling flexibilty (ability to capture both discrete and continuous structures) and computational convenience. We call (5) a “generalized binary” (GB) prior because as 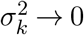, the prior’s support takes only two possible values, 0 or *µ*_*k*_. A strictly binary prior — that is, (5) with *σ*_*k*_ = 0 — is suitable for modeling discrete structures in cell populations, such as distinct molecular subtypes and patient-specific effects, but cannot capture transcriptional variation of a continuous nature, such as biological processes with varying degrees of activity in cells. To get around this limitation, we have proposed the GB prior in which the ratio *σ*_*k*_*/µ*_*k*_ is kept small by taking some pre-specified small value (see below), which encourages discrete structures while still allowing for continuous structures when they fit the data better. Rather than set the parameters 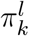 and *µ*_*k*_ by hand, we take an empirical Bayes approach and leverage the information across cells to adapt the priors to the data. The exact steps we take to estimate these parameters are described below.

Note that the diagonal matrix ***D*** from (2) does not appear in (4). This is because we assume that entries of ***L*** in (2) are *a priori* close to 0 or 1 (rather than 0 or *µ*_*k*_) for ease of illustration, and so need to incorporate the scaling factors *µ*_*k*_’s into 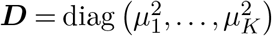 .

Our current EBMF implementation cannot fit ***Y*** *≈* ***LF*** ^*T*^ subject to ***L*** = ***F*** . Therefore, instead of fitting (4), in practice we fit a slightly relaxed matrix decomposition,

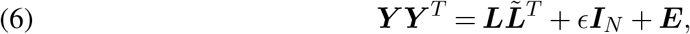

with independent GB priors (5) on ***l***_*k*_ and 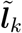 Clearly, (6) is the same as (4) when. While we do not enforce ***L*** and 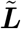 to be the same, in practice, we observe that fitting always produces highly concordant ***l***_*k*_ and 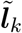 for the majority of components; in our implementation we only keep components *k* such that the Pearson correlation between ***l***_*k*_ and 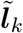 is greater than 0.8, and use ***l***_*k*_’s as GEP membership estimates.

We now describe how to fit the EBMF model (6) with a GB prior (5). Thanks to the modularity of the model fitting algorithm, solving the EBMF problem for any prior family requires only solving a series of much simpler “empirical Bayes normal means” (EBNM) problems with the same prior family (see [35] for the connection between EBMF and EBNM). For an EBNM problem with a GB prior, we have observations ***x*** = (*x*_1_, …, *x*_*n*_)^*T*^ that follow a multivariate Gaussian distribution with unknown means ***θ*** = (*θ*_1_, …, *θ*_*n*_)^*T*^ and known covariance matrix diag 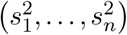, where *θ*_1_, …, *θ*_*n*_ are i.i.d. from a GB distribution. That is,

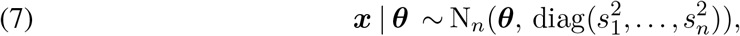

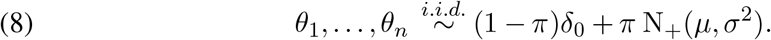

Solving this EBNM problem involves the following two steps.

1. Estimate *µ, σ*^2^, *π* by maximizing the marginal likelihood

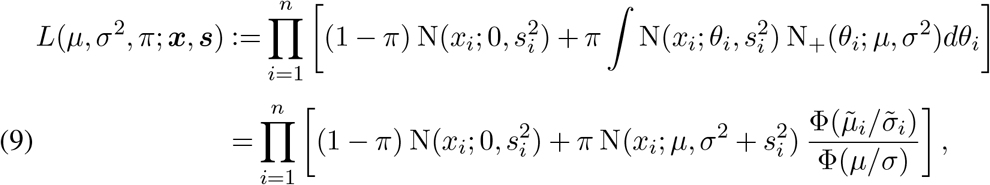

where N(·; *µ, σ*) and N_+_(·; *µ, σ*) respectively denote the probability density function of N(*µ, σ*^2^) and N_+_(*µ, σ*). Φ(·) denotes the cumulative distribution function of a standard normal random variable; 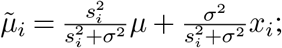 and 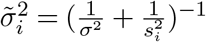 . In practice, we assume that *σ/µ* = *ω* takes some pre-specified small value, e.g., *ω* = 0.02, so we just need to maximize the marginal likelihood in (9) over *µ* and *π*. To facilitate solving this optimization problem, for each observation *i*, we define a latent indicator *z*_*i*_ *∈ {*0, 1*}* that is 1 if *θ*_*i*_ is drawn from N_+_(*µ, σ*^2^) and 0 otherwise. With this data augmentation, we maximize the complete data log-likelihood (10)

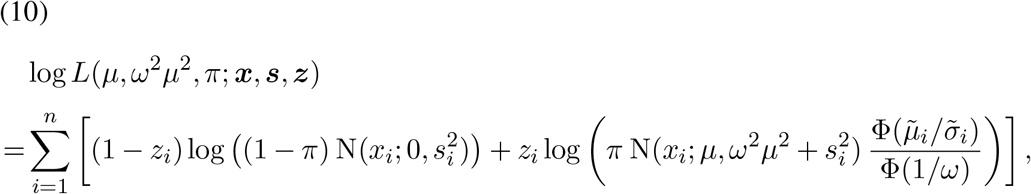

using EM [90], which alternates between the following two steps at each iteration *t* = 1, 2, … until convergence.
  - Given the current estimates of the parameters *π*^(*t*)^, *µ*^(*t*)^, use Bayes theorem to calculate the conditional expectation of *z*_*i*_, i.e., 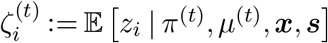 .
  - Given the current 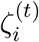, update the estimates of *π, µ* respectively by:

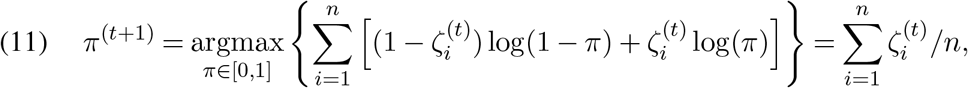

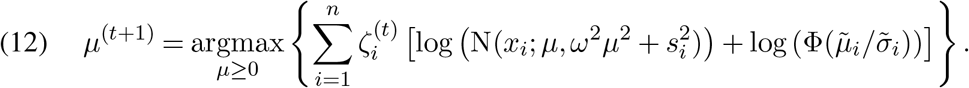

The constrained one-dimensional optimization in (12) does not have a closed-form solution, so we solve it using the R function `optim` with method “L-BFGS-B”. We denote the converged estimates of *µ, π, ζ*_*i*_ by 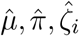.
2. Compute the posterior distribution of ***θ*** given 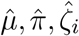 by

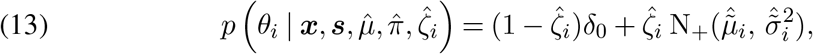

where 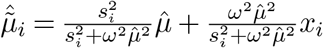 and 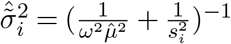 . Finally, we fit the EBMF model (6) with GB prior (5) by passing the above EBNM solver as argument `ebnm.fn` to function `flash` in R package `flashier` [91]. More specifically, fitting (6) consists of the following 4 steps (i) – (iv). ***L*** and 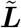 are initialized by steps (i) – (iii), which are refined by a backfitting algorithm performed in step (iv).
  i. Add pairs of 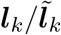 in a greedy manner, using the function `flash_greedy`, where independent point Laplace priors are placed on *l*_*ik*_ and.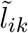.
  ii. Backfit existing pairs of 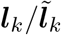 (i.e., by updating each pair while keeping others fixed in a cyclical fashion), using the function `flash_backfit`, until convergence or completion of *T*_1_ iterations.
  iii. Split each ***l***_*k*_ into two nonnegative factors 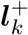 and 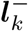, where 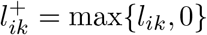 and 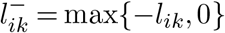, so we have 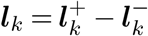 . Do the same for 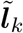 to obtain 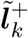 and 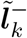.
  iv. Backfit existing pairs of 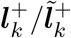 and 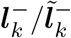 using the function `flash_backfit`, where independent GB priors are placed on 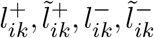, until convergence or completion of *T*_2_ iterations.

#### S.1.1.3. Estimating GEP signatures

We estimate the GEP signature matrix ***F*** by fitting an EBMF model to ***Y*** with fixed ***L***, which is obtained via the covariance decomposition step in *S.1.1.2*,

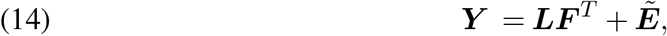

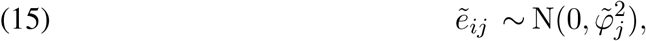

where 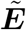 is an *N × J* matrix of independent error terms 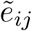, and 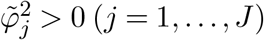 are additional unknowns to be estimated. We note that model (14) is a semi-nonnegative matrix factorization (SNMF) [92], because it places a nonnegativity constraint on ***L*** but not on ***F*** . We also note that fitting (4) is, in principle, equivalent to fitting (14) with an orthonormal constraint on ***F*** (i.e., ***F*** ^*T*^ ***F*** = ***I***_*K*_), which can be seen by post-multiplying each side of (14) by its transpose.

For a GEP *k*, ***f***_*k*_ characterizes the patterns of co-regulation of all genes. It is reasonable to assume that most genes do not exhibit differential expression in GEP *k* compared to other GEPs, so we put a point Laplace prior on *f*_*jk*_, independent over *j* and *k*, which encourages shrinkage of small coefficients to 0 but avoids over-shrinkage of large coefficients.

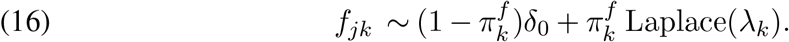

Similar to S.1.1.2, the component-specific hyper-parameters 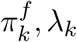 of the point Laplace prior are estimated from data by pooling information across all genes using an empirical Bayes method. For each *k*, the genes with largest (and positive) estimates of *f*_*jk*_ are interpreted as the genes driving the GEP *k*. The EBMF model (14) with point Laplace prior (16) is fit using `flashier`, where we call the function `flash` and set argument `ebnm.fn = ebnm::ebnm_point_laplace`. ***F*** ^*T*^ is initialized by (***L***^*T*^ ***L***)^*−*1^ ***L***^*T*^ ***Y*** .

#### S.1.1.4. Quantifying uncertainty in the GEP signatures

Since mean field variational approximations, which are used in the EBMF framework, are known to underestimate uncertainty in posterior distributions [35], we performed additional calculations to obtain better uncertainty quantifications of the LFC estimates, which we then used to compute posterior *z*-scores and other posterior quantities for *f*_*jk*_.

First, for each gene *j*, we fit a linear regression model in which the log-pc data for gene *j*, ***y***_*j*_ = (*y*_1*j*_, …, *y*_*Nj*_)^*T*^, was the response variable, and the regressor matrix was the matrix of GEP membership estimates ***L***. We fit the linear regression models using the `lm` function in R. After fitting each gene-wise regression model, we extracted the point estimates 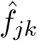 and standard error estimates *Ŝ*_*jk*_ for the corresponding regression coefficients.

Next, we performed adaptive shrinkage, separately on the LFC estimates for each GEP *k*, to improve the estimation accuracy of the standard errors. To implement this step, we use the `ash` function from the `ashr` R package [93]. This adaptive shrinkage method takes as input a vector of effect estimates 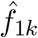, …, 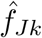 and associated standard errors *Ŝ*_1*k*_, …, *Ŝ*_*Jk*_. The revised posterior mean estimates and standard errors returned by the adaptive shrinkage method are then used to calculate the final test statistics, including posterior *z*-scores (defined as the posterior mean divided by the posterior standard error [94]) and local false sign rates, or *lfsr* [93], which are then used in the volcano plots in the Supplementary Fig. S18 and S19.

Note that this approach to quantifying uncertainty of LFC estimates assumes that the membership estimates are “known”, so does not fully account for uncertainty in the matrix factorization, but in practice it is an improvement over the uncertainty estimates provided by `flashier`.

### S.1.2. Details for simulation design

We first simulated a UMI count matrix ***X***^null^ containing scRNA-seq measurements of *J* = 10, 000 genes in *N* = 3, 200 cells using the Splatter framework [95],

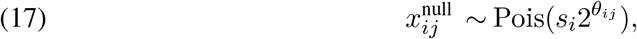

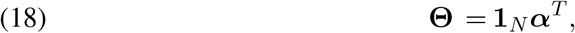

where ***α*** is a *J ×* 1 vector of gene-specific intercept terms that represent the relative expression levels for each gene. The parameters are generated from the Splat simulation model [95] using input parameters estimated from one PDAC dataset. Then we modified ***X***^null^ using binomial thinning [96] to produce ***X*** such that

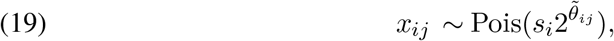

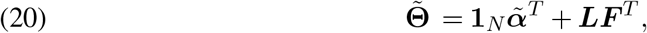

where 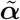 is a *J ×* 1 vector of modified gene-specific intercept terms due to thinning, and ***L*** and ***F*** are respectively *N × K* and *J × K* matrices representing cell memberships and gene signatures for *K* = 11 GEPs. Fig. 1A displays ***L*** for one simulation dataset.

Each GEP is characterized by up-regulation (relative to the “baseline” level of expression determined by 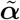) of 75 to 500 non-overlapping genes, with a fold change having an average of 3 and ranging from 1.5 to 11 in magnitude. We repeated the above procedure to generate a total of 20 replicate datasets.

### S.1.3. Implementation details for MF methods

Here we describe in detail how we ran the alternative MF methods that were implemented in the simulations.

- **Patient-by-patient NMF:** We fitted a NMF with 3 programs (an intercept term included) to the UMI counts, separately for each patient, by calling the `nnmf` function from the R package `NNLM` [97] with its default settings. Then we compared all programs by hierarchical clustering, using the fraction of top 50 genes (quantified by the Jaccard index [15]) that are shared between every pair of programs as the similarity measure. Following [10], we manually identified clusters of highly correlated programs, which were defined as “meta-programs”. For a given meta-program, the cell-wise memberships of all patients are calculated as follows. If a patient is included in the associated cluster of programs, the cell-wise membership estimates come from the loadings of the corresponding program of that patient; otherwise the cell-wise membership estimates are set to zero.
- **Combined NMF (UMI counts):** We fitted a Poisson NMF to the UMI counts ***X***, which is equivalent to fitting a multinomial topic model [7, 98, 99], by calling the `fit_topic_model` function from the R package `fastTopics` [100] with its default settings.
- **Combined NMF (log-pc counts):** We fitted NMF with mean squared error loss function to ***Y*** by calling the `nnmf` function from the R package `NNLM` [97] with its default settings.
- **Consensus NMF (UMI counts):** We ran consensus NMF [5] with Frobenius loss to ***X*** using the Python package `cNMF` with its default settings.
- **PCA**: We performed PCA on the log-pc counts ***Y*** by calling the R function `svd`.
- **EB-SNMF, point-exp prior or GB prior:** we used the R package `flashier` to fit the model (14) to the log-pc counts ***Y***, with a point Laplace prior (16) on ***F***, and either a GB prior (5) or point-exponential prior on ***L***. No orthogonality constraint is placed on ***F***, which is the key difference from GBCD. More specifically, fitting EB-SNMF involves the following 4 steps. ***L*** and ***F*** are initialized by steps (i) – (iii), which are refined by a backfitting algorithm in step (iv).
  i. Add pairs of ***l***_*k*_*/****f***_*k*_ in a greedy manner, using the function `flash_greedy`, where independent point Laplace priors are placed on *l*_*ik*_ and *f*_*jk*_.
  ii. Backfit existing pairs of ***l***_*k*_*/****f***_*k*_ (i.e., by updating each pair while keeping others fixed in a cyclical fashion), using the function `flash_backfit`, until convergence or completion of *T*_1_ iterations.
  iii. Split each ***l***_*k*_ into two nonnegative factors 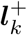 and 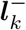, where 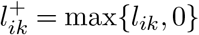 and 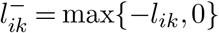, so we have 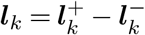 .
  iv. Backfit existing pairs of 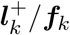 and 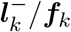 using the function `flash_backfit`, where independent GB (or point exponential) priors are placed on [inrq] and independent point Laplace priors are placed on *f*_*jk*_, until convergence or completion of *T*_2_ iterations.
- **CD, point-exp prior:** we ran a in which of GBCD where we replaced the GB prior (5) on ***L*** with a point exponential prior. The implementation is essentially the same as that of GBCD, as described in detail in Sec. S.1.1.

#### Choice of K in the simulation study

Determining the number of components *K* is generally a challenging task for any matrix factorization method. For NMF to UMI counts, we set *K* to its true value 11. For NMF to log-pc counts, we set *K* = 12 to allow the inclusion of an additional component that models the gene-specific baseline expression levels. For consensus NMF to UMI counts, *K* is determined from 8–12 based on the trade-off between stability and error. For PCA to log-pc counts, we set *K* = 8, 10, 12 to assess its performance when different numbers of principal components are included, and the PCA result shown in Fig. 1 was produced using *K* = 10. All EB-based methods, including GBCD, can perform automatic selection of *K* [35]. In particular, users of `flashier` just need to specify an upper bound *K*_max_ for *K*, among which the components whose prior is estimated to be a point mass at 0 are essentially removed from the model. We set *K*_max_ = 16 to implement all EB-based methods.

### S.1.4. Implementation details for harmonization methods

We implemented several widely used batch effect correction methods which were specifically developed to integrate scRNA-seq data from multiple sources. In particular, we used each of these methods to correct for patient effects in the simulated data, and assessed their performance in recovering the subtype-related and continuous GEPs which are shared across multiple patients. Here we describe in detail how these methods were implemented in the simulations.

#### LIGER

We implemented Liger [25] on the UMI counts using the R package `rliger` (version 1.0.0), following the suggested pipeline to integrate multiple scRNA-seq datasets as specified in the package vignette. We set the inner dimension of factorization `k=20` when calling the `optimizeALS` function. We assessed the impact of the regularization parameter *λ*, which controls the amount of penalization for patient-specific effects, on its performance (Supplementary Fig. S4). The Liger result shown in Fig. 1 was produced using the default value of *λ* = 5.

#### CCA (Seurat)

We implemented Seurat 3 integration [26] on the log-pc counts using the R package `Seurat` (version 4.1.1), following the standard integration workflow described in the package vignette. We first identified the top 5000 highly variable genes separately for scRNA-seq data from each of the 8 patients using the `FindVariableFeatures` function. We then found a set of shared anchors across patients by canonical correlation analysis, using the `FindIntegrationAnchors` function with `anchor.features = 5000` and `dims = 1:50`, and performed data integration across patients based on these pre-computed anchors using the `IntegrateData` function with `dims = 1:50`.

#### MNN Correct

We implemented MNN Correct [27] on the log-pc counts by calling the `mnnCorrect` function in the R package `batchelor` (version 1.8.1).

#### Conos

We implemented Conos [28] on the log-pc counts using the R package `conos` (version 1.5.0). We first identified the top 8000 highly variable genes separately for scRNAseq data from each of the 8 patients using the `FindVariableFeatures` function in the R package `Seurat` (version 4.1.1) (so that a sufficiently large number of genes can be used for the subsequent data integration step). We then constructed a Conos object using the `Conos$new` function, built the joint graph using the `buildGraph` function with `k = 15`, `k.self = 5`, and `n.odgenes = 5000`, and computed batch corrected gene expression data using the `correctGenes` function with `n.od.genes = 8000`.

We assessed the performance of Liger using the returned matrix `H.norm`, the nonnegative normalized cell loading estimates. For Seurat 3, MNN Correct and Conos, we applied NMF with *K* = 12 to the batch corrected log-pc counts returned by each method, and assessed their performance using the estimated program membership matrix ***L*** from NMF. Arguments of all functions called to implement these methods were kept at their default values unless otherwise specified above.

### S.1.5. Tumor scRNA-seq datasets

#### HNSCC data

The HNSCC dataset was generated using SMART-Seq2, and contained transcriptional profiles for 5,902 cells from 18 patients [10]. Cells were confidently classified as malignant or normal cells using several complementary approaches in this study. Our GBCD analysis focused on tumors with at least 50 sequenced malignant cells, which involved 2,176 malignant cells from 10 primary HNSCC tumors and 5 matched lymph node metastases from these patients. We removed genes which were expressed in fewer than 20 cells, mitochondrial genes, and ribosomal genes, leaving a total of 17,113 genes. Since SMART-Seq2 does not yield UMI counts, we directly used the normalized expression data from [10], which are obtained as log_2_(TPM_*i,j*_*/*10 + 1) where TPM_*i,j*_ is the transcript-per-million value for gene *j* in cell *i*.

#### PDAC data

The three PDAC datasets come from recently published scRNA-seq studies on PDAC. Details about these datasets are provided in Table 2. For all cohorts, we excluded neuroendocrine tumors and analyzed the tumors which were confirmed to be PDAC by histology, and focused on analyzing malignant cells only. For cohorts B and C, cells were annotated as malignant or normal in the respective studies [46, 47]. For cohort A, such information was not available, so we distinguished tumor cells from normal cells in the surrounding tumor microenvironment based on transcriptional profiles. In particular, we clustered cells in cohort A by their gene expression patterns using the functions `FindNeighbors` and `FindClusters` from the R package `Seurat` (version 4.1.1), and determined the identity of each cell cluster using known marker genes. We also performed copy number variation (CNV) analysis on cohort A using inferCNV of the Trinity CTAT Project at https://github.com/broadinstitute/inferCNV, and found that the cells classified as malignant based on expression patterns exhibited significantly higher CNV levels, lending evidence to our identification of malignant cells. We further filtered out low-quality cells with a large fraction of mitochondrial counts (*>* 10%) and genes which were expressed in fewer than 25 cells in any cohort, as well as mitochondrial genes and ribosomal genes, leaving a total of 13,844 genes measured in 35,670 cells. More specifically, with the filtering procedures described above, we focused our analysis on 18,920, 10,936 and 5,814 high-quality malignant cells respectively in cohort A, B and C, which initially contained 31,195, 41,986, and 23,042 cells respectively. We performed library size normalization and log-transformation on the UMI counts, as described in Sec. S.1.1.1.

### S.1.6. Gene-set enrichment analysis

For each identified GEP in HNSCC and PDAC analyses, we performed gene-set enrichment analysis of the most strongly up-regulated genes, which we defined as having a fold change estimate of at least 1.5 and being among the top 2% ranked genes based on fold change estimates. We reported the significantly enriched gene sets from MSigDB Hallmark and CP:Reactome gene sets, which had a Bonferroni-corrected *p*-value *<* 0.05 from Fisher’s exact test.

### S.1.7. Measuring concordance between gene expression signatures

To assess the degree of overlap between an expression signature from the literature and a given GEP *k* identified by GBCD, we performed the Wilcoxon rank sum test to compare the distributions of *f*_*jk*_ between (a) the top 100 ranked genes of this gene signature and (b) a matched control gene-set of random genes (matched to have similar average expression levels as (a)). The matched control genes were selected following the strategy used in [10]: we grouped all *J* genes into 50 bins based on their average expression levels in all cells, then for each gene in (a), we randomly chose 100 genes from the same expression bin and added them to (b), so that (b) has a distribution of average expression levels comparable to that of (a) and is 100-fold larger. The concordance between this previously derived expression signature and our GEP *k* was then quantified by *−* log_10_(*p*-value), where the *p*-value is produced by an one-sided Wilcoxon rank sum test which tests whether the median of *f*_*jk*_ in (a) is smaller than that in (b).

### S.1.8. Classification of GBCD-identified GEPs

We determined whether a given GEP *k* is patient-specific or shared among multiple patients as follows. For patient *m*(= 1, …, *M*), we denote the proportion of cells whose membership in GEP *k* exceeds 0.05 (which can be considered active in this GEP) as prop_*m*_. Letting *m*^*′*^ = argmax_*m*=1,…,*M*_ prop_*m*_, we define the “patient-specificity score” *υ*_*k*_ for GEP *k* as

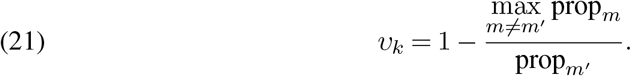

*υ*_*k*_ takes a value between 0 and 1. A *υ*_*k*_ close to 1 means that GEP *k* is predominantly expressed in patient *m*^*′*^, and a *υ*_*k*_ close to 0 means that another patient has a similar percentage of cells expressing GEP *k* compared to patient *m*^*′*^. In other words, *υ*_*k*_ reflects the degree of patient-specific expression of GEP *k*. GEP *k* is classified as “patient-specific” if *υ*_*k*_ *>* 0.5, and “shared” if *υ*_*k*_ *≤* 0.5.

For each shared GEP *k*, we performed one-way ANOVA on the GEP membership values ***l***_*k*_ with the patient identity of cells as the explanatory variable, and calculated the proportion of total variation explainable by patient identity. This proportion quantifies how strongly a shared GEP’s memberships vary between patients as opposed to within patients. In the results, we refer to this measure as “inter-patient variation”.

### S.1.9. Quantifying the spatial structure of GEP signatures

For each GEP *k* identified by GBCD, we studied whether the gene signature demonstrates systematic patterns across chromosome locations, in the sense that genes located closer to one another in chromosomal base-pair position tend to show more similar *f*_*jk*_ than genes located farther away. To do so, we fit a nonparametric regression model with *f*_*jk*_ as the response and the base-pair position as the predictor to all genes *j*, separately on each autosomal chromosome, *l* = 1, …, 22, using the function `gam` from the R package `mgcv` [101], then we recorded the model deviance. We quantified the degree of spatial structure for a given GEP *k* as

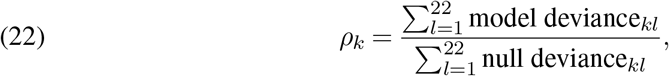

where the “null deviance” is based on a nonparametric regression model with a constant intercept term as the only predictor. A larger *ρ*_*k*_ indicates that a higher proportion of observed variation in *f*_*jk*_ can be explained by the chromosomal location of genes, which suggests greater spatial structure. In the results, we refer to *ρ*_*k*_ as the “chromosomal score” for GEP *k*.

### S.1.10. Survival analysis on PDAC bulk RNA-seq data

PDAC bulk RNA-seq data from [52, 56, 58] were downloaded from cBioPortal [102, 103] along with clinical annotations including overall survival information (i.e., time to death or right censoring). PDAC data from [57] were provided by Ontario Institute for Cancer Research based on a data sharing agreement with University of Chicago. Only surgically resected primary tumors that were histologically confirmed to be PDAC with no evidence of metastasis at diagnosis were included in the survival analysis, yielding a total of 391 patients (TCGA [56], *n* = 102; CPTAC [58], *n* = 87; QCMG [52], *n* = 70; PanCuRx [57], *n* = 132). Gene expression levels are quantified by log_2_-transformed TPM values.

For each of GEPs 1-14 and literature-derived subtype signatures, we computed an expression score in each tumor by comparing the average expression level of the top 200 ranked genes of this gene signature relative to a control gene-set [10], using the `Seurat` function `AddModuleScore`, and then, separately in each study, normalized expression scores across all tumors in that study to have zero mean and unit standard deviation. Stepwise variable selection in a Cox proportional hazards regression model, with overall survival as the endpoint and age, sex, tumor stage and all the program expression scores as covariates, was performed using the R package `My.stepwise` in [59]. Among the covariates, age, sex, and tumor stage are always included in the model, whereas the program expression scores are selected by the stepwise variable selection procedure.

### S.1.11. Optimization of the GBCD pipeline

As described in the **Computing requirements** section, our original GBCD implementation works with the (non-sparse) *N × N* matrix ***Y Y*** ^*T*^, which was used to generate all the results presented in the paper. However, this approach scales poorly with *N*, and therefore is unsuitable for large scRNA-seq data sets. We improved the GBCD implementation in the R package `gbcd`, which instead works directly with the (sparse) *N ×J* matrix ***Y*** and achieves significant reductions in runtime and memory usage for large *N*.

To verify that the two approaches produced similar results, we ran GBCD on the PDAC data, which is a large and highly complex dataset and so should allow us to detect sensitivity of GBCD results to any subtle changes in the computational pipeline. Without explicitly computing ***Y Y*** ^*T*^, the GBCD model fitting completed in 21 hours and required 48 GB; when explicitly computing ***Y Y*** ^*T*^, the GBCD model fitting completed in 50 hours and required 200 GB of memory. While there were some small differences in the results obtained, overall the two sets of results were very consistent (Supplementary Fig. S17).

The most time-consuming steps in fitting GBCD are the steps (ii) and (iv) which involve running *T*_1_ and *T*_2_ backfit iterations, respectively (Sec. S.1.1.2). We also assessed the impact of *T*_1_ and *T*_2_ on GBCD results, again using the PDAC data (the results presented in the paper were produced with *T*_1_ = 500 and *T*_2_ = 200). Supplementary Fig. S17 shows the time spent on backfitting as a function of the number of iterations. The runtime per backfit iteration in step (iv) was much longer than that in step (ii). This was likely due to the fact that there were more factors to update in each iteration in step (iv) than in step (ii). While the choice of *T*_1_ strongly affected a few GEPs (e.g., GEP2), the estimates of all GEPs stabilized at a *T*_2_ setting of 25. While it is difficult to draw general conclusions, our experiments suggest that GBCD results are not sensitive to *T*_2_, and can be set to a small number such as 25 to reduce runtime when *N* is large. Using the improved implementation on the PDAC data (i.e., working with ***Y*** rather then ***Y Y*** ^*T*^), setting *T*_1_ = 500 and *T*_2_ = 25 reduced computing time from 21 to 14 hours and achieved essentially the same results as our original implementation (which required 50 hours).

## Computing environment

Computations with simulated and real datasets were performed in R 4.1.0 [104], linked to the OpenBLAS 0.3.13 optimized numerical libraries, on Linux machines (Scientific Linux 8) with Intel Xeon E5-2680v4 (“Broadwell”) processors. Fitting GBCD to a single simulation dataset which contained expression data for 3,200 cells and 10,000 genes) required about 30 min and 2 GB of memory.

## Data availability

Most of the single-cell and bulk RNA-sequencing data used in this study are publicly available and downloadable following the URLs provided in the papers that generated the corresponding datasets, except for the data produced by the PanCuRx Translational Research Initiative [45], which require controlled access through a data sharing agreement with the PanCuRx Data Access Committee at the Ontario Institute for Cancer Research. Data and results necessary to reproduce the figures in the paper are available online (see URLs).

## Code availability

The R package `gbcd` implementing the GBCD method and the scripts for performing the simulations and the analyses of the HNSCC and PDAC data are available online (see URLs).

