## Supplementary figures for "Dissecting tumor transcriptional heterogeneity from single-cell RNA-seq data by generalized binary covariance decomposition"

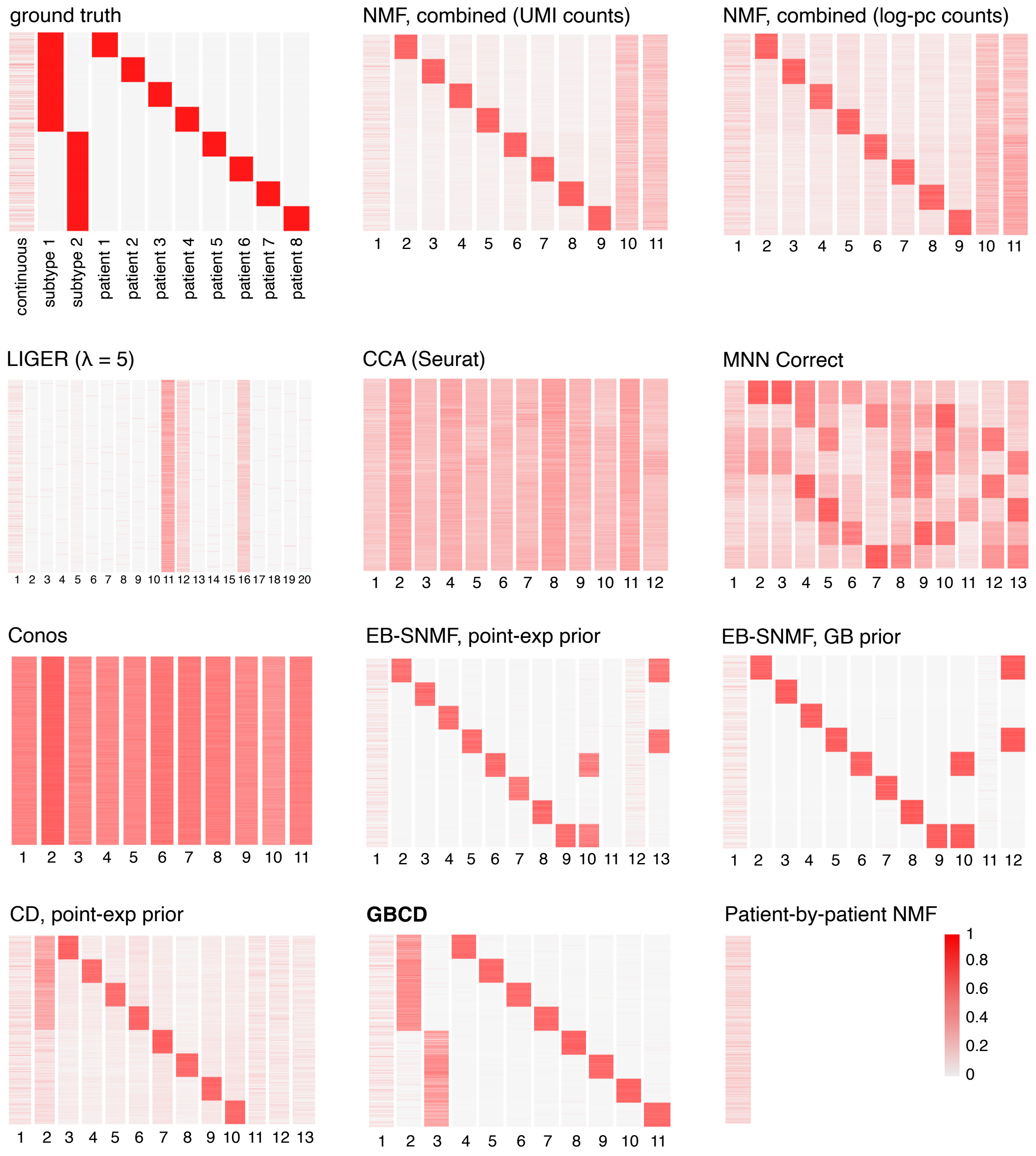


Supplementary Figure S1: GEP membership estimates for one of the 20 simulated data sets. In each heatmap, rows are cells, columns are GEPs. The columns were scaled separately so that the largest value in each column was always 1. Note that all methods identified one or more components that were strongly correlated with cellular detection rate; these components were not included in the heatmaps. Also note the patient-by-patient NMF identified only a single GEP.

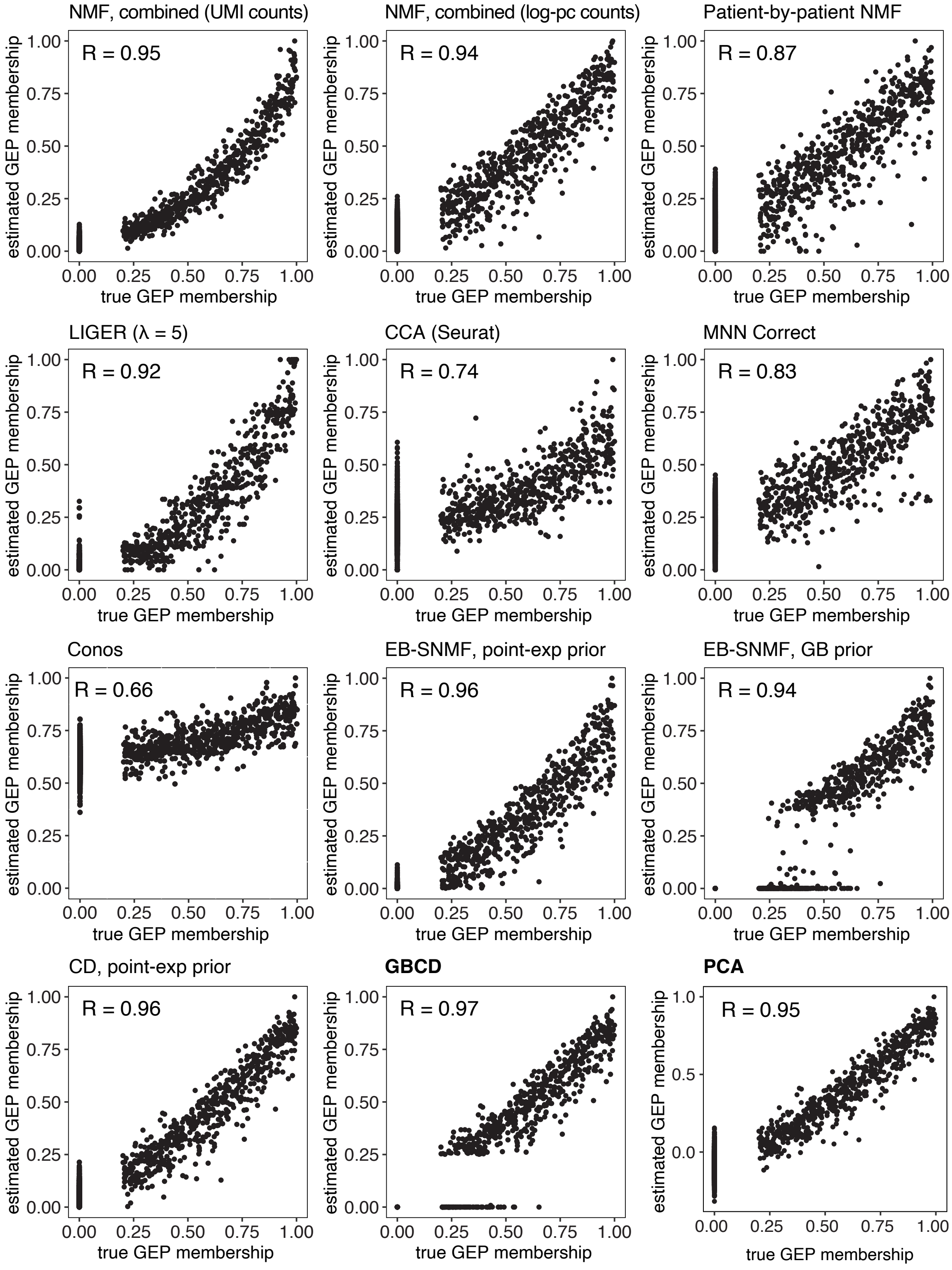


Supplementary Figure S2: GEP membership estimates from Supplementary Fig. S1 are shown in more detail for the continuous GEP only. The scatterplots compare the true and estimated memberships for each method. For each plot, the membership values are rescaled so that the maximum membership is always 1.

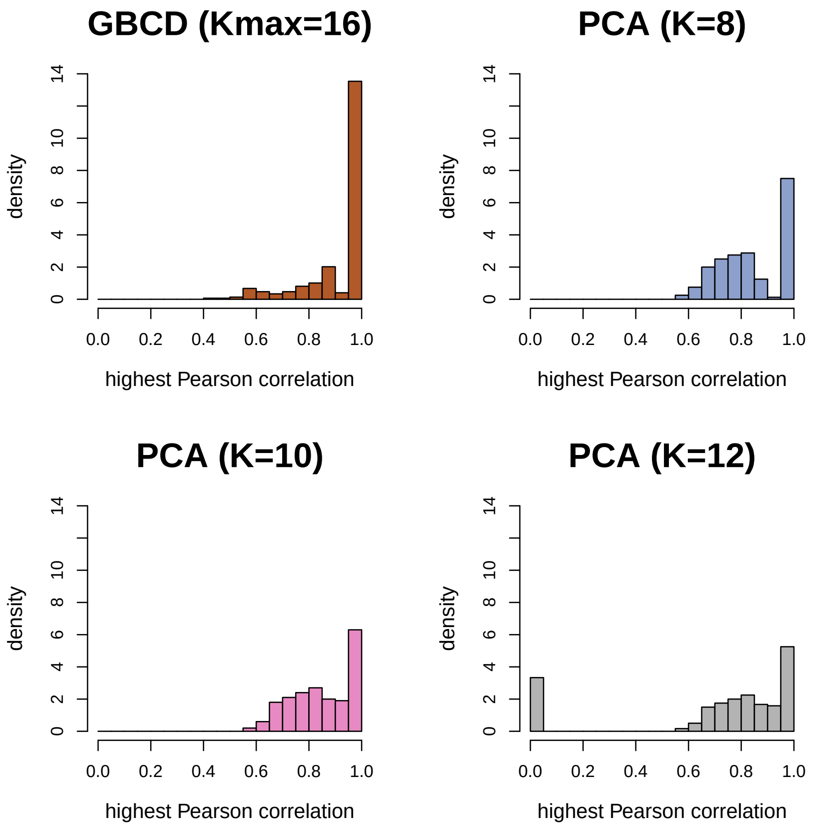


Supplementary Figure S3: Specificity of GBCD and PCA in recovering the true GEPs in the simulation study. For each GEP identified by a method, the highest Pearson correlation between its membership estimate and the true GEP membership among all 11 true GEPs is calculated, which measures how well this estimated GEP recovers a true GEP in the simulated data. The distribution of the highest Pearson correlation across all GEPs estimated from 20 simulated datasets is shown for GBCD and PCA with 8, 10 and 12 PCs.

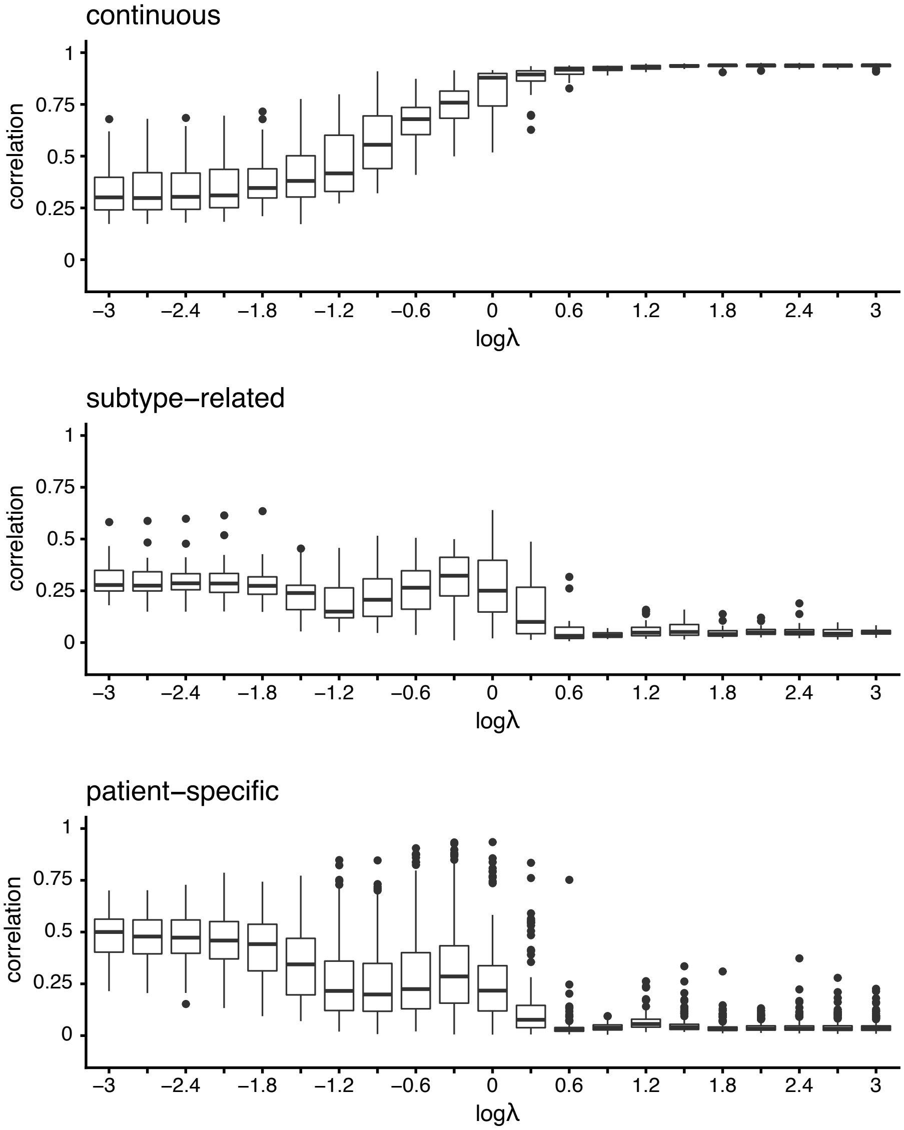


Supplementary Figure S4: Performance of LIGER in recovering the continuous, subtype-related and patient-specific GEPs across all 20 simulated datasets as the regularization parameter $\lambda$ was varied. For each true GEP, accuracy of the LIGER's estimate was measured using the highest Pearson correlation between the true GEP membership and the estimated membership among all GEPs identified by LIGER.

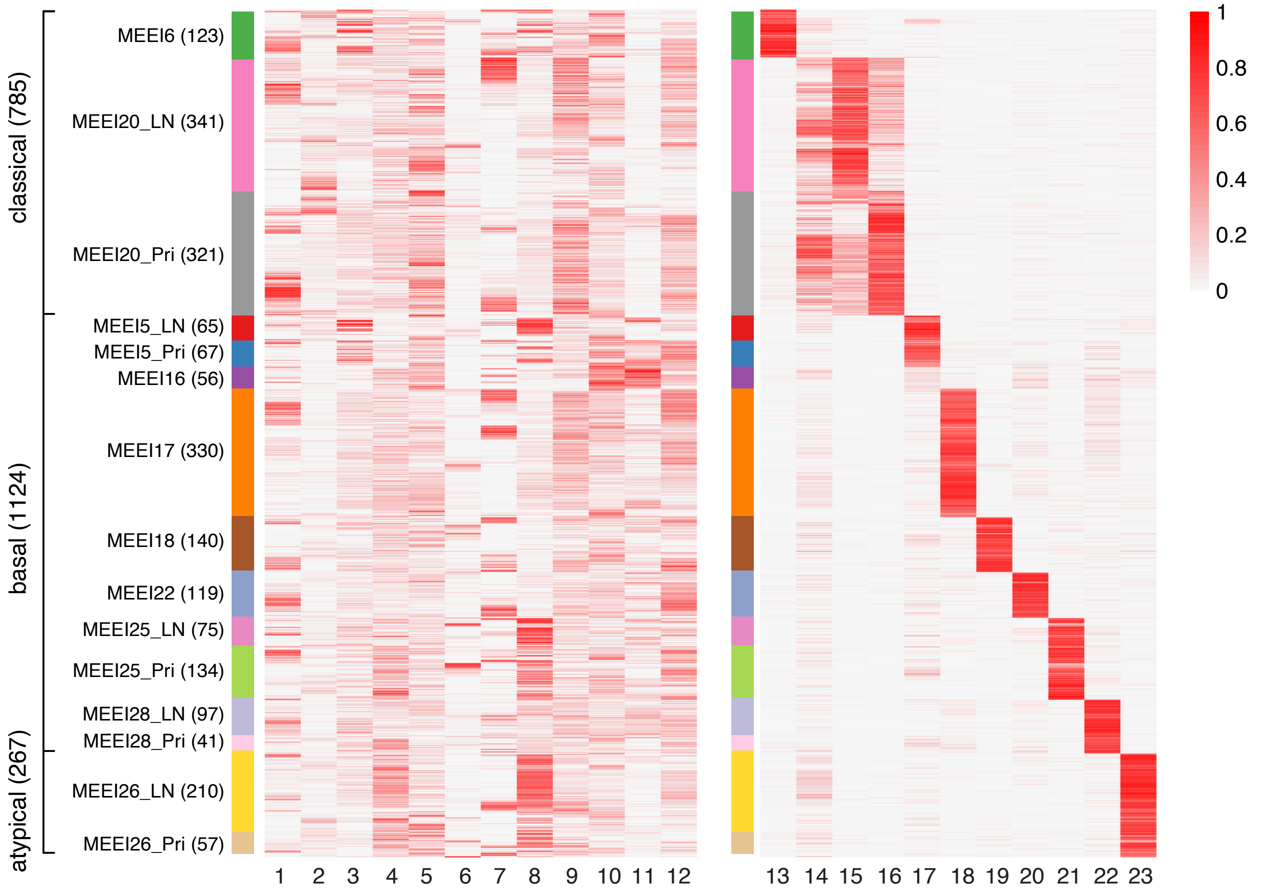


Supplementary Figure S5: Result of applying NMF to the HNSCC log-pc count data. The heatmap shows the membership estimates for the 2,176 cells (rows) and the 23 GEPs identified by NMF (columns). Cells are arranged top-to-bottom by tumor molecular subtype and patient, and GEPs are grouped left-to-right based on whether they are more patient-specific (GEP 13–23) or more shared across patients (GEP 1–12). For the heatmap, membership values were rescaled separately for each GEP so that the maximum membership for each GEP was always 1.

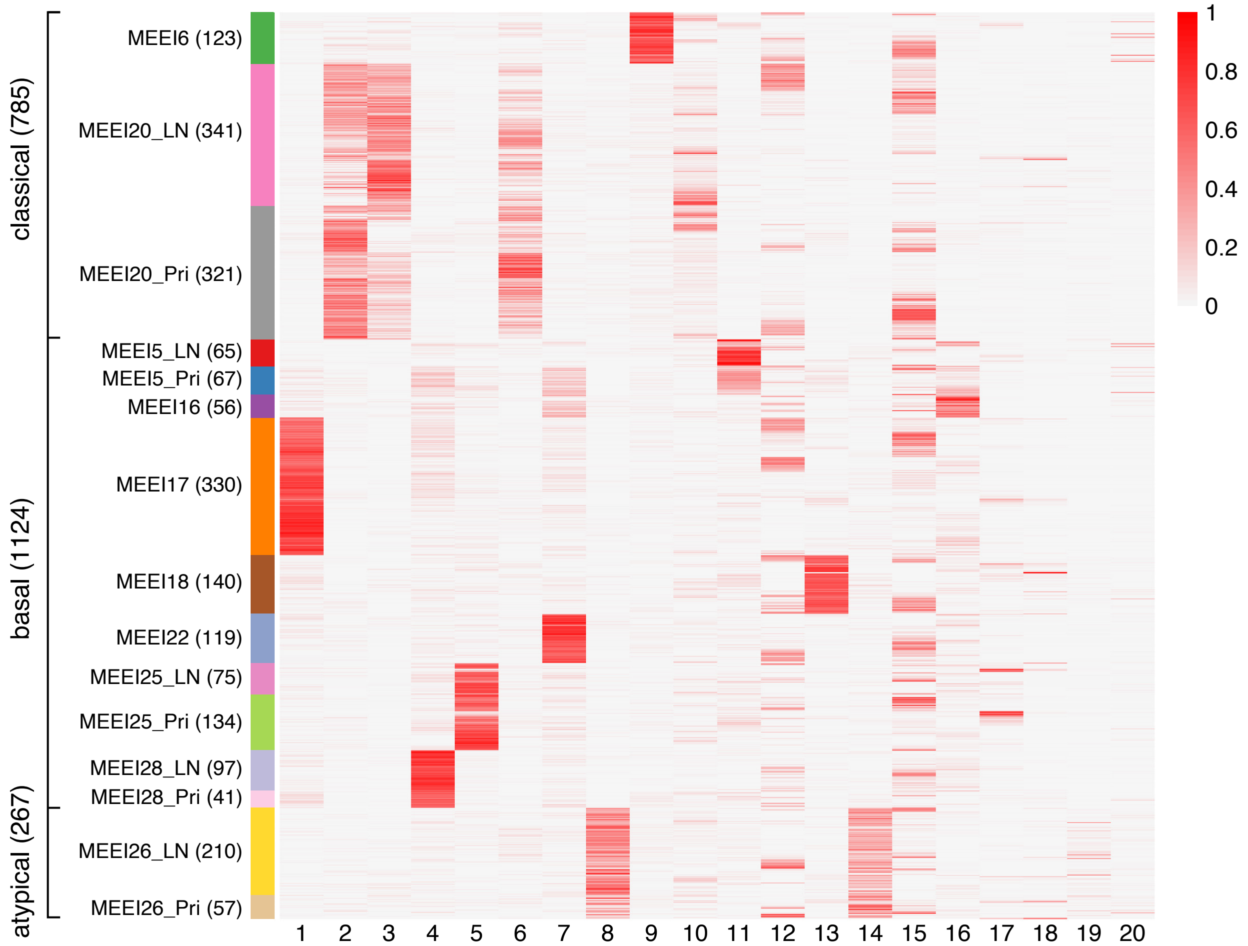


Supplementary Figure S6: Result of applying consensus NMF to the HNSCC log-pc count data. The heatmap shows the membership estimates for the 2,176 cells (rows) and the 20 GEPs identified by consensus NMF (columns). The number of GEPs was selected based on the trade-off between stability and error. Cells are arranged top-to-bottom by tumor molecular subtype and patient. For the heatmap, membership values were rescaled separately for each GEP so that the maximum membership for each GEP was always 1.

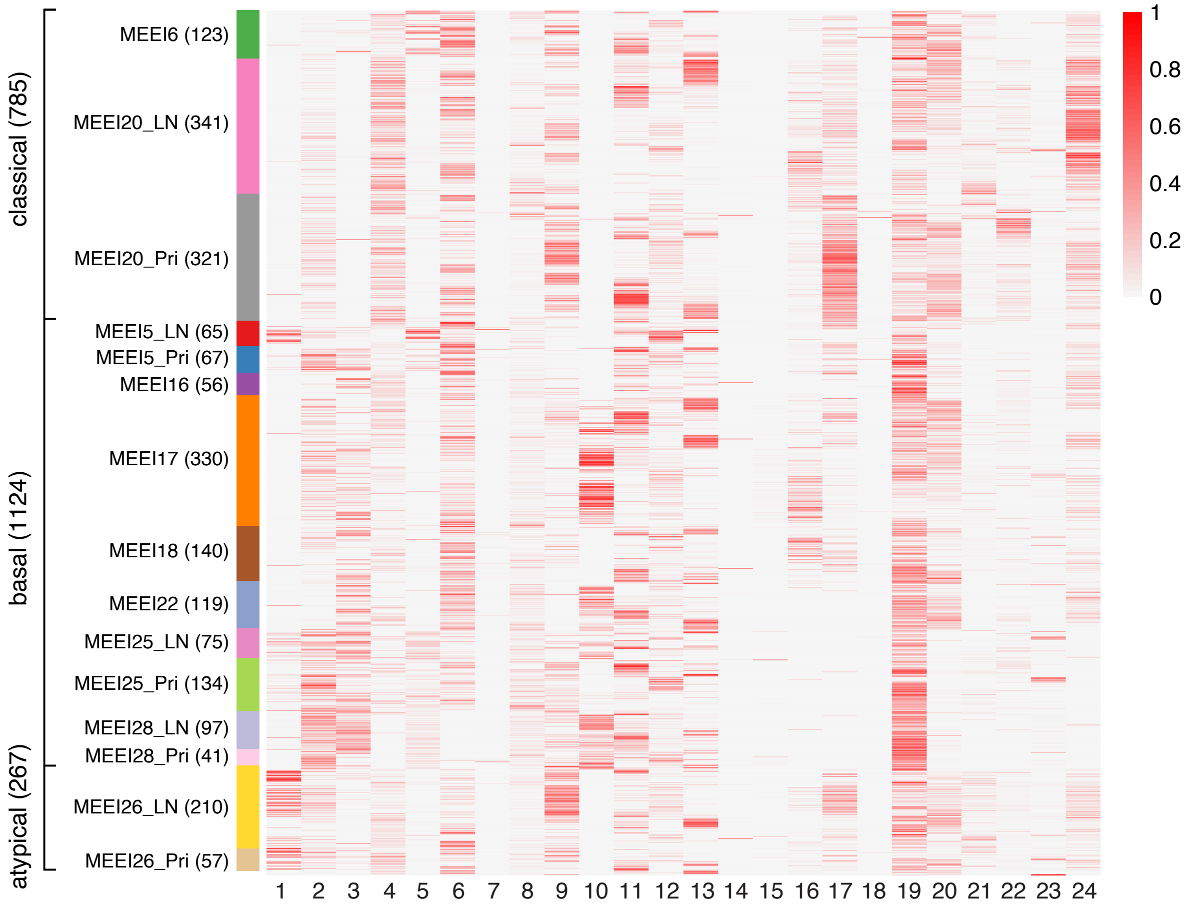


Supplementary Figure S7: Result of applying LIGER to the HNSCC log-pc count data. The heatmap shows the membership estimates for the 2,176 cells (rows) and the 24 GEPs identified by LIGER (columns). Cells are arranged top-to-bottom by tumor molecular subtype and patient. Membership values were rescaled separately for each GEP so that the maximum membership for each GEP was always 1.

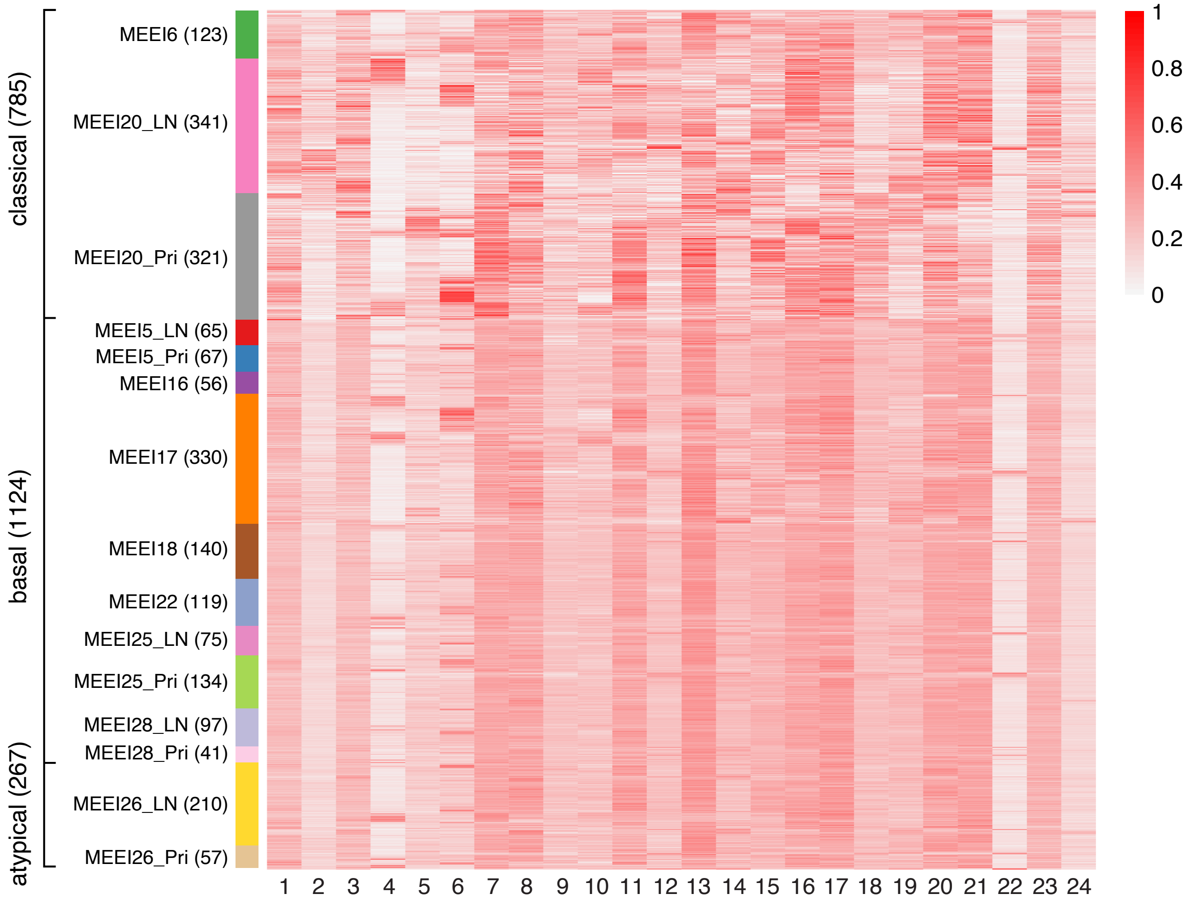


Supplementary Figure S8: Result of applying CCA (implemented in Seurat) to the HNSCC log-pc count data. The heatmap shows the membership estimates for the 2,176 cells (rows) and the 24 GEPs identified by CCA (columns). Cells are arranged top-to-bottom by tumor molecular subtype and patient. Membership values were rescaled separately for each GEP so that the maximum membership for each GEP was always 1.

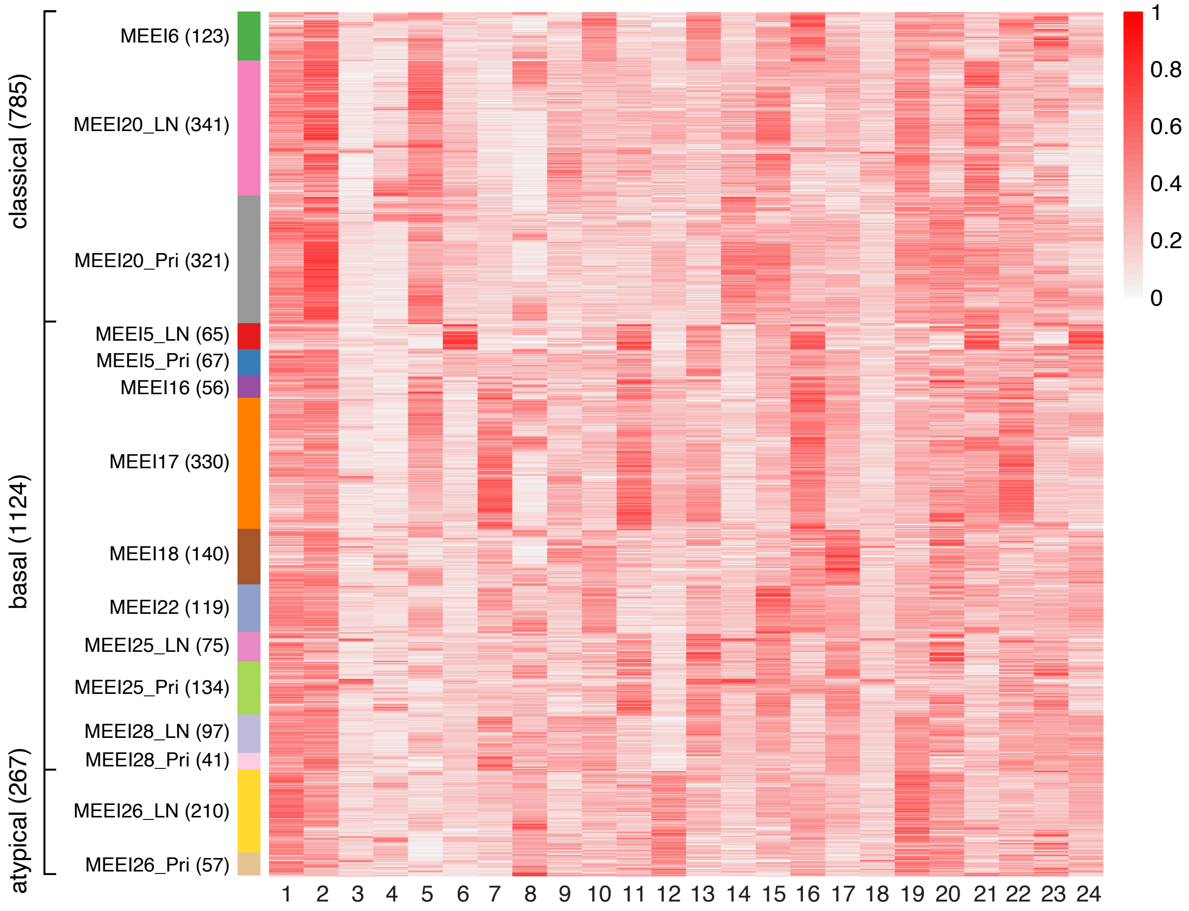


Supplementary Figure S9: Result of applying MNN Correct to the HNSCC log-pc count data. The heatmap shows the membership estimates for the 2,176 cells (rows) and the 24 GEPs identified by MNN Correct (columns). Cells are arranged top-to-bottom by tumor molecular subtype and patient. Membership values were rescaled separately for each GEP so that the maximum membership for each GEP was always 1.

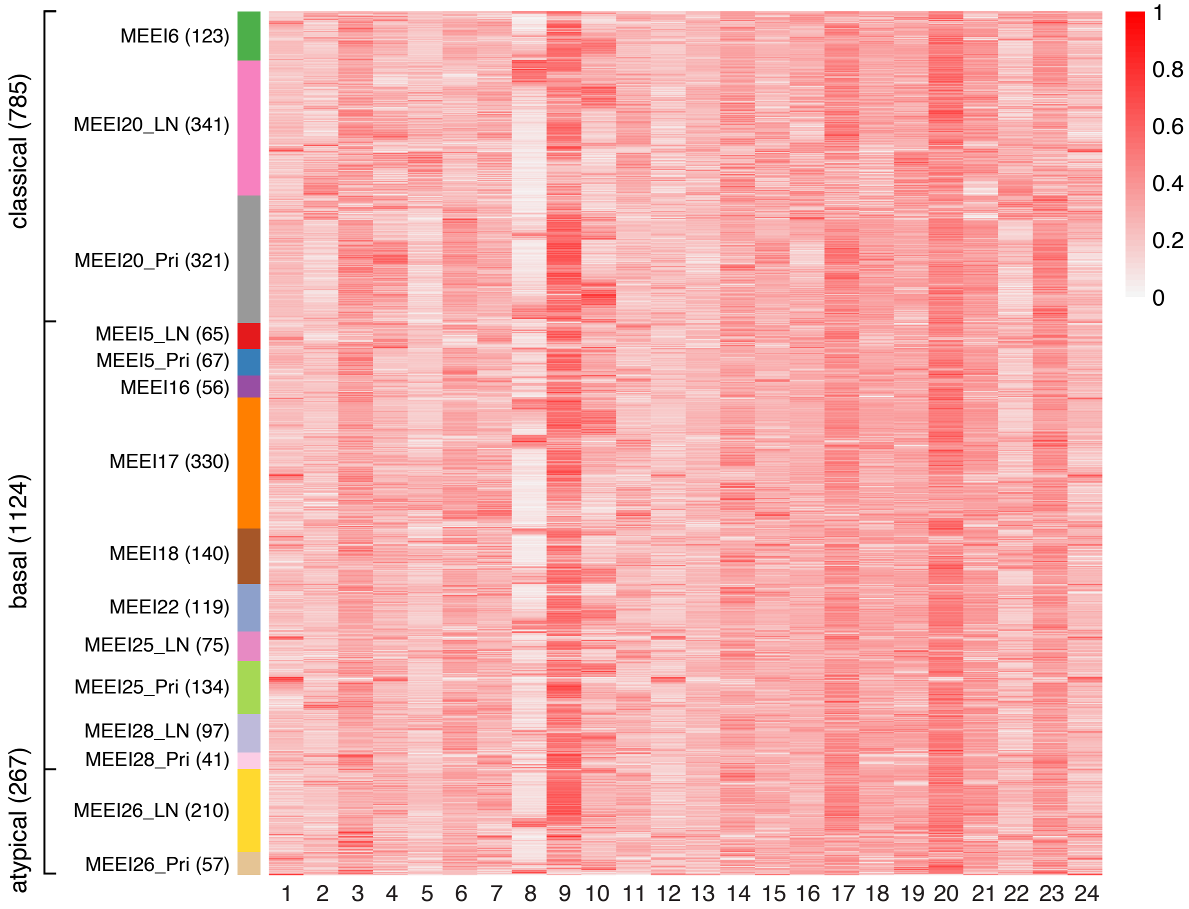


Supplementary Figure S10: Result of applying Conos to the HNSCC log-pc count data. The heatmap shows the membership estimates for the 2,176 cells (rows) and the 24 GEPs identified by Conos (columns). Cells are arranged top-to-bottom by tumor molecular subtype and patient. Membership values were rescaled separately for each GEP so that the maximum membership for each GEP was always 1.

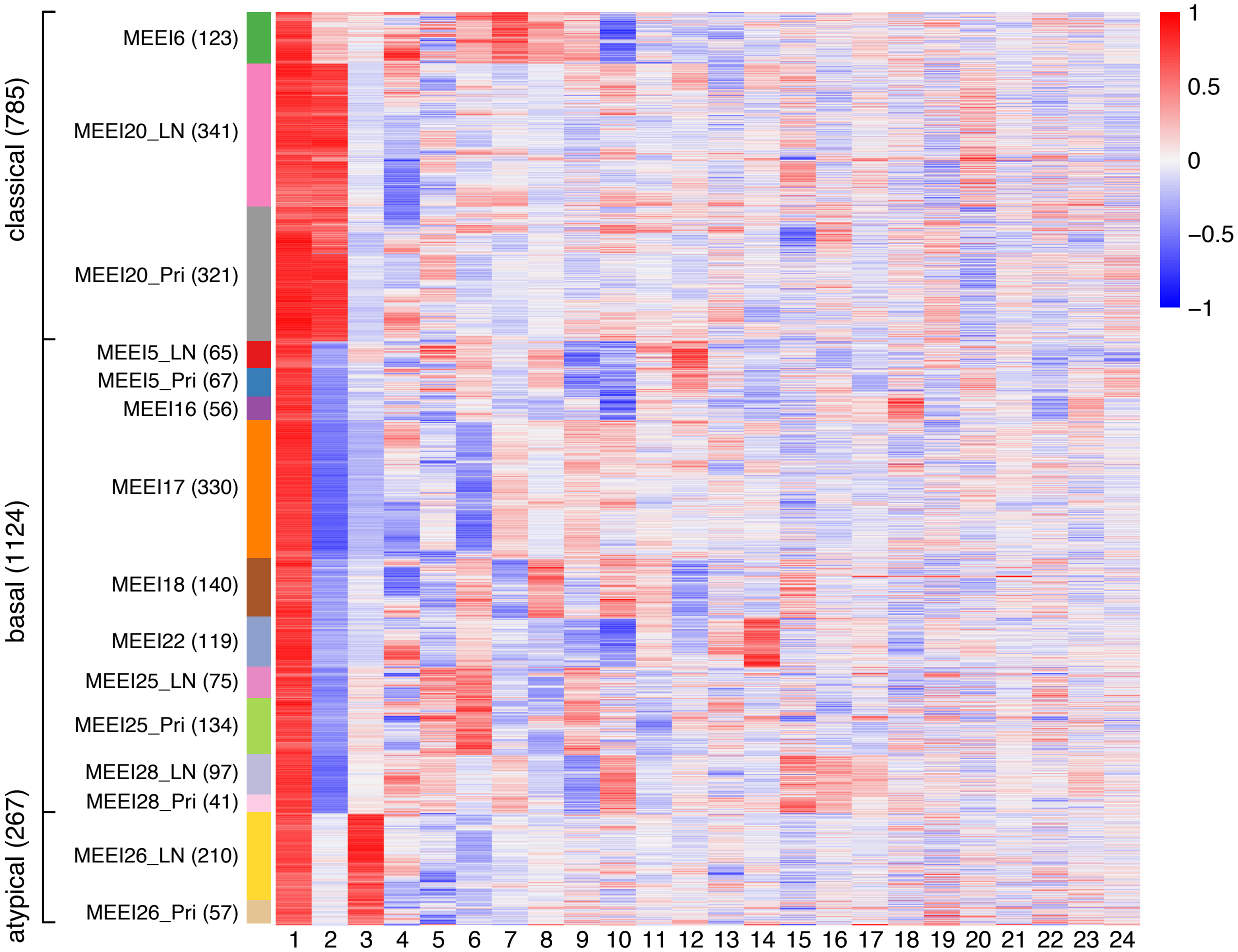


Supplementary Figure S11: Result of applying PCA to the HNSCC log-pc count data. The heatmap shows the principal component (PC) scores for the 2,176 cells (rows) and the 24 PCs. Cells are arranged top-to-bottom by tumor molecular subtype and patient. PC scores were rescaled separately for each PC so that the maximum PC score for each PC was always 1.

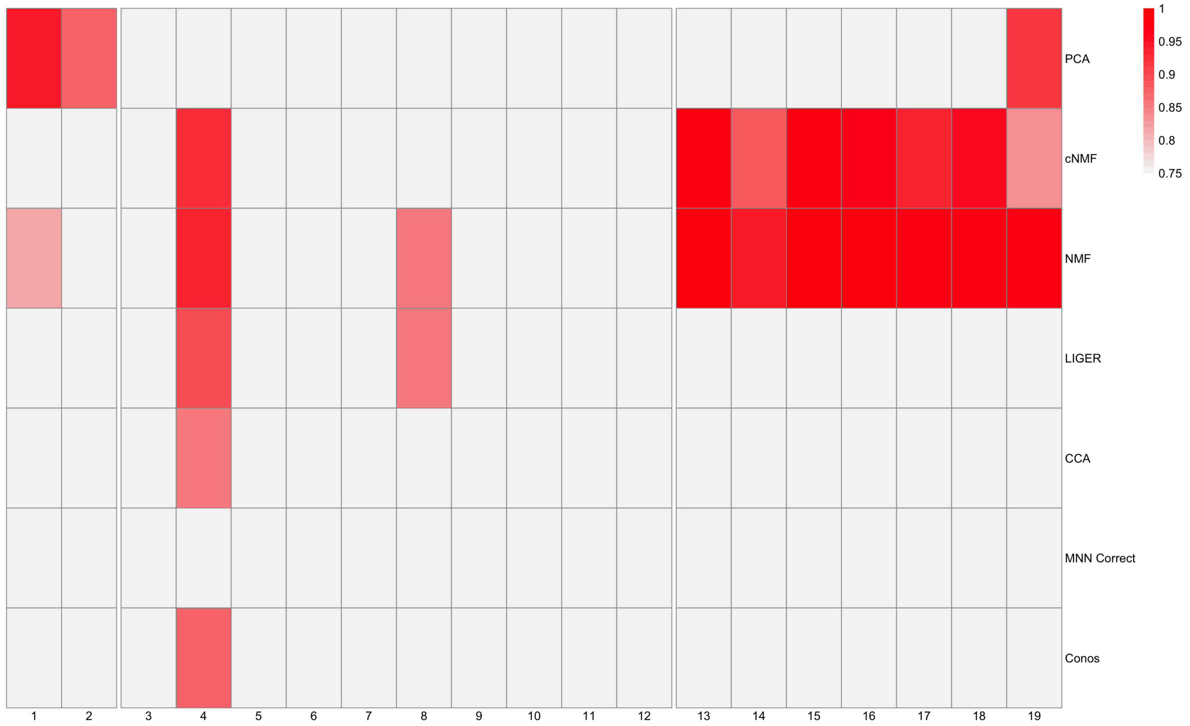


Supplementary Figure S12: Comparison of GEPs estimated by alternative methods to the 19 GEPs identified by the GBCD fit to the HNSCC data. For each of the 19 GBCD GEPs, consistency of an alternative method's estimate was measured using the highest Pearson correlation between the GBCD GEP membership and the estimated membership among all GEPs identified by the method.

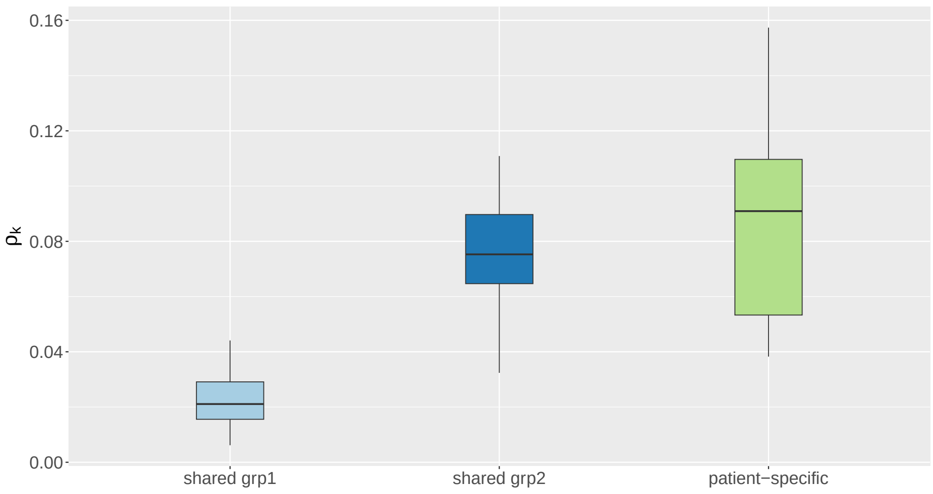


Supplementary Figure S13: Boxplots of $\rho_k$, which quantifies spatial structure of GEP signatures, for GEPs identified from the GBCD fit to the PDAC data, separately for GEP 1–14 ("shared grp1"), 16–20 ("shared grp2") and 21–33 ("patient-specific").

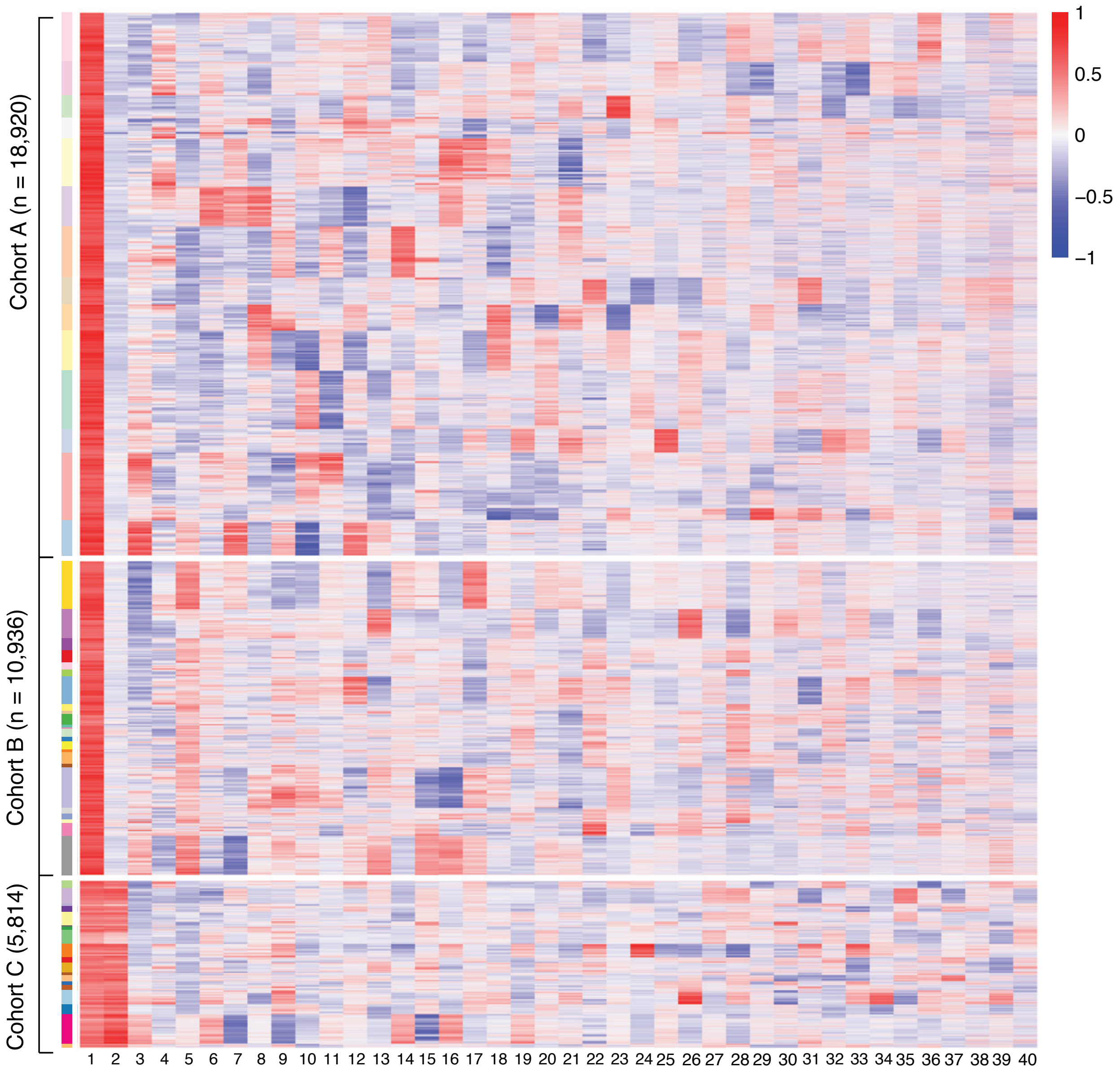


Supplementary Figure S14: Result of applying PCA to the PDAC log-pc count data. The heatmap shows the principal component (PC) scores for the 35,670 cells (rows) and the top 40 PCs (columns). Cells are arranged top-to-bottom by study and patient. Within each study, patients are ordered by the proportion of cells expressing GBCD GEP1, which strongly correlates with the classical subtype of PDAC. PC scores were rescaled separately for each PC so that the maximum PC score for each PC was always 1.

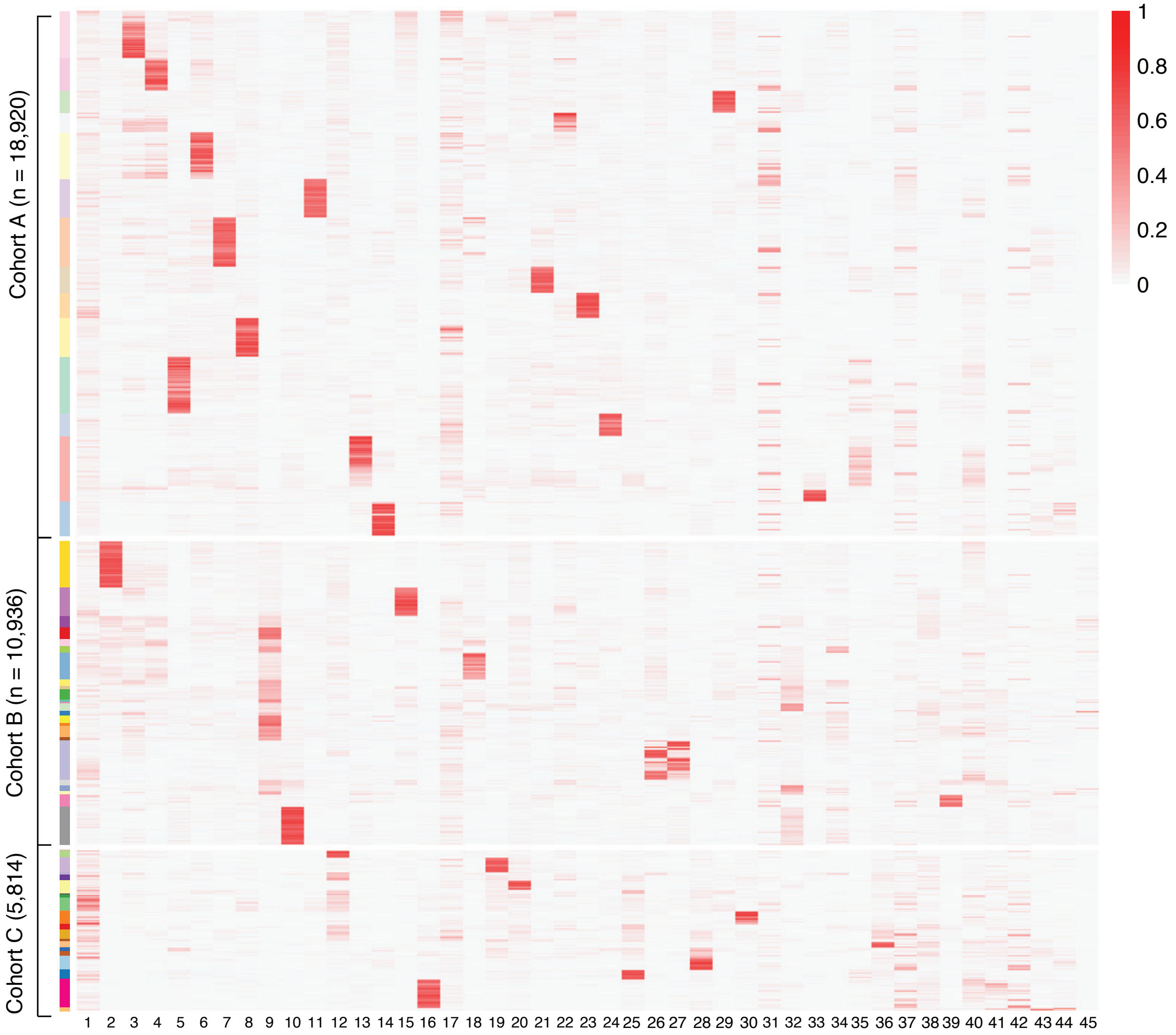


Supplementary Figure S15: Result of applying consensus NMF to the PDAC log-pc count data. The heatmap shows the membership estimates for the 35,670 cells (rows) and the 45 GEPs identified by consensus NMF (columns). The number of GEPs were selected based on the trade-off between stability and error. Cells are arranged top-to-bottom by study and patient. Within each study, patients are ordered by the proportion of cells expressing GBCD GEP1, which strongly correlates with the classical subtype of PDAC. For the heatmap, membership values were rescaled separately for each GEP so that the maximum membership for each GEP was always 1.

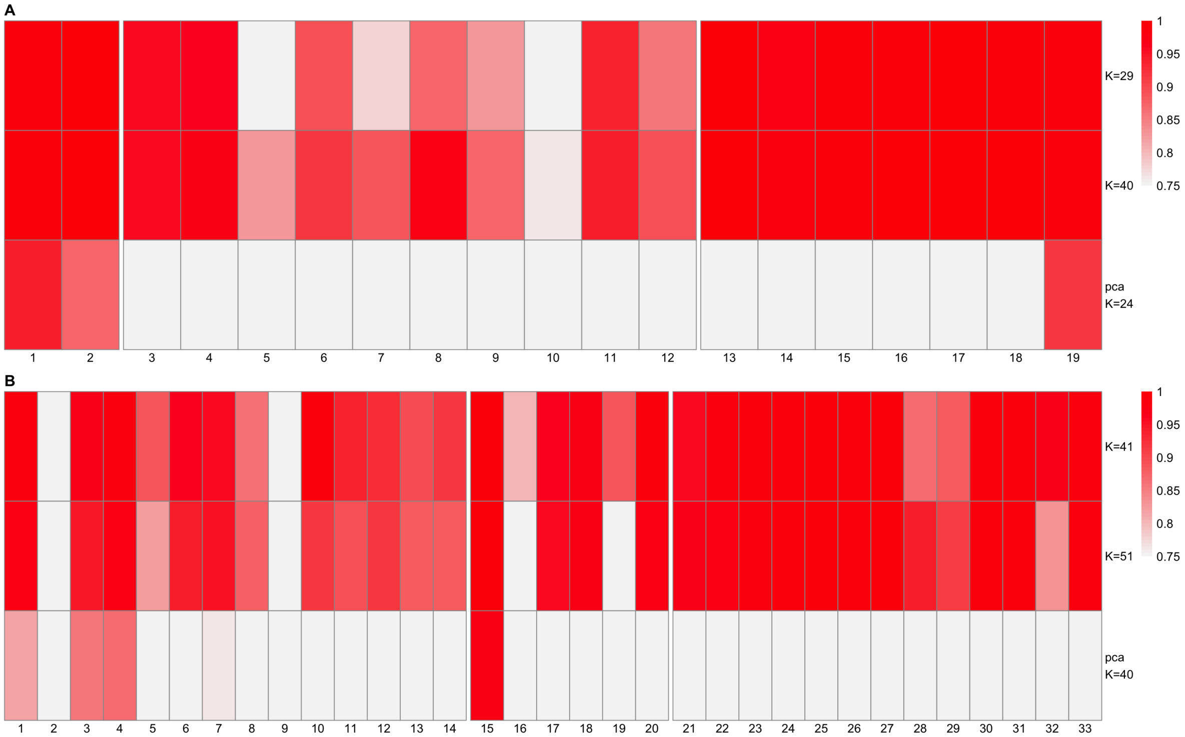


Supplementary Figure S16: Sensitivity analysis of GBCD results to $K$, the number of GEPs, in the HNSCC and PDAC data. The GBCD results analyzed in the main paper contained 19 GEPs for the HNSCC data and 33 GEPs for the PDAC data. For both data applications, GBCD was implemented with two larger values for $K$, and the results were compared to those reported in the paper. For each of the 19 HNSCC GEPs and 33 PDAC GEPs reported in the main paper, consistency of another GBCD fit with a larger $K$ was measured using the highest Pearson correlation between the membership of the given GEP and the estimated membership among all GEPs identified by another GBCD fit, which was shown in Panel A for HNSCC data and Panel B for PDAC data. Consistency of PCA (with 24 PCs for the HNSCC data and 40 PCs for the PDAC data) was similarly summarized.

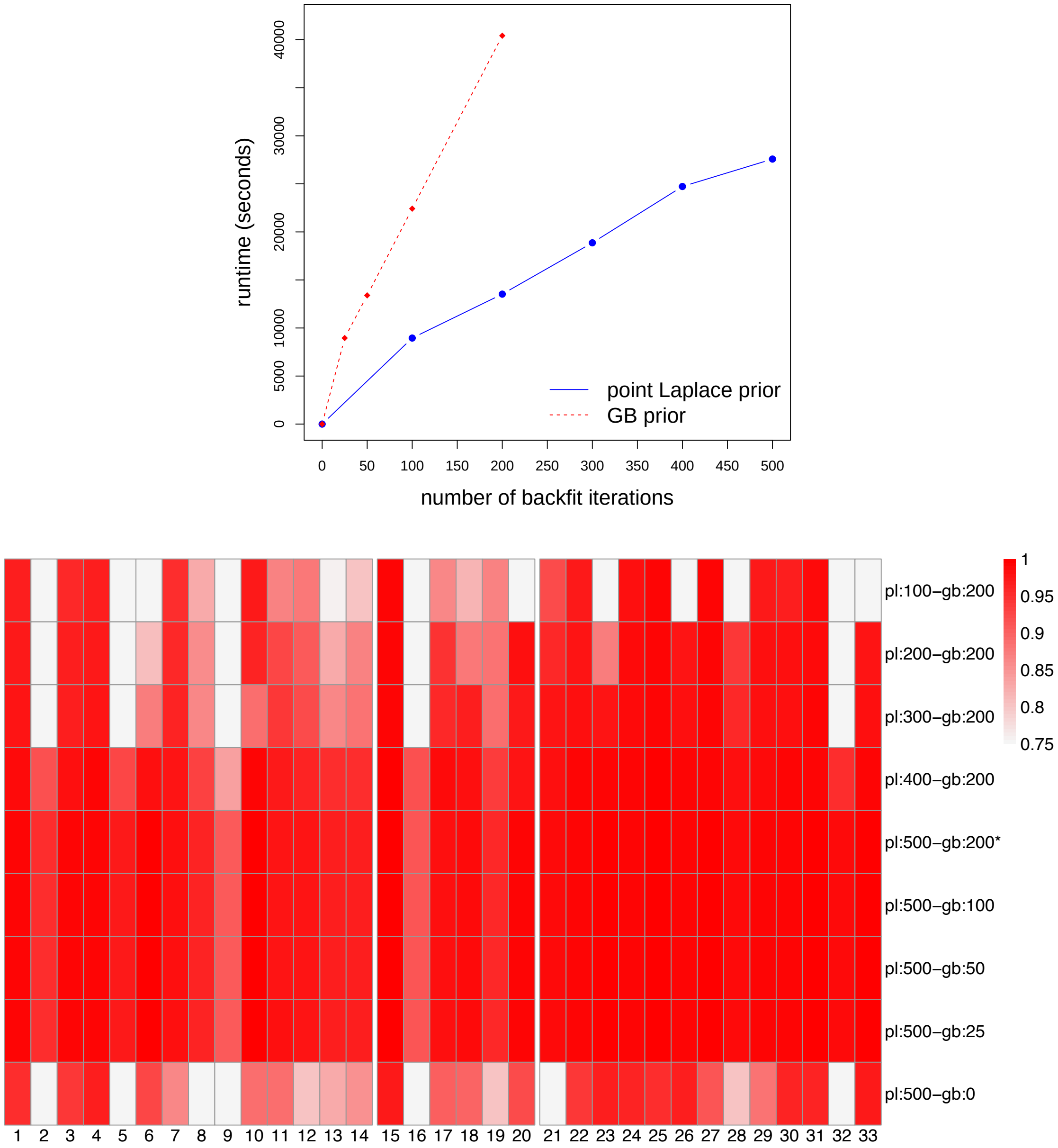


Supplementary Figure S17: GBCD implementation pipeline analysis using the PDAC data. The top panel displays the time spent on backfitting GEP membership matrix $\boldsymbol{L}$ at different stages (described in detail in Section S.1.11) as a function of the number of backfit iterations, using the improved implementation (i.e., working directly with $\boldsymbol{Y}$ rather than forming $\boldsymbol{Y}\boldsymbol{Y}^T$). The bottom panel compares the GEP membership estimates produced by the improved implementation with different numbers of backfit iterations $T_1$ and $T_2$ to those presented in the paper, which were produced using the original implementation (which forms $\boldsymbol{Y}\boldsymbol{Y}^T$ explicitly) with $T_1$=500 and $T_2$=200. More specifically, we examined how well each GEP reported in the paper was recovered by another GBCD fit using the highest Pearson correlation between the membership estimate of the reported GEP and the membership estimates among all GEPs identified by the other GBCD fit. The central row (marked with an asterisk) furnishes a *direct* comparison of the two implementations with $T_1$=500 and $T_2$=200, thus allowing for an assessment of the concordance of GBCD results between the two approaches (working with $\boldsymbol{Y}$ or $\boldsymbol{Y}\boldsymbol{Y}^T$). In theory, results should be identical; we suspect that the minor differences in GEPs 9 and 16 are due to numerical errors that are propagated over many backfit iterations.

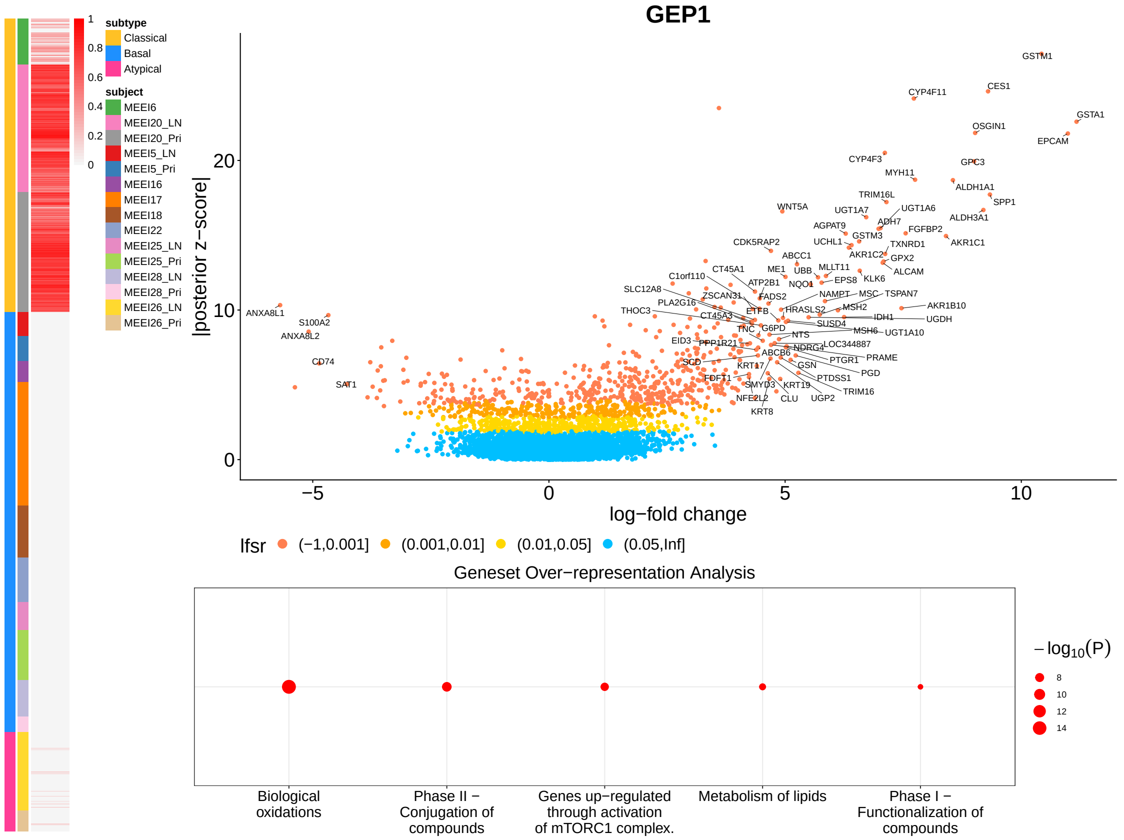
GEP1
subtype
Classical
Basal
Atypical
subject
MEEI6
MEEI20_LN
MEEI20_Pri
MEEI5_LN
MEEI5_Pri
MEEI16
MEEI17
MEEI18
MEEI22
MEEI25_LN
MEEI25_Pri
MEEI28_LN
MEEI28_Pri
MEEI26_LN
MEEI26_Pri
|posterior z−score|
log−fold change
lfsr (−1,0.001] (0.001,0.01] (0.01,0.05] (0.05,Inf]
GSTM1
CES1
CYP4F11
GSTA1
OSGIN1
EPCAM
CYP4F3
GPC3
MYH11
ALDH1A1
SPP1
TRIM16L
WNT5A
UGT1A7
UGT1A6
ALDH3A1
AGPAT9
ADH7
FGFBP2
GSTM3
AKR1C1
CDK5RAP2
UCHL1
TXNRD1
ABCC1
AKR1C2
GPX2
CT45A1
ME1
UBB
MLLT11
ALCAM
C1orf110
ATP2B1
NQO1
EPS8
KLK6
SLC12A8
ZSCAN31
FADS2
NAMPT
MSC
TSPAN7
PLA2G16
HRASLS2
MSH2
AKR1B10
THOC3
CT45A3
ETFB
IDH1
UGDH
TNC
G6PD
SUSD4
MSH6
UGT1A10
NTS
EID3
PPP1R21
NDRG4
LOC344887
SCD
ABCB6
PTGR1
PRAME
KRT17
GSN
PGD
FDFT1
PTDSS1
SMYD3
KRT19
TRIM16
NFE2L2
CLU
UGP2
KRT8
ANXA8L1
S100A2
ANXA8L2
CD74
SAT1
Geneset Over−representation Analysis
Biological oxidations
Phase II − Conjugation of compounds
Genes up−regulated through activation of mTORC1 complex.
Metabolism of lipids
Phase I − Functionalization of compounds
−log10(P)

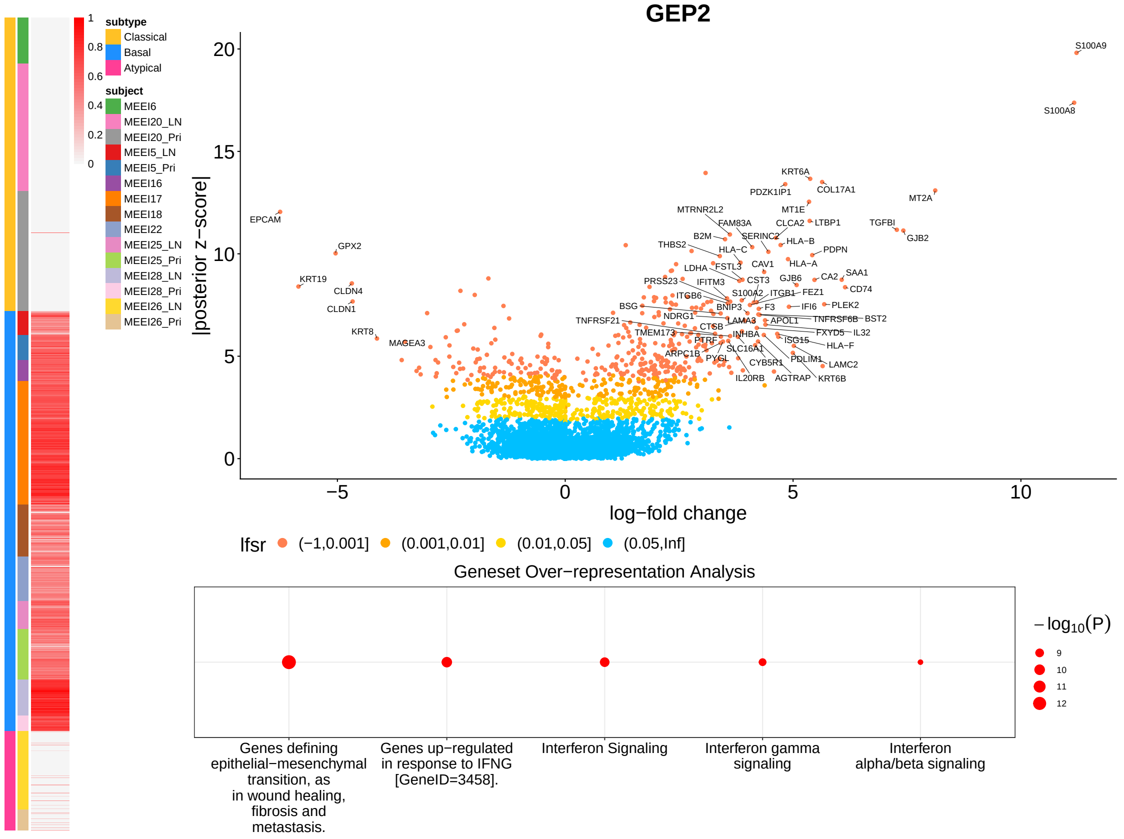
GEP2
subtype
Classical
Basal
Atypical
subject
MEEI6
MEEI20_LN
MEEI20_Pri
MEEI5_LN
MEEI5_Pri
MEEI16
MEEI17
MEEI18
MEEI22
MEEI25_LN
MEEI25_Pri
MEEI28_LN
MEEI28_Pri
MEEI26_LN
MEEI26_Pri
|posterior z−score|
log−fold change
lfsr (−1,0.001] (0.001,0.01] (0.01,0.05] (0.05,Inf]
S100A9
S100A8
KRT6A
PDZK1IP1
COL17A1
MT2A
MTRNR2L2
MT1E
EPCAM
FAM83A
CLCA2
LTBP1
TGFBI
GJB2
B2M
SERINC2
HLA−B
THBS2
HLA−C
PDPN
GPX2
LDHA
FSTL3
CAV1
HLA−A
PRSS23
IFITM3
CST3
GJB6
CA2
SAA1
KRT19
CLDN4
ITGB6
S100A2
ITGB1
FEZ1
CD74
CLDN1
BSG
BNIP3
F3
IFI6
PLEK2
TNFRSF21
NDRG1
LAMA3
APOL1
TNFRSF6B
BST2
KRT8
TMEM173
CTSB
FXYD5
IL32
MAGEA3
PTRF
INHBA
ISG15
HLA−F
ARPC1B
SLC16A1
PYGL
PDLIM1
LAMC2
CYB5R1
IL20RB
AGTRAP
KRT6B
Geneset Over−representation Analysis
Genes defining epithelial−mesenchymal transition, as in wound healing, fibrosis and metastasis.
Genes up−regulated in response to IFNG [GeneID=3458].
Interferon Signaling
Interferon gamma signaling
Interferon alpha/beta signaling
−log10(P)

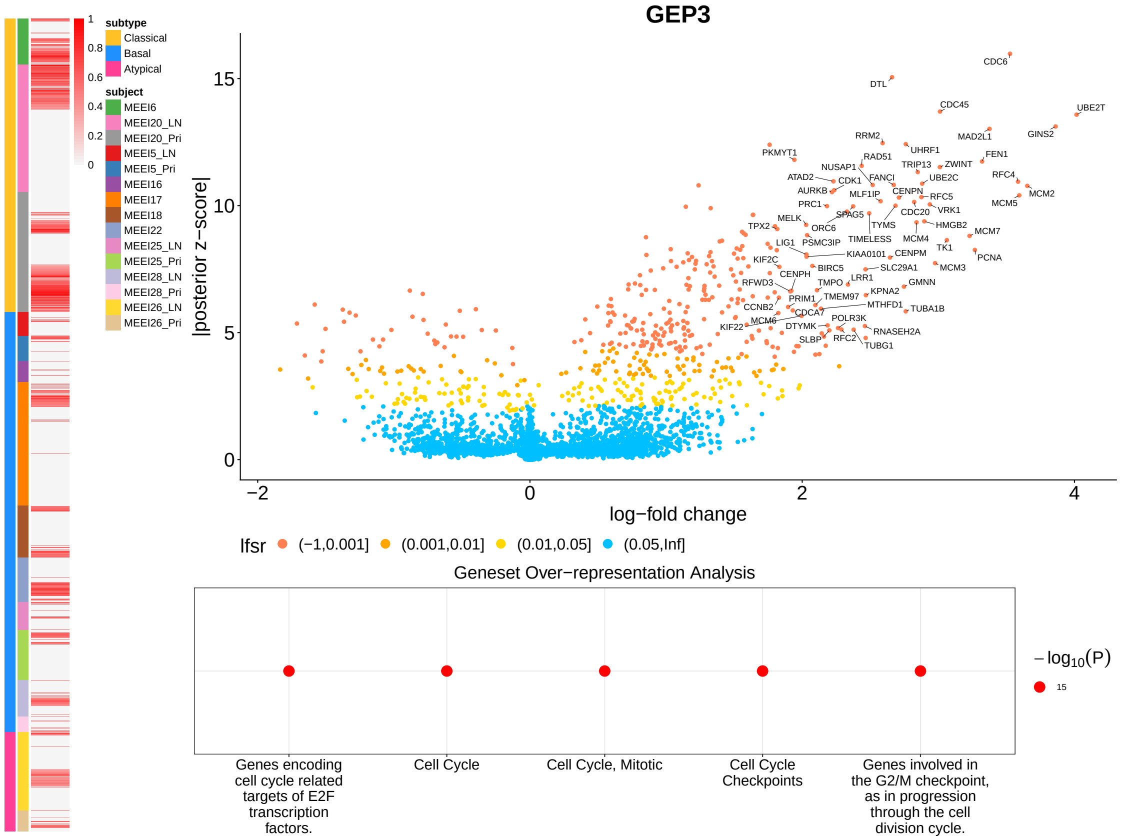

GEP3
subtype
Classical
Basal
Atypical
subject
MEEI6
MEEI20_LN
MEEI20_Pri
MEEI5_LN
MEEI5_Pri
MEEI16
MEEI17
MEEI18
MEEI22
MEEI25_LN
MEEI25_Pri
MEEI28_LN
MEEI28_Pri
MEEI26_LN
MEEI26_Pri
1
0.8
0.6
0.4
0.2
0
|posterior z−score|
log−fold change
CDC6
DTL
CDC45
UBE2T
MAD2L1
GINS2
RRM2
PKMYT1
RAD51
UHRF1
FEN1
NUSAP1
TRIP13
ZWINT
ATAD2
CDK1
FANCI
UBE2C
RFC4
AURKB
MLF1IP
CENPN
RFC5
MCM2
PRC1
CDC20
VRK1
MCM5
MELK
SPAG5
TYMS
HMGB2
TPX2
ORC6
MCM7
LIG1
PSMC3IP
TIMELESS
MCM4
TK1
KIAA0101
CENPM
PCNA
KIF2C
BIRC5
SLC29A1
MCM3
CENPH
LRR1
GMNN
RFWD3
TMPO
KPNA2
PRIM1
TMEM97
MTHFD1
TUBA1B
CCNB2
CDCA7
MCM6
POLR3K
KIF22
DTYMK
RNASEH2A
SLBP
RFC2
TUBG1
lfsr
(−1,0.001]
(0.001,0.01]
(0.01,0.05]
(0.05,Inf]
Geneset Over−representation Analysis
Genes encoding cell cycle related targets of E2F transcription factors.
Cell Cycle
Cell Cycle, Mitotic
Cell Cycle Checkpoints
Genes involved in the G2/M checkpoint, as in progression through the cell division cycle.
$-\log_{10}(P)$
15


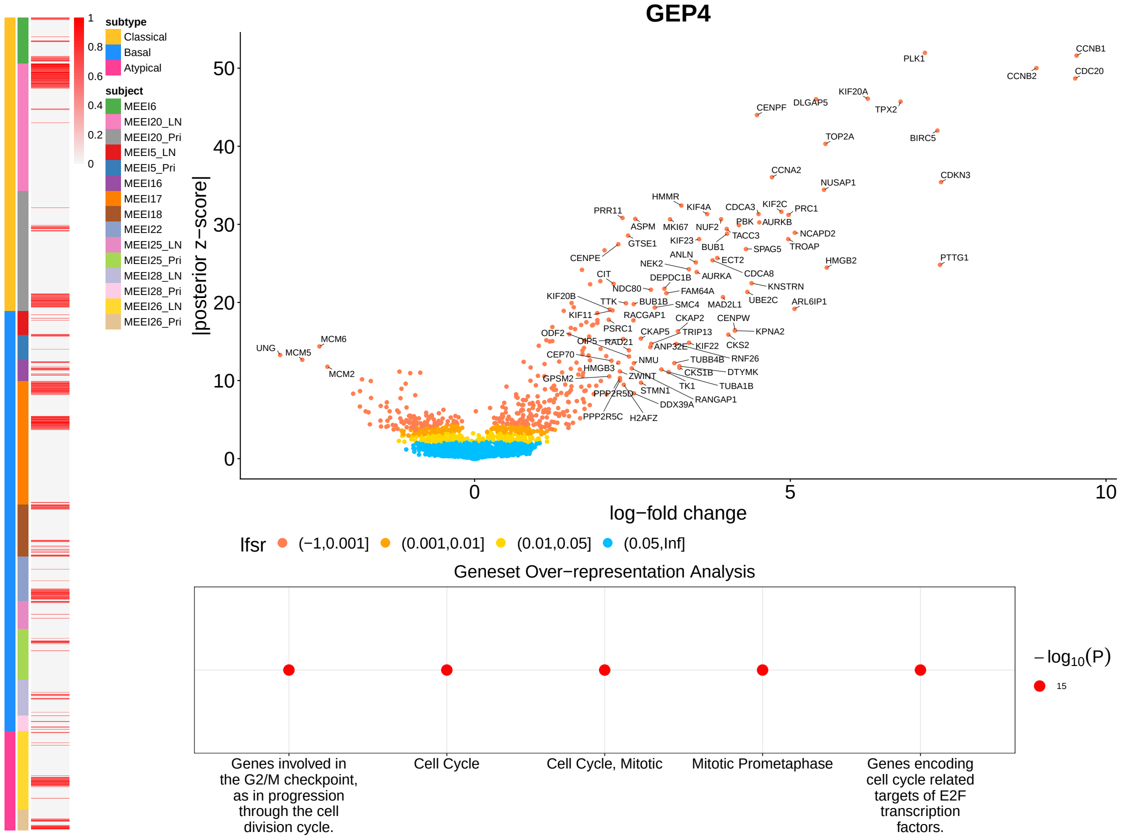

GEP4
subtype
Classical
Basal
Atypical
subject
MEEI6
MEEI20_LN
MEEI20_Pri
MEEI5_LN
MEEI5_Pri
MEEI16
MEEI17
MEEI18
MEEI22
MEEI25_LN
MEEI25_Pri
MEEI28_LN
MEEI28_Pri
MEEI26_LN
MEEI26_Pri
1
0.8
0.6
0.4
0.2
0
|posterior z−score|
log−fold change
CCNB1
PLK1
CCNB2
CDC20
KIF20A
DLGAP5
CENPF
TPX2
TOP2A
BIRC5
CCNA2
CDKN3
NUSAP1
HMMR
KIF4A
CDCA3
KIF2C
PRC1
PRR11
ASPM
MKI67
NUF2
PBK
AURKB
NCAPD2
GTSE1
KIF23
TACC3
BUB1
SPAG5
TROAP
CENPE
ANLN
ECT2
PTTG1
NEK2
HMGB2
CIT
DEPDC1B
AURKA
CDCA8
NDC80
KNSTRN
FAM64A
KIF20B
TTK
BUB1B
SMC4
MAD2L1
UBE2C
ARL6IP1
KIF11
RACGAP1
CKAP2
CENPW
PSRC1
CKAP5
TRIP13
KPNA2
ODF2
CIP5
RAD21
ANP32E
KIF22
CKS2
CEP70
NMU
TUBB4B
RNF26
HMGB3
CKS1B
DTYMK
GPSM2
ZWINT
TK1
TUBA1B
PPP2R5D
STMN1
RANGAP1
DDX39A
PPP2R5C
H2AFZ
UNG
MCM6
MCM5
MCM2
lfsr
(−1,0.001]
(0.001,0.01]
(0.01,0.05]
(0.05,Inf]
Geneset Over−representation Analysis
Genes involved in the G2/M checkpoint, as in progression through the cell division cycle.
Cell Cycle
Cell Cycle, Mitotic
Mitotic Prometaphase
Genes encoding cell cycle related targets of E2F transcription factors.
$-\log_{10}(P)$
15

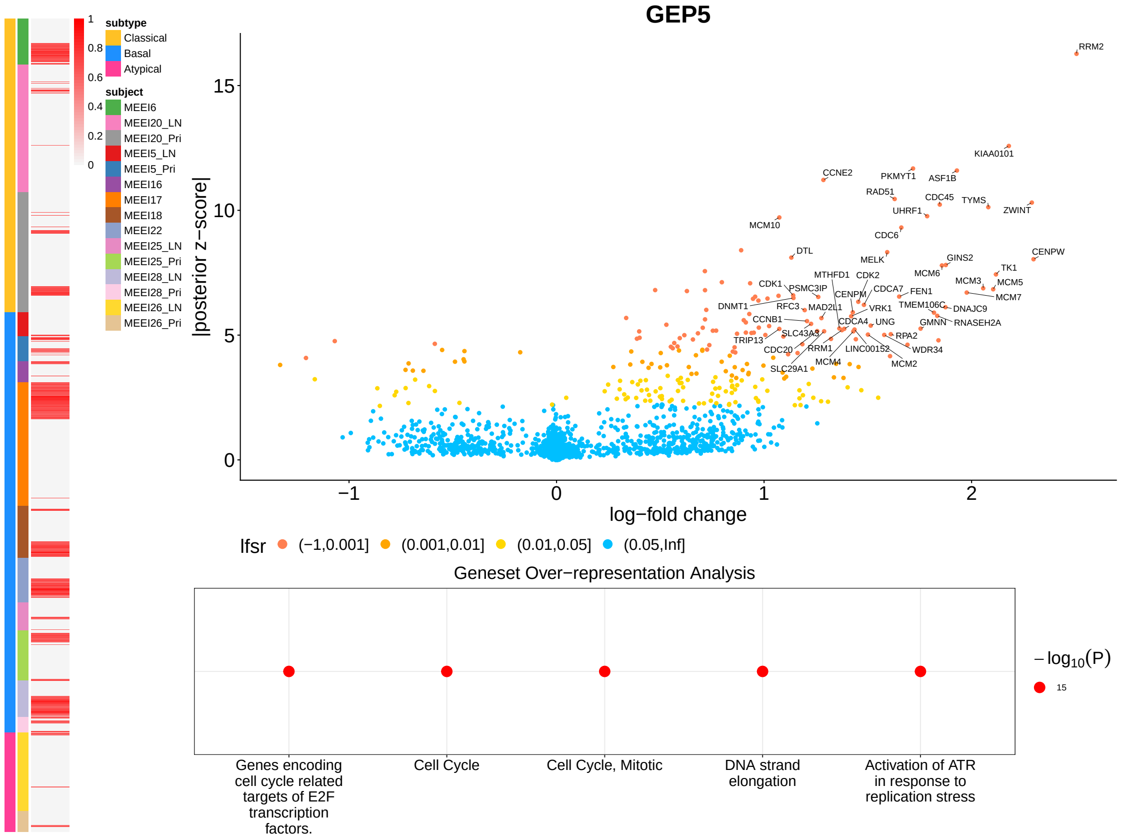

subtype
Classical
Basal
Atypical
subject
MEEI6
MEEI20_LN
MEEI20_Pri
MEEI5_LN
MEEI5_Pri
MEEI16
MEEI17
MEEI18
MEEI22
MEEI25_LN
MEEI25_Pri
MEEI28_LN
MEEI28_Pri
MEEI26_LN
MEEI26_Pri
1
0.8
0.6
0.4
0.2
0
GEP5
|posterior z−score|
log−fold change
RRM2
KIAA0101
CCNE2
PKMYT1
ASF1B
RAD51
CDC45
TYMS
ZWINT
UHRF1
MCM10
CDC6
DTL
MELK
GINS2
CENPW
TK1
MCM6
MCM3
MCM5
MCM7
CDK1
PSMC3IP
MTHFD1
CDK2
CENPM
CDCA7
FEN1
DNMT1
RFC3
MAD2L1
VRK1
TMEM106C
DNAJC9
CCNB1
CDCA4
UNG
GMNN
RNASEH2A
TRIP13
SLC43A3
RPA2
CDC20
RRM1
LINC00152
WDR34
MCM4
MCM2
SLC29A1
lfsr
(−1,0.001]
(0.001,0.01]
(0.01,0.05]
(0.05,Inf]
Geneset Over−representation Analysis
Genes encoding cell cycle related targets of E2F transcription factors.
Cell Cycle
Cell Cycle, Mitotic
DNA strand elongation
Activation of ATR in response to replication stress
$-\log_{10}(P)$
15


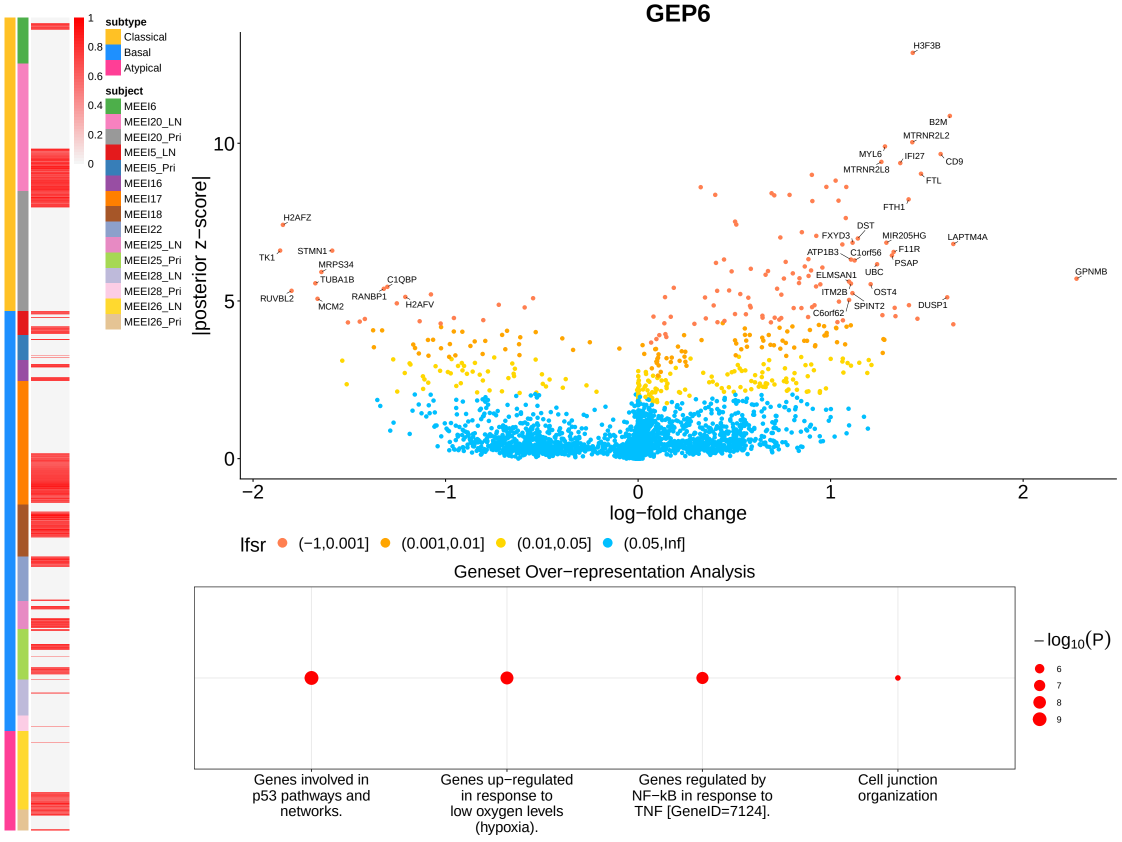

subtype
Classical
Basal
Atypical
subject
MEEI6
MEEI20_LN
MEEI20_Pri
MEEI5_LN
MEEI5_Pri
MEEI16
MEEI17
MEEI18
MEEI22
MEEI25_LN
MEEI25_Pri
MEEI28_LN
MEEI28_Pri
MEEI26_LN
MEEI26_Pri
GEP6
|posterior z−score|
log−fold change
H3F3B
B2M
MTRNR2L2
MYL6
IFI27
CD9
MTRNR2L8
FTL
FTH1
H2AFZ
DST
FXYD3
MIR205HG
LAPTM4A
F11R
TK1
STMN1
ATP1B3
C1orf56
PSAP
MRPS34
UBC
TUBA1B
C1QBP
ELMSAN1
GPNMB
ITM2B
OST4
RUVBL2
RANBP1
SPINT2
DUSP1
MCM2
H2AFV
C6orf62
lfsr
(−1,0.001]
(0.001,0.01]
(0.01,0.05]
(0.05,Inf]
Geneset Over−representation Analysis
Genes involved in p53 pathways and networks.
Genes up−regulated in response to low oxygen levels (hypoxia).
Genes regulated by NF−kB in response to TNF [GeneID=7124].
Cell junction organization
$-\log_{10}(P)$
6
7
8
9

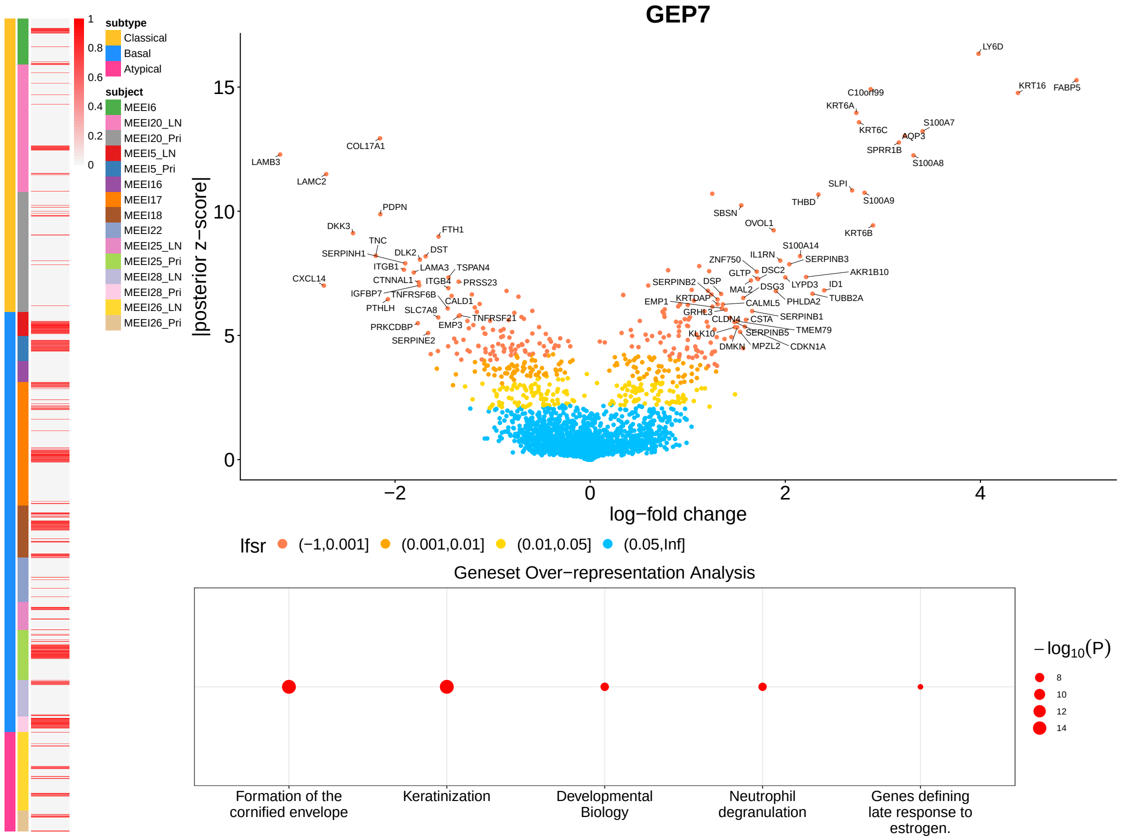
GEP7
subtype
Classical
Basal
Atypical
subject
MEEI6
MEEI20_LN
MEEI20_Pri
MEEI5_LN
MEEI5_Pri
MEEI16
MEEI17
MEEI18
MEEI22
MEEI25_LN
MEEI25_Pri
MEEI28_LN
MEEI28_Pri
MEEI26_LN
MEEI26_Pri
|posterior z−score|
log−fold change
lfsr (−1,0.001] (0.001,0.01] (0.01,0.05] (0.05,Inf]
LY6D
KRT16
FABP5
C10orf99
KRT6A
S100A7
KRT6C
AQP3
SPRR1B
S100A8
LAMB3
COL17A1
LAMC2
SLPI
THBD
S100A9
SBSN
OVOL1
PDPN
DKK3
FTH1
KRT6B
TNC
SERPINH1
DLK2
DST
ITGB1
LAMA3
TSPAN4
CXCL14
CTNNAL1
ITGB4
PRSS23
IGFBP7
TNFRSF6B
PTHLH
CALD1
SLC7A8
TNFRSF21
PRKCDBP
EMP3
SERPINE2
S100A14
IL1RN
ZNF750
SERPINB3
GLTP
DSC2
AKR1B10
SERPINB2
DSP
LYPD3
ID1
MAL2
DSG3
EMP1
KRTDAP
CALML5
PHLDA2
TUBB2A
GRHL3
CLDN4
CSTA
SERPINB1
KLK10
SERPINB5
TMEM79
DMKN
MPZL2
CDKN1A
Geneset Over−representation Analysis
Formation of the cornified envelope
Keratinization
Developmental Biology
Neutrophil degranulation
Genes defining late response to estrogen.
−log10(P)
8
10
12
14

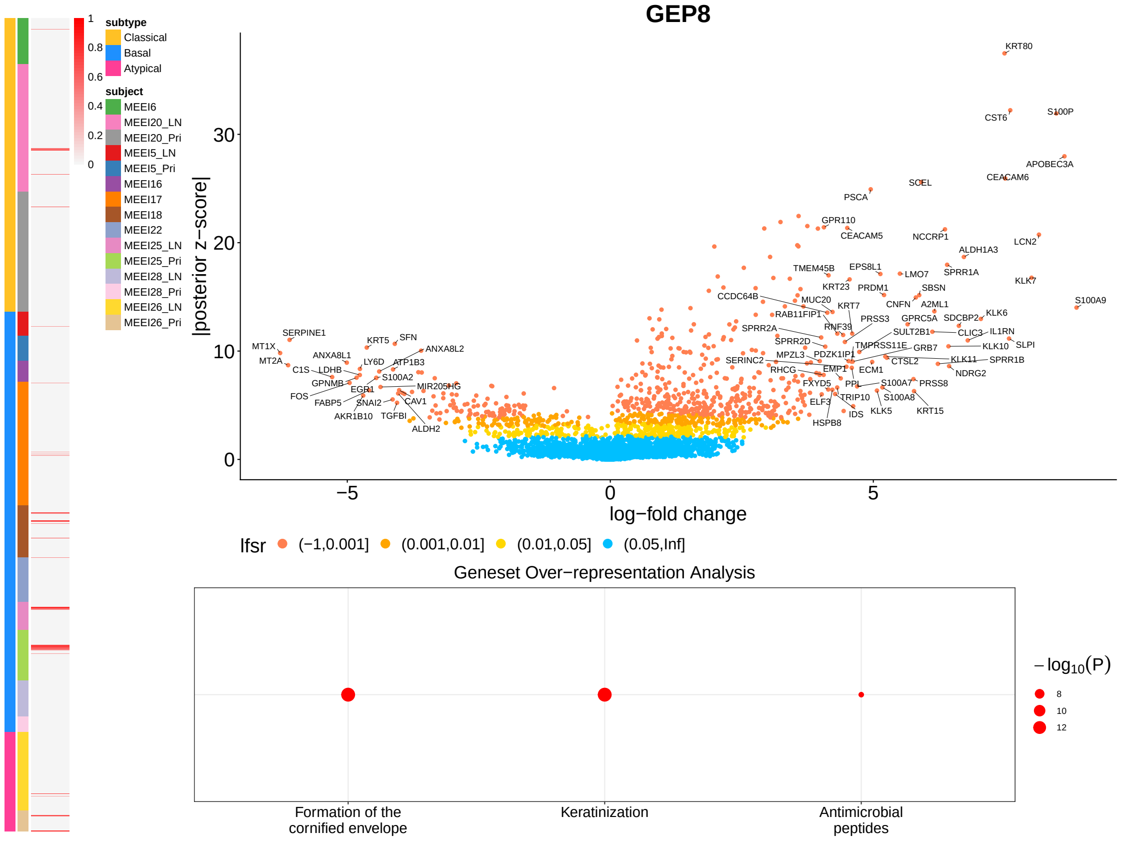
GEP8
subtype
Classical
Basal
Atypical
subject
MEEI6
MEEI20_LN
MEEI20_Pri
MEEI5_LN
MEEI5_Pri
MEEI16
MEEI17
MEEI18
MEEI22
MEEI25_LN
MEEI25_Pri
MEEI28_LN
MEEI28_Pri
MEEI26_LN
MEEI26_Pri
|posterior z−score|
log−fold change
lfsr (−1,0.001] (0.001,0.01] (0.01,0.05] (0.05,Inf]
KRT80
CST6
S100P
APOBEC3A
CEACAM6
SCEL
PSCA
GPR110
CEACAM5
NCCRP1
LCN2
ALDH1A3
TMEM45B
EPS8L1
LMO7
SPRR1A
KRT23
PRDM1
SBSN
KLK7
CCDC64B
MUC20
CNFN
A2ML1
S100A9
RAB11FIP1
KRT7
KLK6
PRSS3
GPRC5A
SDCBP2
SPRR2A
RNF39
SULT2B1
CLIC3
IL1RN
SPRR2D
TMPRSS11E
GRB7
KLK10
SLPI
SERPINE1
MT1X
KRT5
SFN
ANXA8L2
MT2A
ANXA8L1
LY6D
ATP1B3
C1S
LDHB
S100A2
GPNMB
EGR1
MIR205HG
FOS
FABP5
SNAI2
CAV1
AKR1B10
TGFBI
ALDH2
SERINC2
MPZL3
PDZK1IP1
CTSL2
KLK11
SPRR1B
RHCG
EMP1
ECM1
NDRG2
FXYD5
PPL
S100A7
PRSS8
ELF3
TRIP10
S100A8
IDS
KLK5
KRT15
HSPB8
Geneset Over−representation Analysis
Formation of the cornified envelope
Keratinization
Antimicrobial peptides
−log10(P)
8
10
12

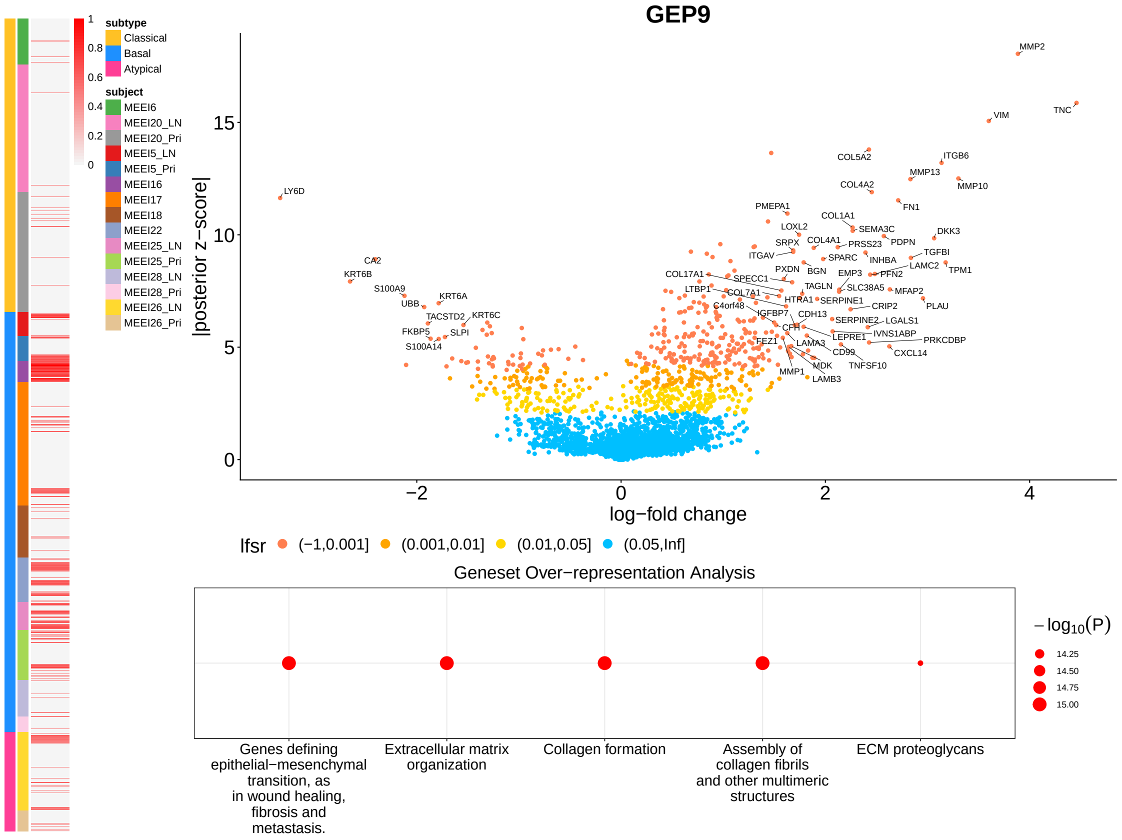
GEP9
log−fold change
|posterior z−score|
lfsr (−1,0.001] (0.001,0.01] (0.01,0.05] (0.05,Inf]
Geneset Over−representation Analysis
Genes defining epithelial−mesenchymal transition, as in wound healing, fibrosis and metastasis.
Extracellular matrix organization
Collagen formation
Assembly of collagen fibrils and other multimeric structures
ECM proteoglycans
subtype
Classical
Basal
Atypical
subject

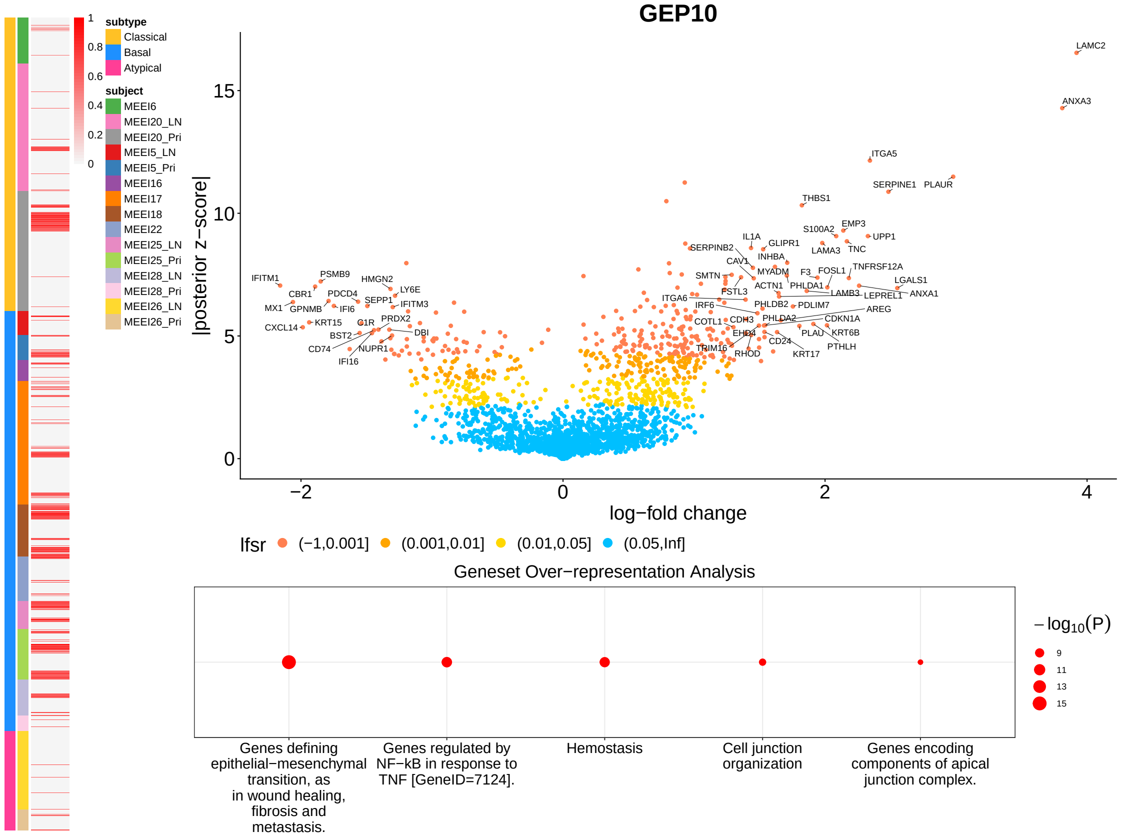
GEP10
log−fold change
|posterior z−score|
lfsr (−1,0.001] (0.001,0.01] (0.01,0.05] (0.05,Inf]
Geneset Over−representation Analysis
Genes defining epithelial−mesenchymal transition, as in wound healing, fibrosis and metastasis.
Genes regulated by NF−kB in response to TNF [GeneID=7124].
Hemostasis
Cell junction organization
Genes encoding components of apical junction complex.
subtype
Classical
Basal
Atypical
subject

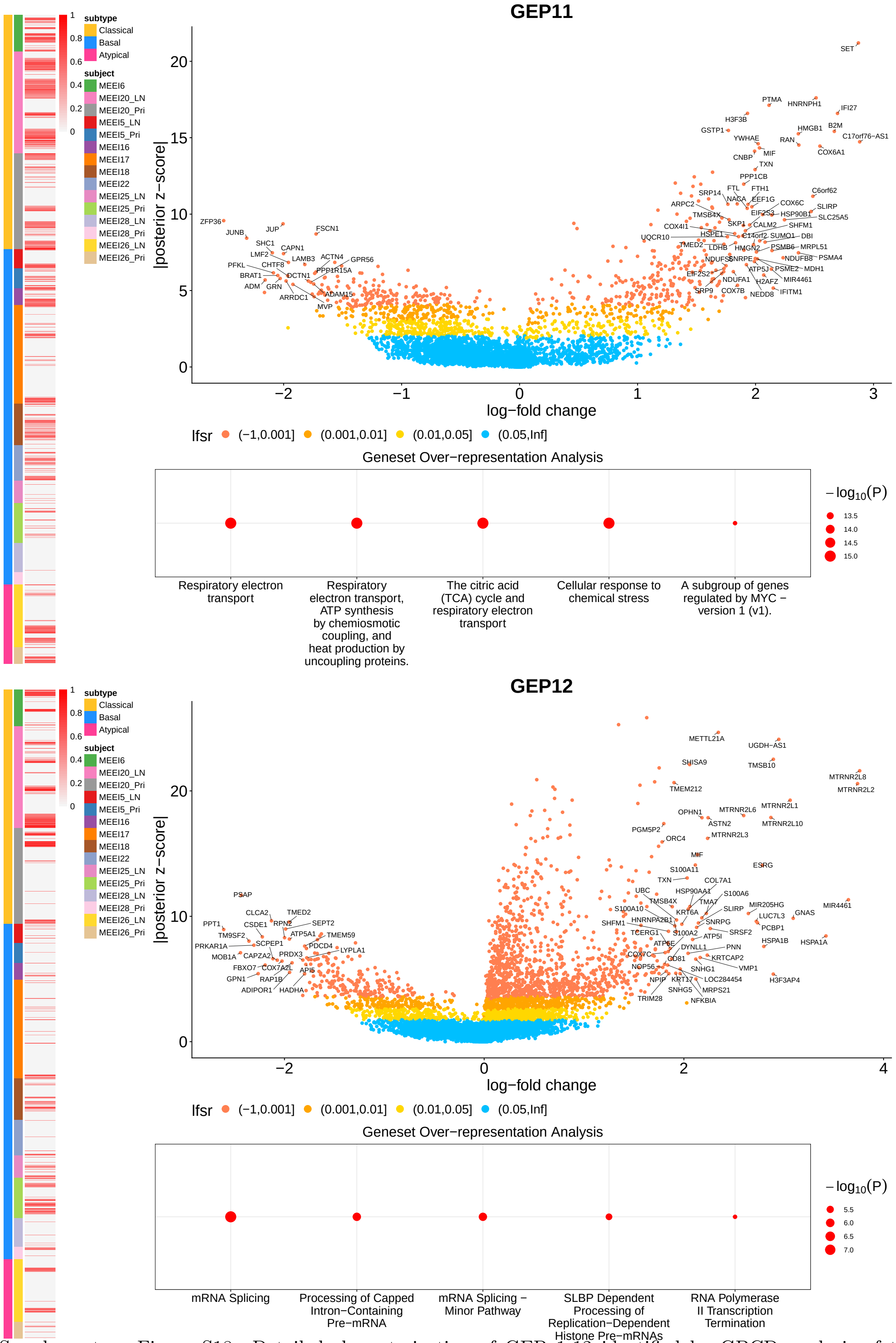


Supplementary Figure S18: Detailed characterization of GEP 1-12 identified by GBCD analysis of the HNSCC scRNA-seq dataset.

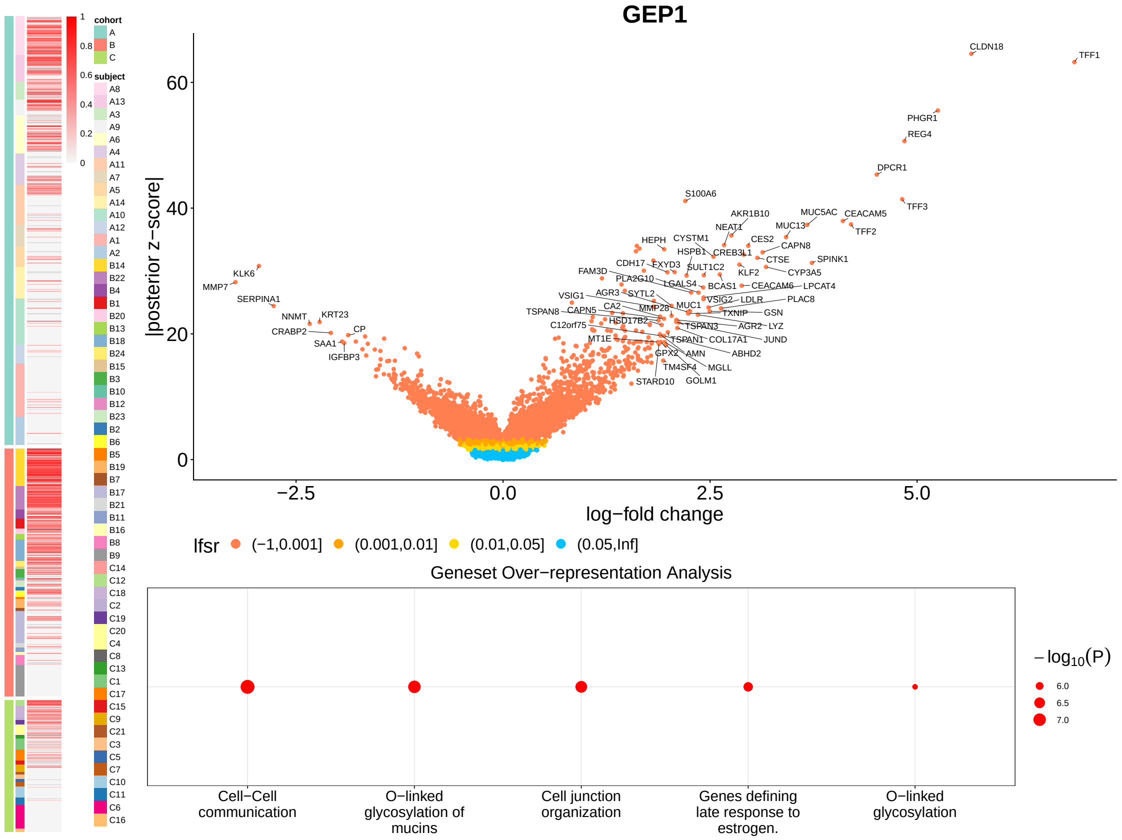
GEP1
|posterior z−score|
log−fold change
lfsr (−1,0.001] (0.001,0.01] (0.01,0.05] (0.05,Inf]
Geneset Over−representation Analysis
Cell−Cell communication
O−linked glycosylation of mucins
Cell junction organization
Genes defining late response to estrogen.
O−linked glycosylation
−log10(P)
cohort
subject

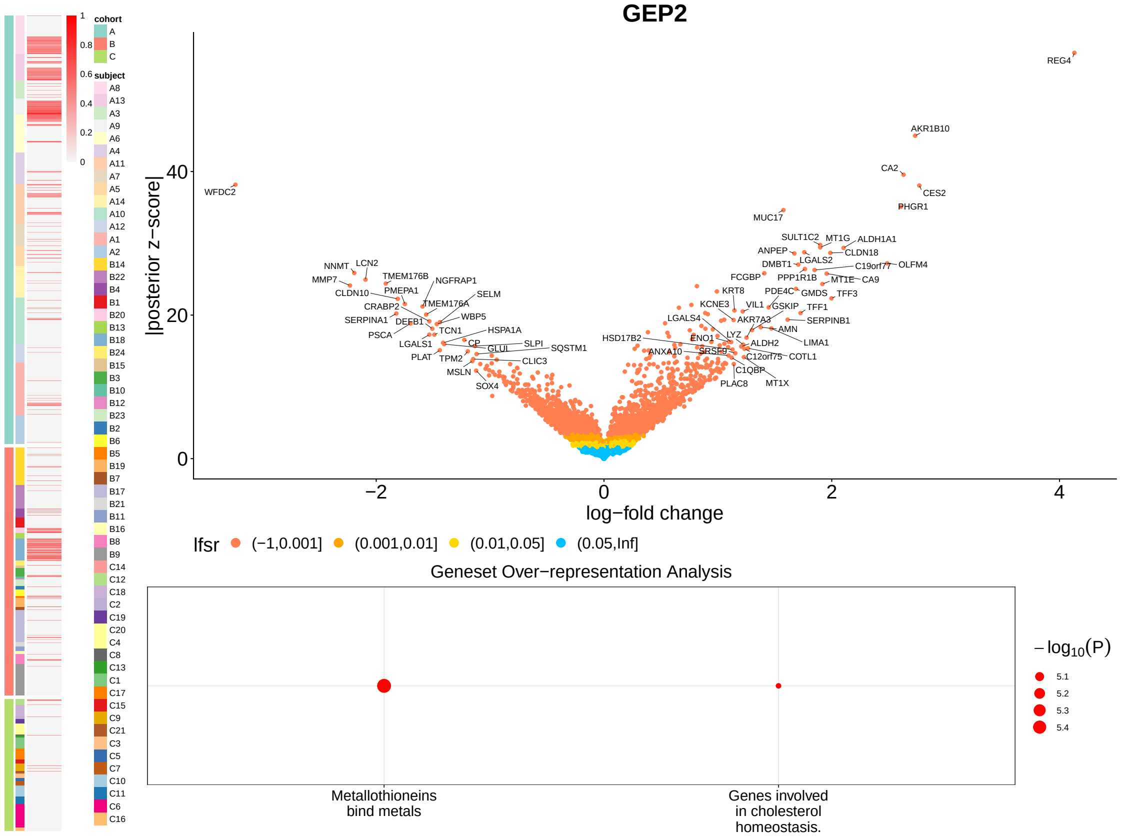
GEP2
|posterior z−score|
log−fold change
lfsr (−1,0.001] (0.001,0.01] (0.01,0.05] (0.05,Inf]
Geneset Over−representation Analysis
Metallothioneins bind metals
Genes involved in cholesterol homeostasis.
−log10(P)
cohort
subject

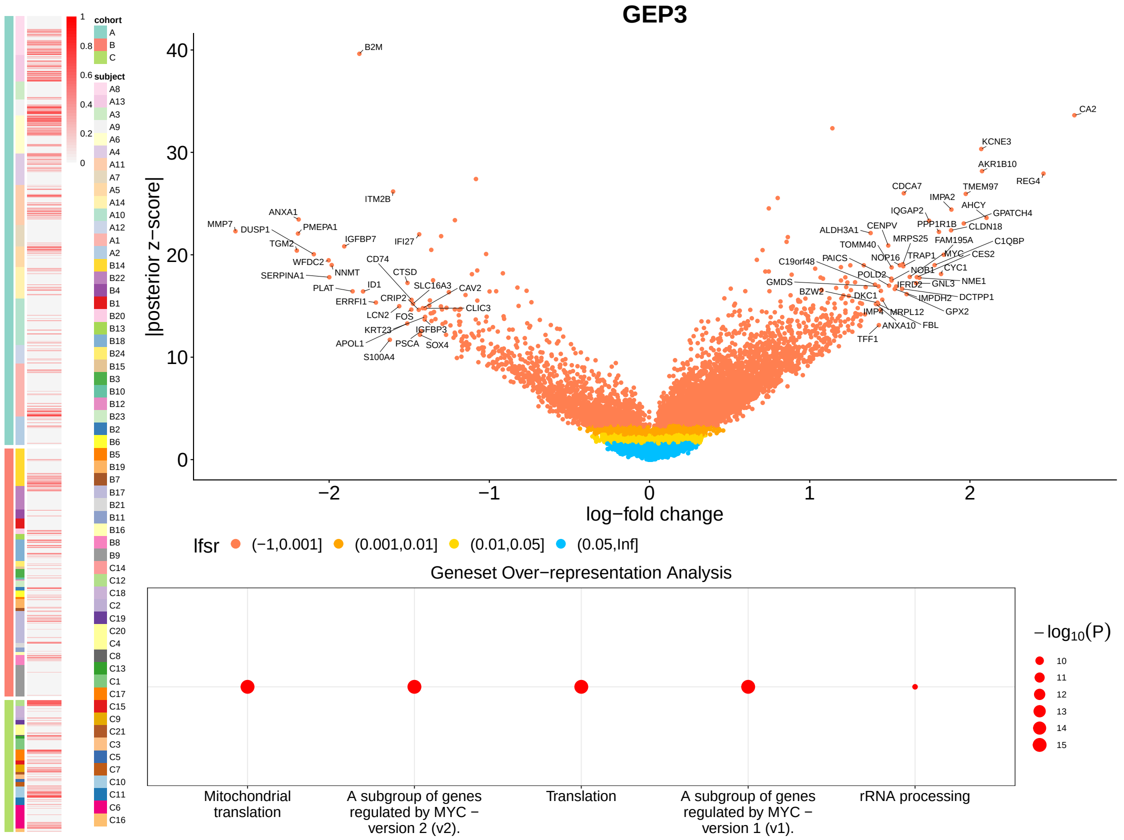
GEP3
cohort
A
B
C
subject
|posterior z−score|
log−fold change
B2M
CA2
KCNE3
AKR1B10
REG4
CDCA7
TMEM97
IMPA2
AHCY
IQGAP2
GPATCH4
ITM2B
ANXA1
MMP7
DUSP1
PMEPA1
TGM2
IGFBP7
IFI27
WFDC2
CD74
SERPINA1
NNMT
CTSD
PLAT
ID1
SLC16A3
CAV2
ERRFI1
CRIP2
LCN2
FOS
CLIC3
KRT23
IGFBP3
APOL1
PSCA
SOX4
S100A4
ALDH3A1
CENPV
PPP1R1B
CLDN18
MRPS25
FAM195A
C1QBP
TOMM40
NOP16
TRAP1
MYC
CES2
C19orf48
PAICS
NOB1
CYC1
GMDS
POLD2
IFRD2
GNL3
NME1
BZW2
DKC1
IMPDH2
DCTPP1
IMP4
MRPL12
GPX2
ANXA10
FBL
TFF1
lfsr
(−1,0.001]
(0.001,0.01]
(0.01,0.05]
(0.05,Inf]
Geneset Over−representation Analysis
Mitochondrial translation
A subgroup of genes regulated by MYC − version 2 (v2).
Translation
A subgroup of genes regulated by MYC − version 1 (v1).
rRNA processing
−log10(P)

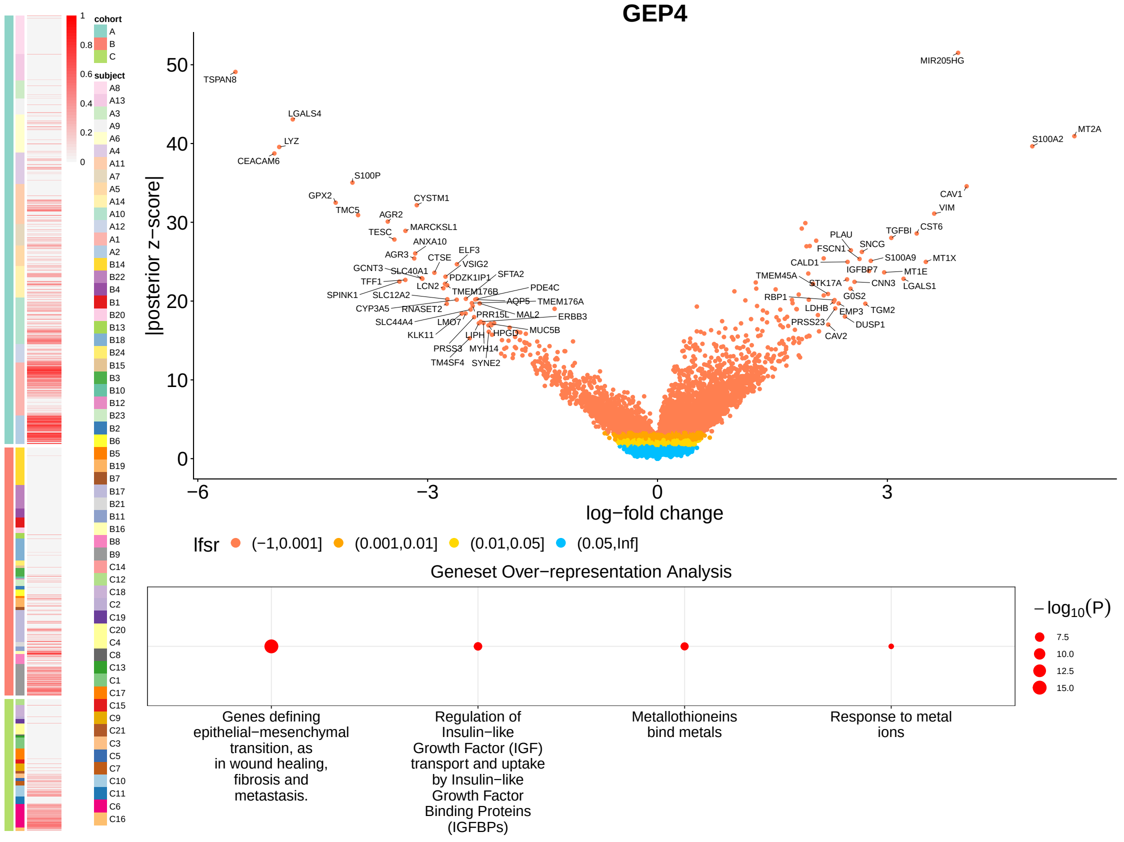
GEP4
cohort
A
B
C
subject
|posterior z−score|
log−fold change
MIR205HG
TSPAN8
LGALS4
LYZ
CEACAM6
S100A2
MT2A
S100P
GPX2
CYSTM1
TMC5
AGR2
CAV1
VIM
TESC
MARCKSL1
ANXA10
TGFBI
CST6
PLAU
FSCN1
SNCG
AGR3
CTSE
ELF3
CALD1
IGFBP7
S100A9
MT1X
GCNT3
SLC40A1
VSIG2
MT1E
TFF1
LCN2
PDZK1IP1
SFTA2
TMEM45A
STK17A
CNN3
LGALS1
SPINK1
SLC12A2
TMEM176B
PDE4C
RBP1
G0S2
CYP3A5
RNASET2
AQP5
TMEM176A
LDHB
TGM2
SLC44A4
LMO7
PRR15L
MAL2
ERBB3
EMP3
KLK11
LIPH
HPGD
MUC5B
PRSS23
DUSP1
PRSS3
MYH14
CAV2
TM4SF4
SYNE2
lfsr
(−1,0.001]
(0.001,0.01]
(0.01,0.05]
(0.05,Inf]
Geneset Over−representation Analysis
Genes defining epithelial−mesenchymal transition, as in wound healing, fibrosis and metastasis.
Regulation of Insulin−like Growth Factor (IGF) transport and uptake by Insulin−like Growth Factor Binding Proteins (IGFBPs)
Metallothioneins bind metals
Response to metal ions
−log10(P)

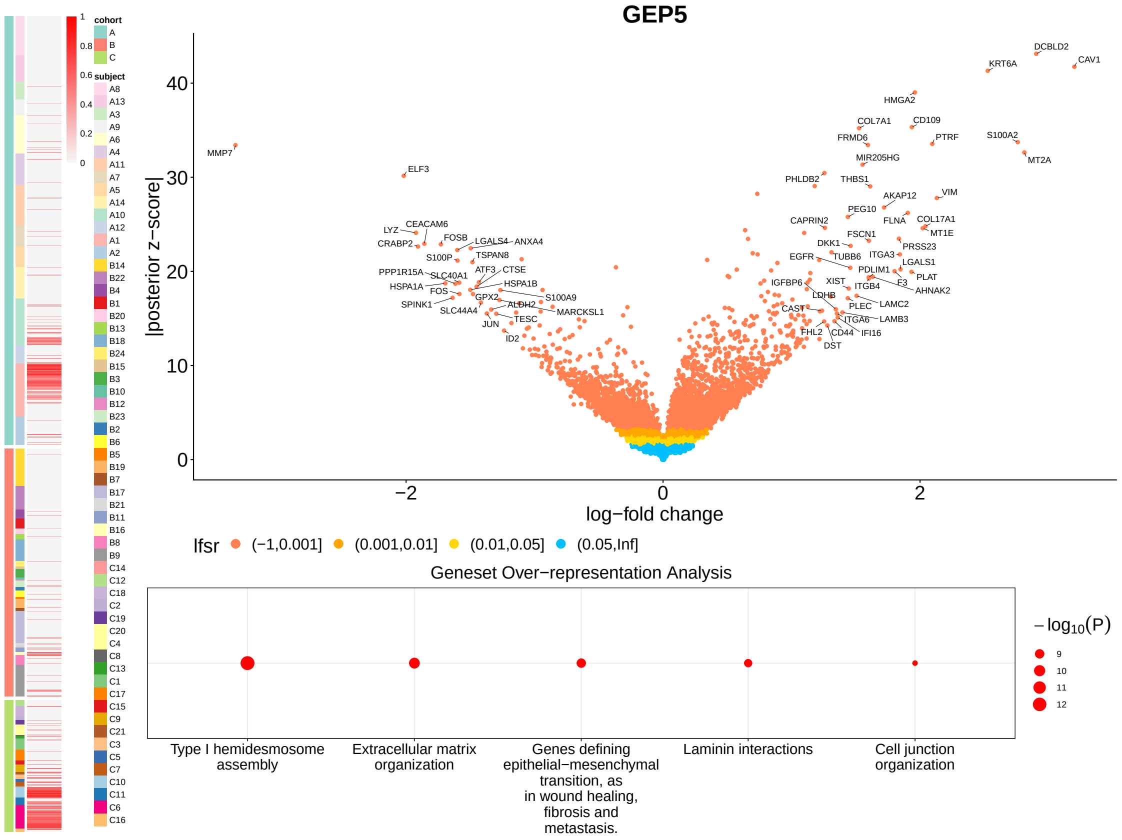
GEP5
cohort
A
B
C
subject
|posterior z−score|
log−fold change
MMP7
ELF3
LYZ
CEACAM6
CRABP2
FOSB
LGALS4
ANXA4
S100P
TSPAN8
PPP1R15A
SLC40A1
ATF3
CTSE
HSPA1A
FOS
HSPA1B
GPX2
S100A9
SPINK1
SLC44A4
ALDH2
MARCKSL1
TESC
JUN
ID2
DCBLD2
KRT6A
CAV1
HMGA2
COL7A1
CD109
FRMD6
PTRF
S100A2
MT2A
MIR205HG
PHLDB2
THBS1
VIM
AKAP12
PEG10
FLNA
COL17A1
CAPRIN2
MT1E
FSCN1
DKK1
PRSS23
EGFR
TUBB6
ITGA3
LGALS1
PDLIM1
PLAT
IGFBP6
XIST
F3
ITGB4
AHNAK2
LDHB
PLEC
LAMC2
CAST
ITGA6
LAMB3
FHL2
CD44
IFI16
DST
lfsr
(−1,0.001]
(0.001,0.01]
(0.01,0.05]
(0.05,Inf]
Geneset Over−representation Analysis
Type I hemidesmosome assembly
Extracellular matrix organization
Genes defining epithelial−mesenchymal transition, as in wound healing, fibrosis and metastasis.
Laminin interactions
Cell junction organization
−log10(P)
9
10
11
12

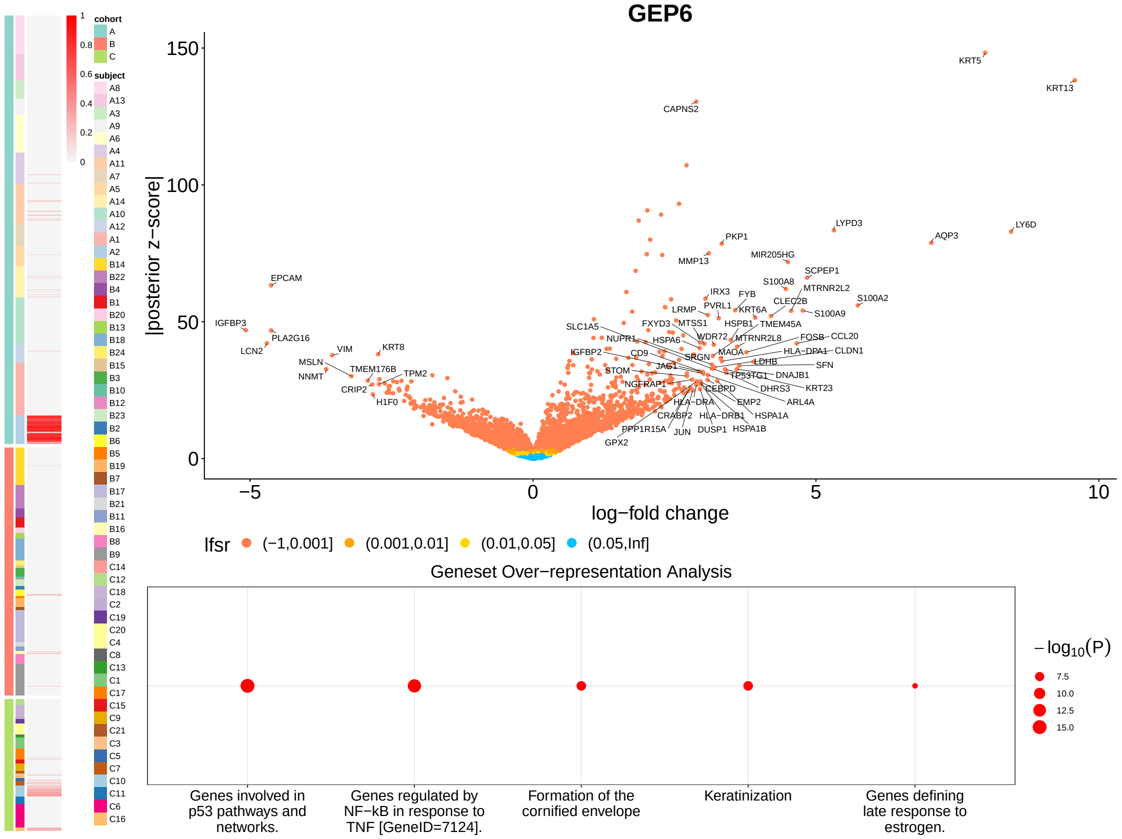
GEP6
cohort
A
B
C
subject
|posterior z−score|
log−fold change
KRT5
KRT13
CAPNS2
LYPD3
LY6D
PKP1
AQP3
MMP13
MIR205HG
SCPEP1
EPCAM
S100A8
MTRNR2L2
IRX3
FYB
PVRL1
KRT6A
CLEC2B
S100A2
LRMP
S100A9
IGFBP3
PLA2G16
SLC1A5
FXYD3
MTSS1
HSPB1
TMEM45A
NUPR1
HSPA6
WDR72
MTRNR2L8
FOSB
CCL20
LCN2
VIM
KRT8
IGFBP2
CD9
SRGN
MAOA
HLA−DPA1
CLDN1
MSLN
TMEM176B
JAG1
LDHB
SFN
NNMT
TPM2
STOM
TP53TG1
DNAJB1
CRIP2
NGFRAP1
CEBPD
DHRS3
KRT23
H1F0
HLA−DRA
EMP2
ARL4A
CRABP2
HLA−DRB1
HSPA1A
PPP1R15A
JUN
DUSP1
HSPA1B
GPX2
lfsr
(−1,0.001]
(0.001,0.01]
(0.01,0.05]
(0.05,Inf]
Geneset Over−representation Analysis
Genes involved in p53 pathways and networks.
Genes regulated by NF−kB in response to TNF [GeneID=7124].
Formation of the cornified envelope
Keratinization
Genes defining late response to estrogen.
−log10(P)
7.5
10.0
12.5
15.0

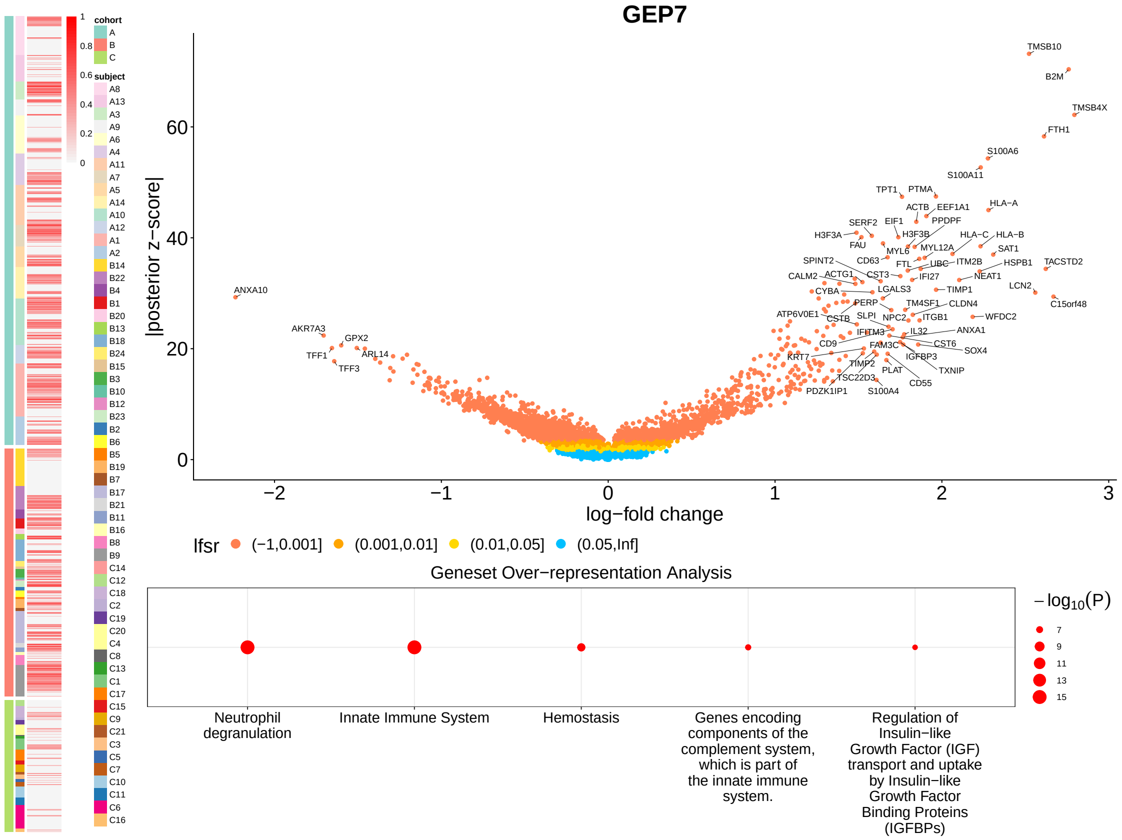
GEP7
cohort
A
B
C
subject
|posterior z−score|
log−fold change
lfsr (−1,0.001] (0.001,0.01] (0.01,0.05] (0.05,Inf]
Geneset Over−representation Analysis
−log10(P)
Neutrophil degranulation
Innate Immune System
Hemostasis
Genes encoding components of the complement system, which is part of the innate immune system.
Regulation of Insulin−like Growth Factor (IGF) transport and uptake by Insulin−like Growth Factor Binding Proteins (IGFBPs)
TMSB10
B2M
TMSB4X
FTH1
S100A6
S100A11
ANXA10
AKR7A3
GPX2
TFF1
ARL14
TFF3

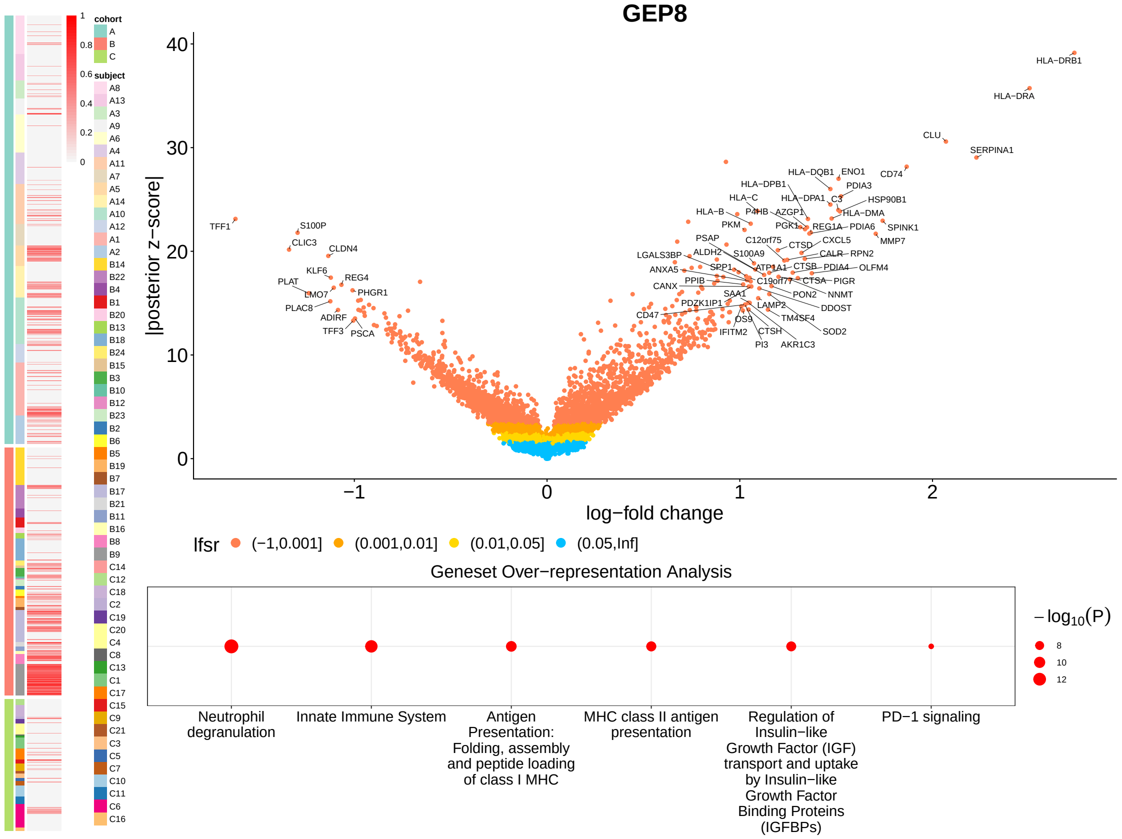
GEP8
cohort
A
B
C
subject
|posterior z−score|
log−fold change
lfsr (−1,0.001] (0.001,0.01] (0.01,0.05] (0.05,Inf]
Geneset Over−representation Analysis
−log10(P)
Neutrophil degranulation
Innate Immune System
Antigen Presentation: Folding, assembly and peptide loading of class I MHC
MHC class II antigen presentation
Regulation of Insulin−like Growth Factor (IGF) transport and uptake by Insulin−like Growth Factor Binding Proteins (IGFBPs)
PD−1 signaling
HLA−DRB1
HLA−DRA
CLU
SERPINA1
CD74
TFF1
S100P
CLDN4
CLIC3

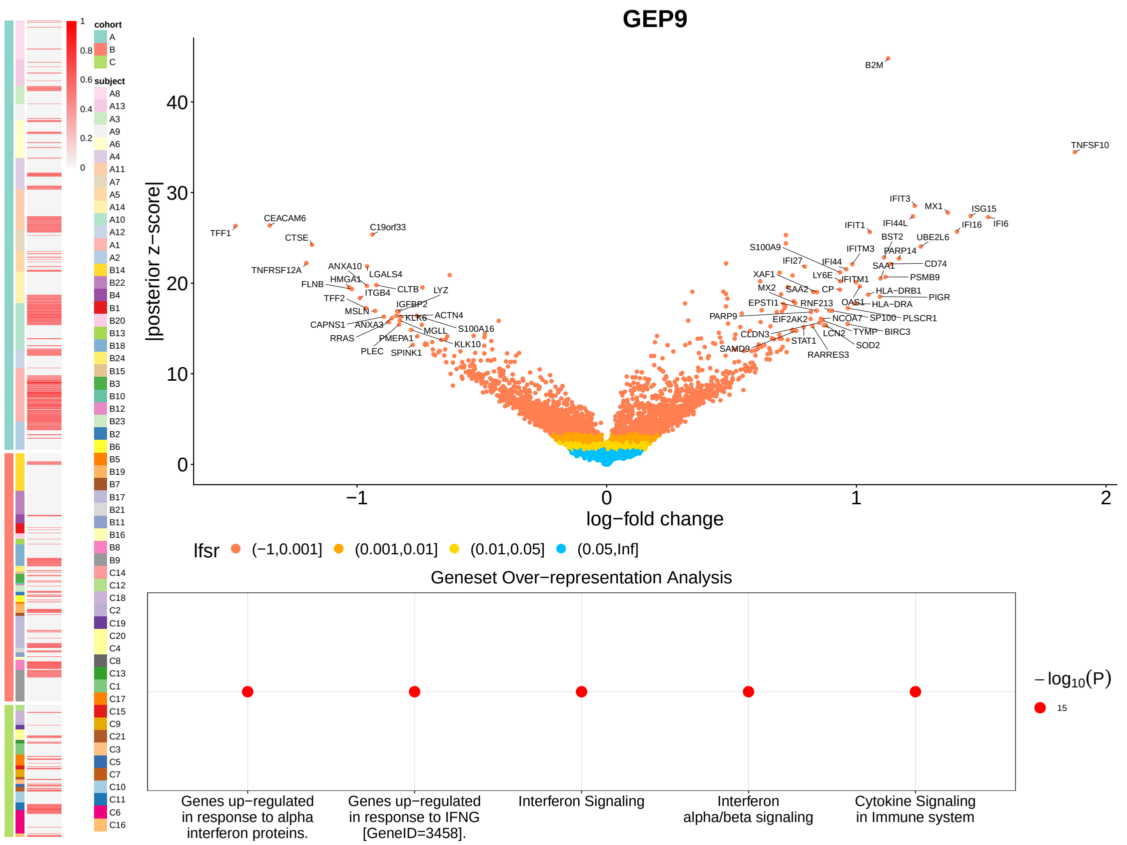
GEP9
cohort
A
B
C
subject
|posterior z−score|
log−fold change
lfsr (−1,0.001] (0.001,0.01] (0.01,0.05] (0.05,Inf]
Geneset Over−representation Analysis
Genes up−regulated in response to alpha interferon proteins.
Genes up−regulated in response to IFNG [GeneID=3458].
Interferon Signaling
Interferon alpha/beta signaling
Cytokine Signaling in Immune system
$-\log_{10}(P)$
15

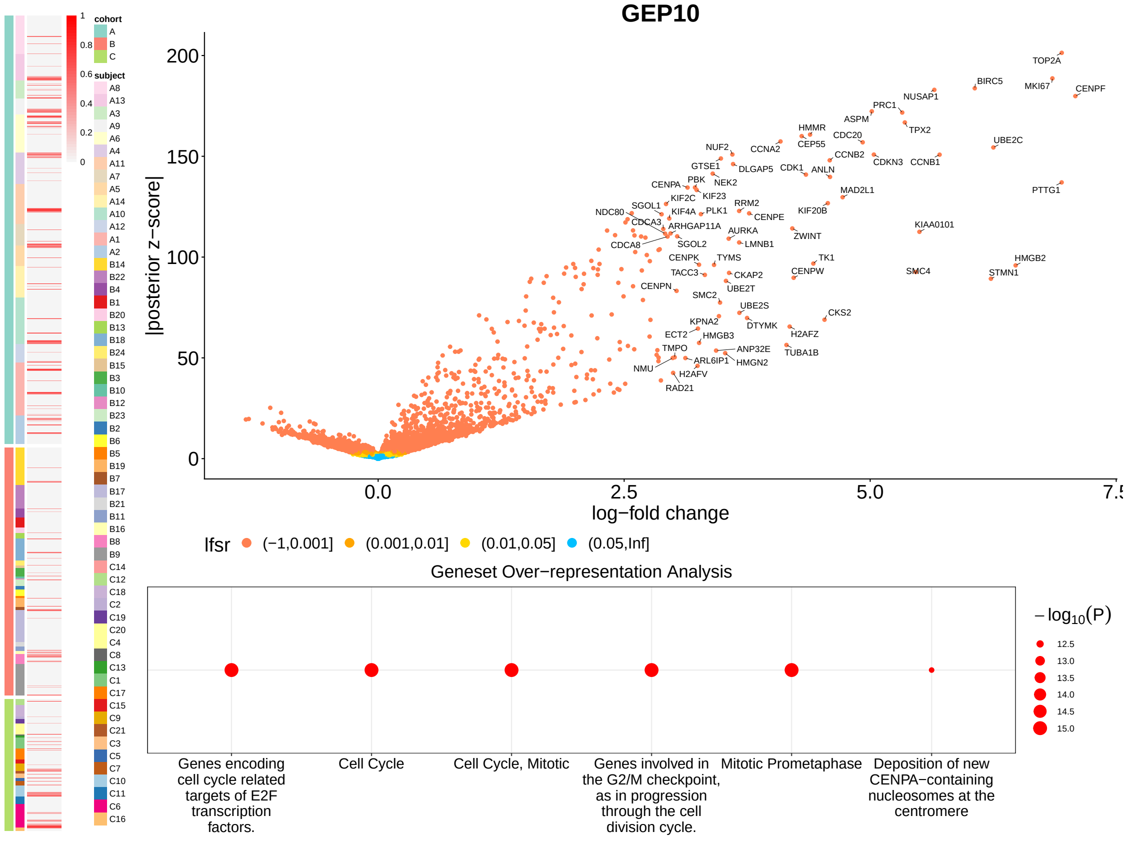
GEP10
cohort
A
B
C
subject
|posterior z−score|
log−fold change
lfsr (−1,0.001] (0.001,0.01] (0.01,0.05] (0.05,Inf]
Geneset Over−representation Analysis
Genes encoding cell cycle related targets of E2F transcription factors.
Cell Cycle
Cell Cycle, Mitotic
Genes involved in the G2/M checkpoint, as in progression through the cell division cycle.
Mitotic Prometaphase
Deposition of new CENPA−containing nucleosomes at the centromere
$-\log_{10}(P)$
12.5
13.0
13.5
14.0
14.5
15.0

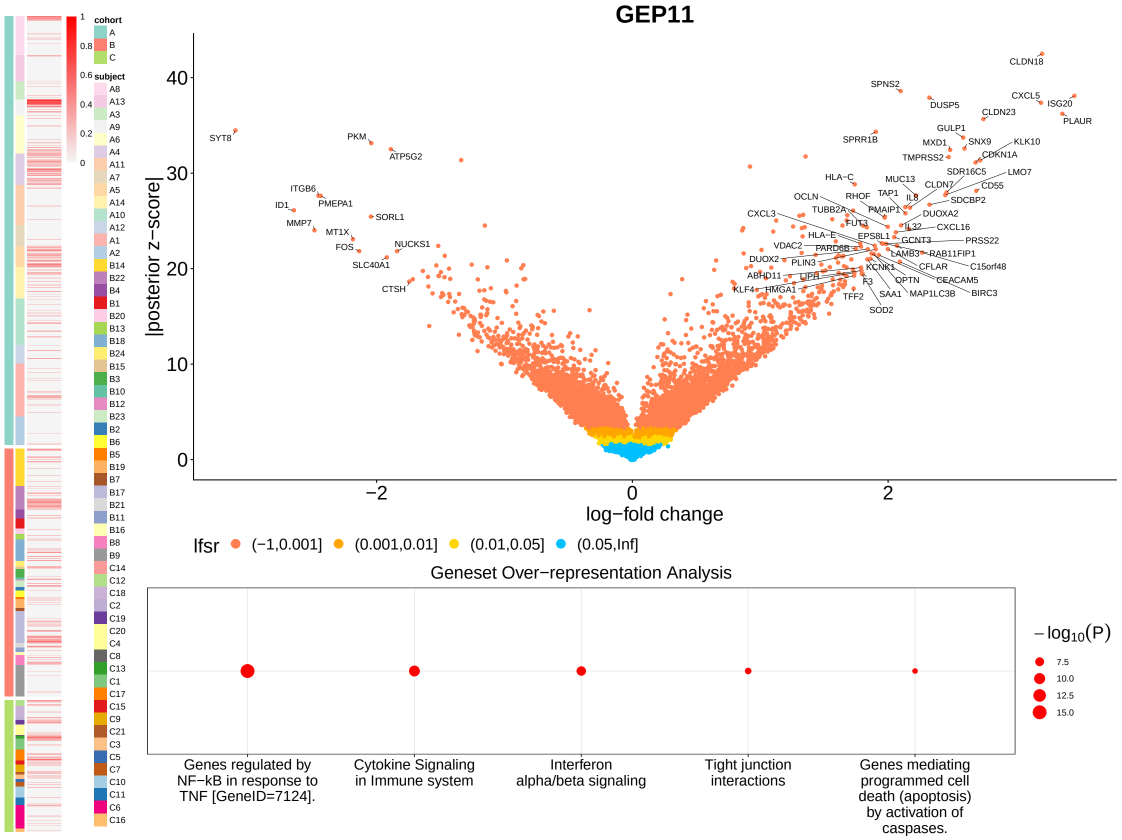
GEP11
cohort
A
B
C
subject
|posterior z−score|
log−fold change
lfsr (−1,0.001] (0.001,0.01] (0.01,0.05] (0.05,Inf]
Geneset Over−representation Analysis
Genes regulated by NF−kB in response to TNF [GeneID=7124].
Cytokine Signaling in Immune system
Interferon alpha/beta signaling
Tight junction interactions
Genes mediating programmed cell death (apoptosis) by activation of caspases.
−log10(P)
7.5
10.0
12.5
15.0

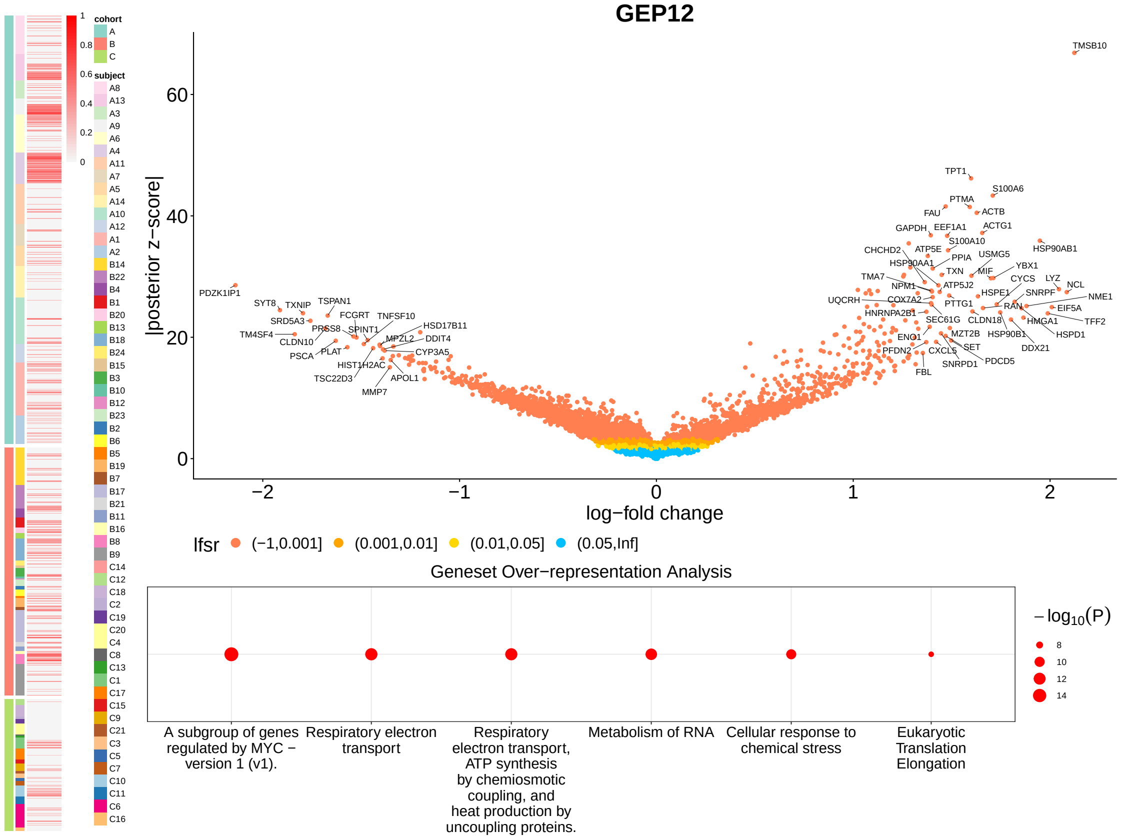
GEP12
cohort
A
B
C
subject
|posterior z−score|
log−fold change
lfsr (−1,0.001] (0.001,0.01] (0.01,0.05] (0.05,Inf]
Geneset Over−representation Analysis
A subgroup of genes regulated by MYC − version 1 (v1).
Respiratory electron transport
Respiratory electron transport, ATP synthesis by chemiosmotic coupling, and heat production by uncoupling proteins.
Metabolism of RNA
Cellular response to chemical stress
Eukaryotic Translation Elongation
−log10(P)
8
10
12
14

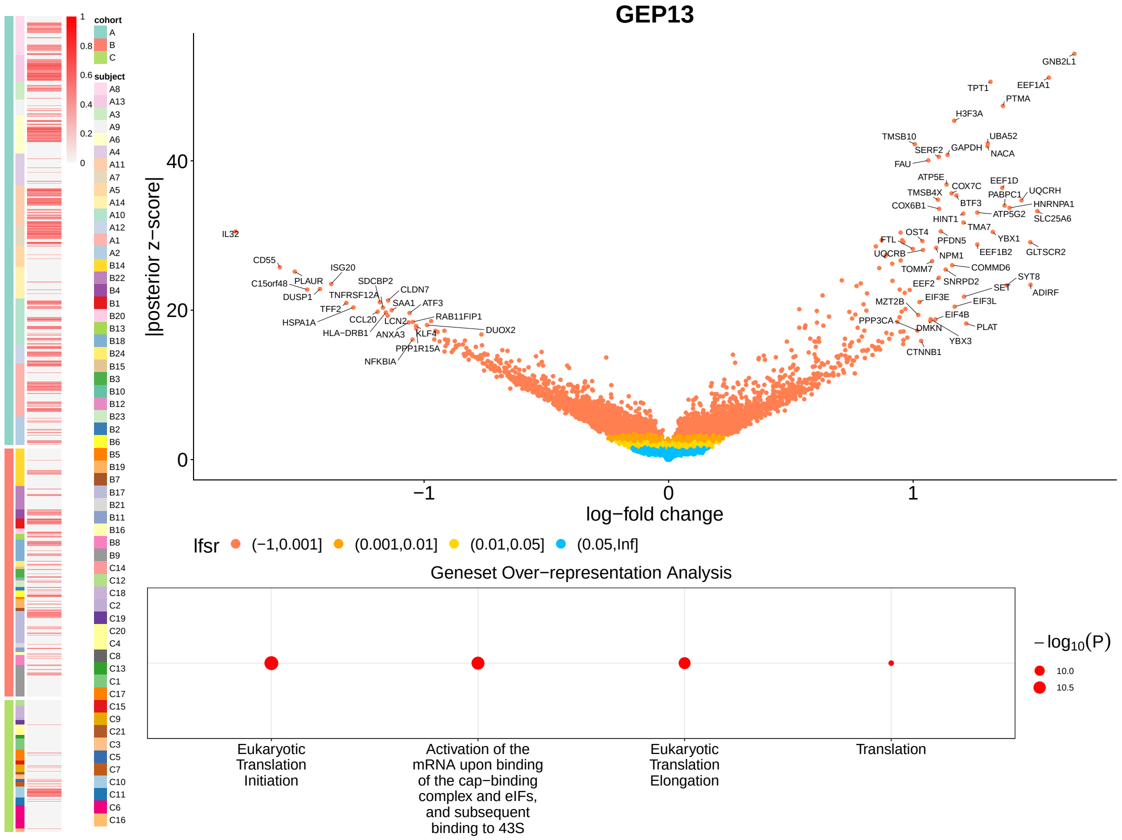


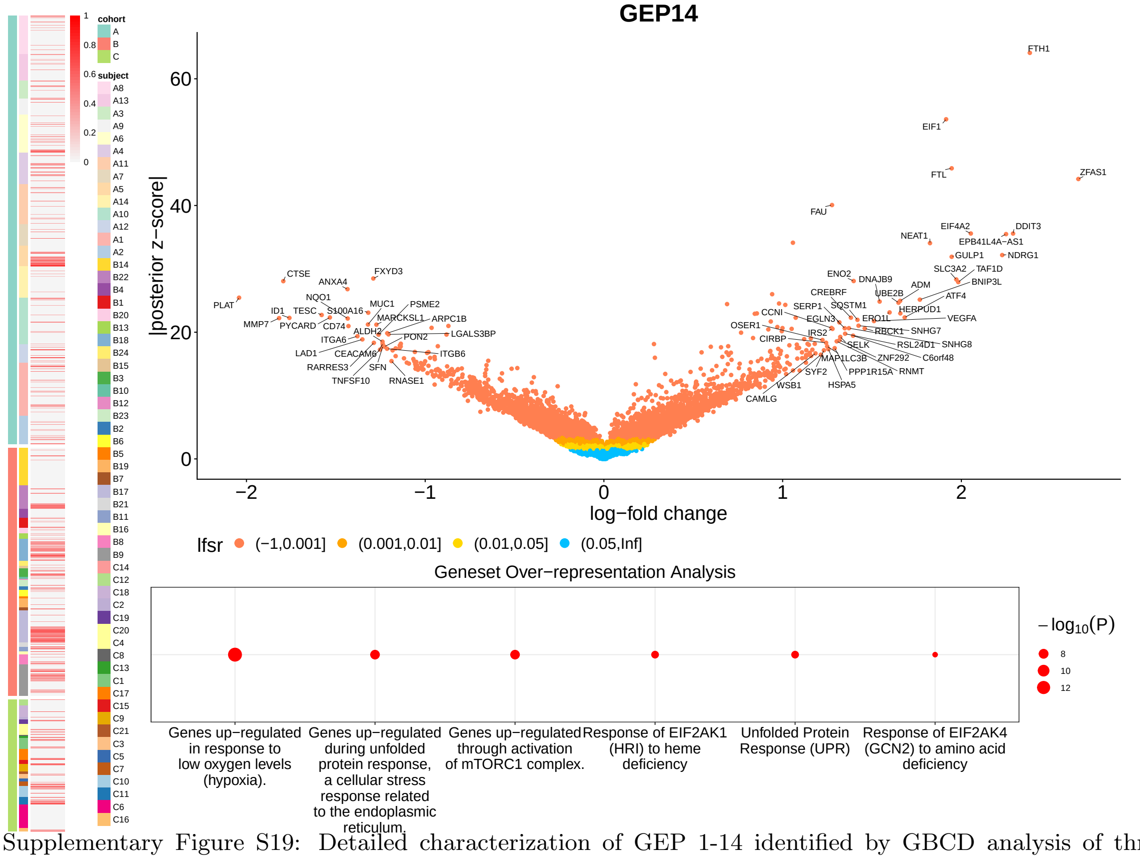


Supplementary Figure S19: Detailed characterization of GEP 1-14 identified by GBCD analysis of three PDAC scRNA-seq datasets.
